# HybPhaser: a workflow for the detection and phasing of hybrids in target capture datasets

**DOI:** 10.1101/2020.10.27.354589

**Authors:** Lars Nauheimer, Nicholas Weigner, Elizabeth Joyce, Darren Crayn, Charles Clarke, Katharina Nargar

**Author notes:** Author for correspondence: Lars Nauheimer.

## Abstract

**Premise of the study:** Hybrids contain divergent alleles that can confound phylogenetic analyses but can provide insights into reticulated evolution when identified and phased. We developed a workflow to detect hybrids in target capture datasets and phase reads into parental lineages using a similarity and phylogenetic framework.

**Methods:** We used Angiosperms353 target capture data for *Nepenthes* including known hybrids to test the novel workflow. Reference mapping was used to assess heterozygous sites across the dataset, detect hybrid accessions and paralogous genes. Hybrid samples were phased by mapping reads to multiple references and sorting reads according to similarity. Phased accessions were included in the phylogenetic framework.

**Results:** All known *Nepenthes* hybrids and nine more samples had high levels of heterozygous sites, reads associated with multiple divergent clades, and were phased into accessions resembling divergent haplotypes. Phylogenetic analysis including phased accessions increased clade support and confirmed parental lineages of hybrids.

**Discussion:** HybPhaser provides a novel approach to detect and phase hybrids in target capture datasets, which can provide insights into reticulations by revealing origins of hybrids and reduce conflicting signal leading to more robust phylogenetic analyses.

## Introduction

Reticulation events caused by hybridization are common and important sources of novelty in angiosperm evolution (Wood et al. 2009, Palfalvi et al. 2020). The detection, investigation and representation of hybridization remains a challenge in phylogenomics (Kellog 2016, Mallet et al. 2016, Spooner et al. 2020). The combination of divergent genomes in hybrids (herein used for any organism that contains divergent genomes due to a hybridization event, e.g. polyploids) introduces conflicting phylogenetic signal and can lead to topologically incorrect or poorly resolved phylogenetic trees (McDade 1992, Soltis et al. 2008). However, the advancement of target-capture data and universal probe-kits such as Angiosperms353 (Johnson et al. 2018) provides an opportunity to gain insight into historical reticulations in angiosperm evolution and reduce phylogenetic conflict, if ortholog (or homeolog in polyploids) gene variants can be identified and separated (phased).

Previously, inclusion of phased gene variants in phylogenetic studies has been used to confirm hybrid status of organisms (Sang and Zhang 1999), the origin of polyploids (Popp and Oxelman 2001), reveal parental lineages (Tripplet et al. 2012, Estep et al. 2014), enable reconstruction of past reticulations (Estep et al. 2014, Brassec and Blattner 2015), and date ancient hybridization events (Marcussen et al. 2015). Using Sanger sequencing, single-gene studies generated sequences for each variant separately using cloning (Sang and Zhang 1999, Popp and Oxelman 2001) and multi-gene studies linked these gene variants using their phylogenetic association in single gene phylogenies (Tripplet et al. 2012, Estep et al. 2014, Marcussen et al. 2015). However, linking phased gene variants was limited by the resolution of single locus phylogenies to detect parental clades and the low numbers of nuclear genes available using Sanger sequencing to generate robust datasets. The availability of universal target capture sequencing bait-kits such as Angiosperms353 (Johnson et al. 2018) has revolutionized the study of angiosperm evolution by enabling the study of evolution across a phylogenetically broad group and potentially recovers sequence reads from all variants for hundreds of nuclear genes. Two challenges remain, assembling reads into phased sequences from each gene variant and linking genes according to their origin to investigate phased accessions.

*De novo* assembly, as implemented in common pipelines such as PHYLUCE (Faircloth 2016), HybPiper (Johnson et al. 2016) and SECAPR (Andermann et al. 2018) can only recover complete phased gene variants when all polymorphic sites are frequent enough to connect all sequence reads across the whole gene locus (Fig. 1A). If this is not the case, sequences from different gene variants (including paralogs) can be unwittingly combined to generate chimeric contigs (Kates et al. 2018). Some approaches utilize paired-end reads to increase the connectivity of polymorphisms across reads (Andermann et al. 2019); however, phased blocks do not always span across all gene loci (Kates et al. 2018). Even if all gene variants are perfectly phased, the problem of linking phased sequences across gene loci and unconnected exons to generate phased haplotypes persists.

**Figure 1.**
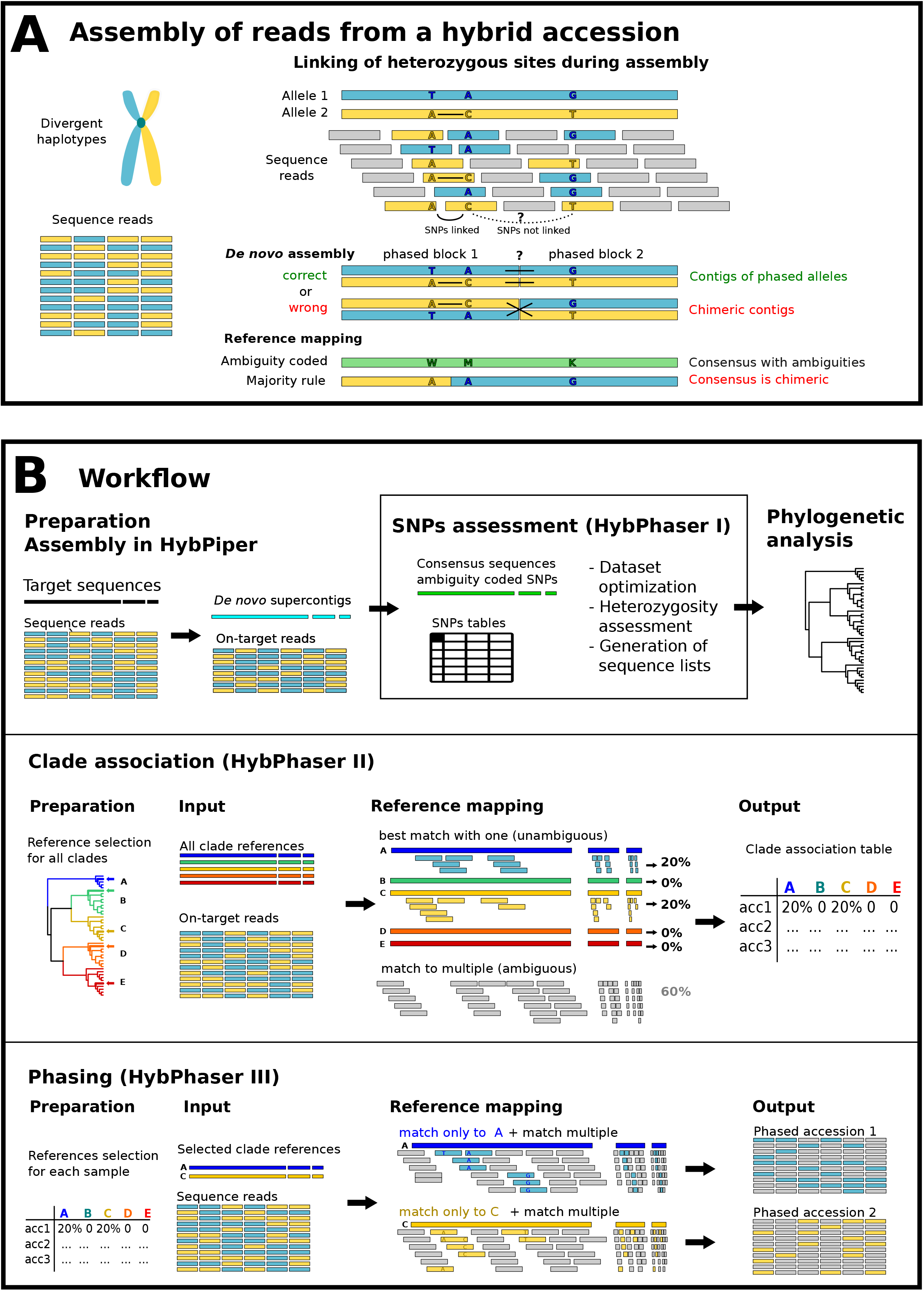
A) Illustration of the linkage of heterozygous sites in the assembly of hybrid accessions that contain reads from two divergent haplotypes (blue-TAG and yellow-ACT). The two SNPs on the left can be linked (continuous line), but they cannot be linked to the third SNP on the right (dotted line). Reference mapping can result in consensus sequence coding SNPs as ambiguities (MWK) or represent the most common nucleotide (AAG) generating a chimeric sequence. *De novo* assembly can either connect phased blocks correctly into phased alleles (TAG/ATC) or generate chimeric sequences (ACG/TAT). B) Illustration of major parts and concepts of the workflow. 1) Assembly in HybPiper, 2) SNPs assessment in HybPhaser, 3) Phylogenetic analysis, 4) Clade association and 5) Phas-ing.

An alternative method for sequence assembly is reference mapping, where reads are mapped to a single reference (as implemented in pipelines such as HybPhyloMaker (Fer & Schmickl 2018). Reads from all gene variants are mapped, and the presence of heterozygous sites generated by multiple haplotypes can be handled in two ways (Fig. 1A); the nucleotide is called using a majority rule (as in HybPhyloMaker) or an IUPAC ambiguity code is used to accommodate divergent nucleotides and prevent the creation of chimeric sequences (Uribe-Convers et al. 2016, Kates et al. 2018). However, phasing of hybrid accessions is not possible.

Here we present HybPhaser, a bioinformatic workflow that can phase hybrid accessions by mapping sequence reads simultaneously to references from the parental clades and sorting reads accordingly to generate accessions that approximate phased haplotypes. This approach performs the phasing prior to the assembly and thus avoids difficulties with linking of heterozygous sites and gene loci but requires a phylogenetic framework and method to select suitable references.

HybPhaser is built as an extension to the *de novo* assembly of HybPiper and consists of three parts (Fig. 1B): (1) assessment of heterozygous sites in assembled sequences to detect putative hybrid accessions; (2) creation of a read-to-clade association framework; and (3) the phasing of read files based on the clade association framework. It extends the HybPiper pipeline further by enabling the modulation of paralog detection based on the heterozygosity of the dataset, facilitating the optimization of the dataset by cleaning samples and loci with poor recovery, and enabling the collation of consensus sequences (with ambiguity codes) as well as *de novo* contigs into sequence lists for subsequent analyses.

Herein we detail the HybPhaser workflow and demonstrate its utility for detecting hybrids and identifying parental lineages using Angiosperms353 data from the carnivorous plant genus *Nepenthes* L. (Nepenthaceae).

## Methods

### Preparation of the dataset and assembly

#### Study group

*Nepenthes* is a palaeotropical genus of c. 160 species known to hybridize readily in horticulture and nature (Clarke et al. 2018). Molecular phylogenetic studies of the genus based on Sanger-sequenced loci have resulted in poorly resolved and supported trees due to limited resolution of the markers, incongruence between plastid and nuclear phylogenies, and the possible inclusion of ITS paralogs (Meimberg and Heubl 2006, Alamsyah and Ito 2013). Recently, phylogenomic approaches including genome skimming (Nauheimer et al. 2019) and Angiosperms353 target capture (Murphy et al. 2020) have provided great advances in resolving the *Nepenthes* phylogeny but were unable to clarify past reticulations evident in the datasets. In a study that sampled almost all recognized species in the genus, Murphy et al. (2020), included two known hybrids and inferred one putative hybrid accession from conflict between supermatrix and gene tree analyses. Given the availability of suitable data with good species coverage and presence of known natural and cultivated hybrids, *Nepenthes* is an suitable model group for demonstrating hybrid detection and phasing using HybPhaser and the Angiosperms353 probe set.

#### Input data

We generated Angiosperms353 target capture data for 125 samples of 68 *Nepenthes* species, including 15 samples of 12 different horticultural hybrids. In addition, we downloaded the Murphy et al. (2020) *Nepenthes* raw read files from the NCBI sequence read archive (Bioproject PRJEB35235, 185 accessions including one known and one suspected hybrid) and added them to the dataset to increase species coverage across the genus. In total, 310 accessions representing 157 *Nepenthes* taxa including 17 accessions of 14 known hybrid taxa were analyzed (Appendix 2).

#### DNA extraction, library enrichment and sequencing

Total genomic DNA was isolated from leaf material stored in silica gel for 125 *Nepenthes* samples. DNA was extracted using the CTAB method (Doyle & Doyle 1987) and DNA extracts were further processed at the Australian Genomic Research Facility (AGRF; Melbourne, Australia) for library preparation using NEBNext Ultra II DNA protocol per manufacturer’s instructions as well as sequencing. Target sequence capture was carried out using the ‘Angiosperms 353 v1’ universal probe-set (ArborBioscience, Ann Arbor, USA) following Johnson et al. (2018). Sequencing was performed on an Illumina HiSeq 2500 sequencer producing 150bp paired-end reads.

#### Read trimming

The sequence reads were trimmed using Trimmomatic (v.0.39, Bolger et al. 2014) to remove sequencing adapters and poor-quality bases (illuminaclip 2:30:10, leading 20, trailing 20, sliding window 4:20). Final reads of less than 30 nucleotides were excluded.

#### HybPiper sequence assembly

*De novo* sequence assembly was performed with HybPiper (v. 1.3.1) using the Angiosperm353 target file (https://github.com/mossmatters/Angiosperms353) in order to efficiently generate gene sequences consisting of concatenated exons and extract reads matching to each gene. HybPiper performs reference mapping to pre-select reads matching each gene for an efficient *de novo* assembly of relevant reads only. If genes consist of multiple exons, assembled contigs are mapped onto the target gene and concatenated to generate a complete gene sequence. We used the nucleotide format and BWA (Li and Durbin 2009) for mapping, but the workflow can also handle amino acid sequences and BLASTX mapping. The script ‘intronerate.py’ was run to retrieve gene sequences that included intron regions. Summary statistics were generated using the script ‘hybpiper_stats.py’ (Appendix 3).

### HybPhaser part I: SNPs assessment

Reference mapping of reads from divergent gene variants lead to heterozygous sites or single nucleotide polymorphisms (SNPs) where gene variants diverge (Fig. 1A). Assessing the distribution of SNPs across samples and loci can provide insights into the occurrence of hybrid accessions and paralogous genes, both of which are expected to have considerable divergence between gene versions. HybPhaser generates consensus sequences through reference mapping and codes SNPs as IUPAC ambiguity characters, which can be recorded, and quantified using R scripts and assessed in generated graphs and tables.

#### Generation of consensus sequences

HybPhaser relies on the output of HybPiper assembly for remapping and consensus sequence generation using the bash script ‘Generate_consensus_sequences.sh’. For each sample and gene, the HybPiper *de novo* contig was used as reference to which the pre-selected on-target reads were mapped using BWA generating a BAM file. Consensus sequence files containing ambiguity codes at SNP sites were generated using bcftools (v1.9, http://samtools.github.io/bcftools/bcftools.html). To prevent sequencing artefacts being coded as SNPs, variants are only called as SNPs if the read depth is ≥10, the alternative nucleotide count is ≥4, and the proportion of alternative nucleotides is ≥0.15. The script ‘Rscript_1a_count_snps_in_consensus_seqs.R’ was then used to collate information on the proportions of SNPs and length of sequences from all samples and loci (Appendix 4).

#### Reducing proportions of missing data

Missing data can have negative consequences for downstream analyses (e.g., for gene tree summary analyses; Mirarab (2019)), and low sequence recovery can indicate poor assembly quality. The HybPhaser script ‘R1b_optimize_dataset.R’ can be used to explore loci and sample quality, and to exclude loci with poor sequence recovery. Thresholds for minimum proportions of loci recovered per sample and samples recovered per locus, as well as minimum proportions of target sequence length can be adjusted in the configuration script ‘Configure_1_SNPs_assessment.R’. An initial run without dataset optimization resulted in phylogenies with poorly placed low-quality samples. To improve the quality of the dataset while keeping important samples, we set thresholds to remove all loci that had sequences recovered for less than 20% of samples or less than 25% of the target sequence length recovered and all samples that had less than 20% of loci or less than 45% of the target sequence length recovered.

#### Removal of putative paralogous genes

Genes that had been subject to gene or genome duplications can be retained as paralogs leading to the sequencing of multiple gene variants that are potentially assembled into chimeric gene sequences (Morales-Briones et al. 2018) or to high proportions of SNPs in reference mapping (Andermann et al. 2020).

The assessment of heterozygosity in HybPhaser provides a method to detect putative paralogous genes because they are expected to have higher values compared to non-paralogous genes. HybPhaser flags genes as putative paralogs that have an unusually high proportion of SNPs compared to other genes, first as average across all samples to detect paralogs shared across many samples, and second in each sample individually to detect paralogs that might not be shared with other samples. HybPhaser generates a graph and boxplot to visualise SNP distribution, and manually set a threshold for removing these putative paralogs for all samples. Alternatively, one can choose to automatically remove samples and loci with SNP frequencies that are statistical outliers (here defined as more than 1.5x the interquartile range (IQR) above the 3^rd^ quartile). Here we used the statistical outlier method to flag loci with a frequency of SNPs greater than 1.5x the interquartile range as paralogs in the script ‘Configure_1_SNPs_assessment.R’ (Appendix 6). This was done across the whole *Nepenthes* dataset, to identify and flag loci that are paralogous in all *Nepenthes* samples, as well as within each sample, so that paralogous loci unique to each sample could also be identified.

#### Assessment of locus heterozygosity and allele divergence

Hybrids inherit divergent alleles from their parental species; therefore, hybrid samples are expected to have a high proportion of loci that contain SNPs (here called locus heterozygosity or LH) and a high proportion of SNPs across all loci (here called allele divergence or AD), which should correlate to the divergence of the parental lineages. Using the script ‘Rscript_1c_summary_table.R’, we generated tables and figures that summarized the locus heterozygosity and allele divergence of each sample to identify putative hybrid *Nepenthes* samples (Fig. 2A, Appendix 7). Finally, the script ‘Rscript_1d_generate_sequence_lists.R’ was used to collate sequence lists for downstream analyses. The script collates all loci for each sample or all samples for each loci, either HybPiper *de novo* contigs or HybPhaser consensus sequences, and either with dataset optimization (paralogs and poor-quality data removed) or without.

**Figure 2.**
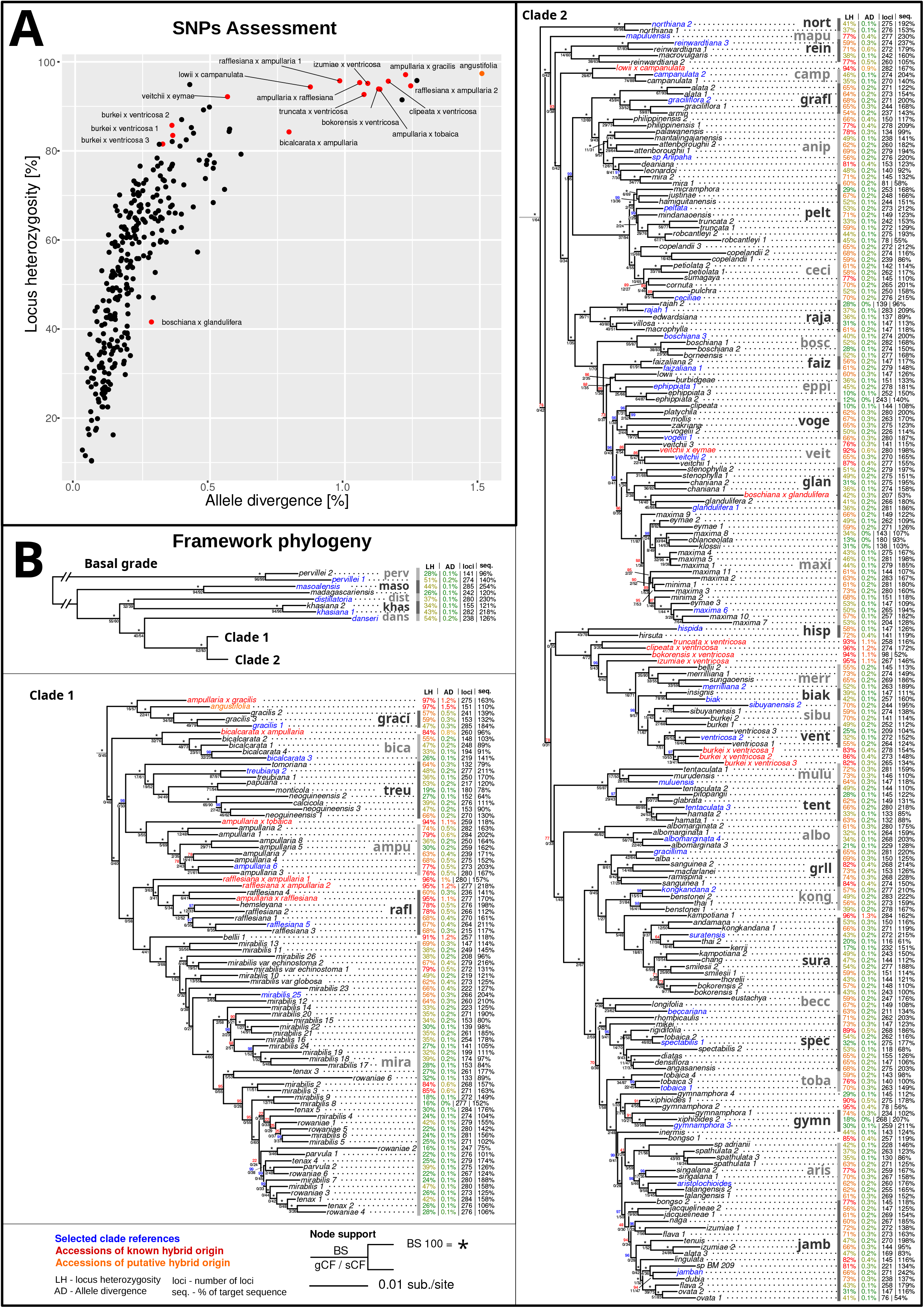
A) Scatterplot displaying the locus heterozygosity and allele divergence of samples. Known hybrids (red dots) and putative hybrid (orange dot) are labeled. B) Phylogenetic tree of consensus supermatrix displayed in three parts, basal grade with two diverging clades, clade 1 below, and clade 2 on the right. Summary statistics for each accession is given, locus heterozygosity (LH), allele divergence (AD), number of loci (loci), and proportion of target sequence recovered (seq.). Clades selected for clade association are shown in grey with bars. Clade references are displayed in blue, known hybrids in red and putative hybrid in orange. Node support is shown above the node in bootstrap (BS) (*=BS100), and below the node in gene and site concordance factors (gCF/sCF).

#### Framework phylogeny

The sequence lists produced by HybPhaser can be used for alignment and further phylogenetic analyses. Here, we first inferred phylogenetic relationships in *Nepenthes* based on unphased accessions to act as a framework for phasing later in the HybPhaser workflow. To do this, we used the consensus sequences for each locus after reducing missing data and removing putative paralogs. We preferred to use consensus sequences over contigs, as they mask heterozygous sites with ambiguity codes and therefore reduce conflicting phylogenetic signal from any hybrids. We aligned the consensus sequences using MAFFT (version 7.467, Ktoah and Standley 2013) and removed columns with more than 50% gaps using TrimAl (version 1.4.rev22, Capella-Gutierrez et al. 2009). Phylogenetic analyses were performed using maximum likelihood in the IQ-TREE2 package (version 2.0.5, Minh et al. 2020). A supermatrix phylogeny was created by concatenating all locus alignments and using the ‘edge proportional’ partition model with single loci as partitions. Phylogenetic clade support was estimated using ultra-fast bootstrap with 1000 replicates (Hoang et al. 2018) as well as gene and site concordance factors (gCF, sCF; Minh et al. 2020), which indicate the concordance of single gene trees and of 1000 randomly chosen parsimony informative sites with the phylogeny of the concatenated dataset. To determine gCF, single gene phylogenies were generated using IQ-TREE2 applying the best-fit model for each locus selected by ModelFinder (Kalyaanamoorthy et al. 2017). The supermatrix phylogeny was rooted using *Nepenthes pervillei* Blume, the sister to all other *Nepenthes* species (Murphy et al. 2020).

### HybPhaser part II: Clade association

To select suitable references for phasing, the association of sequence reads with divergent clades must be established. Hybrid accessions contain reads from divergent parental clades, which can be detected by mapping sequence reads to multiple references simultaneously. If the references are chosen carefully, reads from hybrid accessions will associate with parental clades. In HybPhaser part II, the software BBSplit (BBMap v38.47, Bushnell B. – sourceforge.net/projects/bbmap/) is used to maps reads simultaneously to multiple references and record the proportion of reads matching to references from clades across the phylogeny (Fig. 1B). The reads of samples with non-hybrid origin will only map to one clade reference; however, the reads of hybrid samples will map to multiple clades, reflecting the divergent origins of their alleles. This establishes the read to clade association required for the phasing in part III.

#### Reference selection

The selection of clade references from the framework phylogeny is critical to the efficacy of the clade association step and should be carefully considered. Clade references should be evenly distributed across the phylogeny and have low locus heterozygosity as well as low allele divergence but high coverage of target sequences. References that are too closely related will result in many ambiguous matches, while references that are too distantly related can lead to reads not matching either reference. Determining the optimal number of clade references might require multiple iterations of clade association. Here, we selected 44 clade references from the framework phylogeny (Fig. 2A), which enabled unambiguous association of reads from hybrid samples with both closely related and more distantly related parental lineages.

#### Clade association

Sequence reads files vary in the proportion of reads that matched to the target sequences. To only take into account on-target reads, we extracted those reads from the BAM file generated by HybPiper using the bash script ‘extract_mapped_reads.sh’. The software BBSplit (BBMap v38.47) was then used to map each on-target read file simultaneously to all 44 selected clade references, recording the proportions of reads that mapped to either reference unambiguously. The script ‘Rscript_2a_prepare_BBSplit_script.R’ was used to generate a BBSplit commands file with the selected clade reference names and abbreviations. The mapping results were summarized using the script ‘Rscript_2b_collate_BBSplit_results.R’ (Appendix 8).

### HybPhaser part III: Phasing

#### Phasing of sequence reads

Once sample reads have been mapped to the clade references and hybrid samples as well as their parental clades have been identified, HybPhaser uses BBSplit to phase the reads of the hybrid samples. In this step, BBSplit distributes reads that match unambiguously to one clade references into the respective read file and writes read files that match similarly well onto multiple references into all read files for that sample (Fig. 1B). Therefore, the resulting phased accessions differ only in the reads that contain sites in which the references diverge. Ideally, hybrids containing one haplotype from each parent will be phased into two accessions representing both haplotypes. Similarly, polyploids containing multiple haplotypes can be phased into multiple accessions, in which case the proportions of reads can indicate the proportions of haplotypes.

For phasing, we considered samples with multiple clade association as well as a high LH (>80%). While there are samples with multiple clade associations that have low LH, they are unlikely to be hybrids that can be unambiguously phased. However, we make one exception, the known hybrid *Nepenthes boschiana* Korth. × *glandulifera* Chi.C.Lee, which had low LH (41.6%) likely due to low sequence recovery impacting variant calling. In total, we selected 26 samples for phasing, which included all 17 accessions of known hybrids and nine additional samples that had not previously been identified as hybrids.

The R script ‘Rscript_3a_prepare_phasing_script.R’ was used to generate a bash script to run all 26 BBSplit commands that mapped the original sequence read files to the selected references for each chosen sample. Finally, the script ‘Rscript_3b_collate_phasing_stats.R’ was used to collate phasing stats and generate a table with the proportions of reads mapped to either reference, which can give insights into the proportions of haplotypes and ploidy levels.

#### Processing of new haplotype accessions in HybPiper and HybPhaser part I

The newly generated read files of the phased accessions were passed through the HybPiper and HybPhaser part I pipelines using the R scripts C1 and R1a-d with the same settings as for the non-phased samples. For the dataset optimization step, we removed the same 24 loci flagged as putative paralogs for the non-phased accessions and performed the paralog removal step for each sample individually flagging and removing statistical outliers using the scripts (Appendix 10).

#### Combined dataset of phased and non-phased accessions

In order to analyze the phased accessions in the context of the whole genus, we generated a dataset that combined the phased accessions with all non-phased accessions using the R scripts ‘Configure_4_Combining_sequence_lists.R’ and ‘Rscript_4a_combine_phased_with_normal_sequence_lists.R’. Alignments and phylogenetic analyses were then performed with the same methodology as was used for the framework phylogeny using the consensus sequences.

HybPhaser is compatible with Linux and consists as a collection of bash and R scripts that are protected under the terms of a free software license agreement GNU and are available online on GitHub: https://github.com/LarsNauheimer/HybPhaser. Sequence data generated for this study is available at the NCBI sequence read archive.

## Results

### HybPiper sequence assembly

The HybPiper assembly was performed successfully for 307 of the 310 samples (Appendix 3). Paralog warnings were issued for 38 samples and between one and five genes per sample, on average 0.16 paralogous genes per sample.

### Reducing proportions of missing data

A total of 20 samples and 41 loci were excluded from the dataset (Appendix 5) due to poor sequence recovery. Ten of the excluded samples had less than 20% of loci recovered, and all 20 had a total sequence length of less than 45% of the target sequence length recovered. Thirty-nine loci had sequences recovered for less than 20% of the samples and five loci had on average across all samples less than 50% of the target sequence length of the gene recovered. The remaining dataset comprised 290 samples and 312 loci.

### Removal of paralogous genes

24 loci were flagged as putative paralogous genes across all with an average proportion of SNPs of 0.0062 or more (Appendix 6). On average 7.5 loci per sample were flagged as putative paralogs by comparing the SNPs between all genes in each sample individually.

### Assessment of locus heterozygosity and allele divergence

The sampled known hybrids displayed considerably higher values of heterozygosity (LH) and allele divergence (AD) compared to other samples, although the distinction between hybrids and non-hybrids was not clear cut. Across all samples, LH varied gradually between 10.3% and 97.4% and AD varied between 0.03% and 1.51%. Known and suspected hybrids had an average of 89.4% LH and 0.86% AD, while all other samples had an average of 52.3% LH and 0.21% AD (Fig. 2A, Appendix 7). All included hybrids except one had a LH of more than 80%. The hybrid accession *Nepenthes boschiana* × *glandulifera*, which had poor sequence recovery (50.5% of target sequence length before optimization), had very low values of LH (41.6%) and AD (0.28%) compared to the other hybrids.

### Framework phylogeny alignment generation

The supermatrix dataset consisted of 285 genes and 290 taxa, with an aligned sequence length of 423,105 base pairs (bp) and 30.4% missing data. Overall, clades in the framework phylogny were well supported, and largely corresponded to the clades retrieved by Murphy et al. (2020) (Fig. 2B). Many clades obtained maximum bootstrap support, although the gCF values were often low. Some of the backbone nodes received very low support. Most hybrid accessions fell within the clade containing one of the parental taxa where the hybrid was often found in a basally diverging position within the clade (e.g., *N. ampullaria* Jack *× gracilis* Jebb & Cheek; Fig. 2B).

### Clade association

Interpretation of clade association must be done for each sample individually, taking into consideration the phylogenetic distance between that sample and the clade references, the phylogenetic distance between clade references, and the sequence length of the clade reference sequences. Most samples had a single clade association with a higher proportion of reads matching unambiguously to one clade reference compared to the others (Fig. 3, Appendix 8). In contrast, all known hybrids showed associations with multiple often divergent clades. In most cases the association with both clades is strong (e.g., *N. ampullaria* × *gracilis*) in others the association with one clade is weak (e.g., *N. truncata* Macfarl. × *ventricosa* Blanco) or very weak (e.g., *N. clipeata* Danser*× ventricosa*). Even *N. boschiana* × *glandulifera*, which had low values of LH and AD, showed association to both parental clades. Nine non-hybrid accessions with high LH (>80%) showed associations with two clades (e.g., *N. kampotiana*-1 Lecomte, *N. bellii*-1 K.Kondo, *N. xiphioides*-1B.R.Salomon & R.G.Maulder). Samples with lower values of LH generally showed either a single clade association, or multiple associations of closely related references.

**Figure 3.**
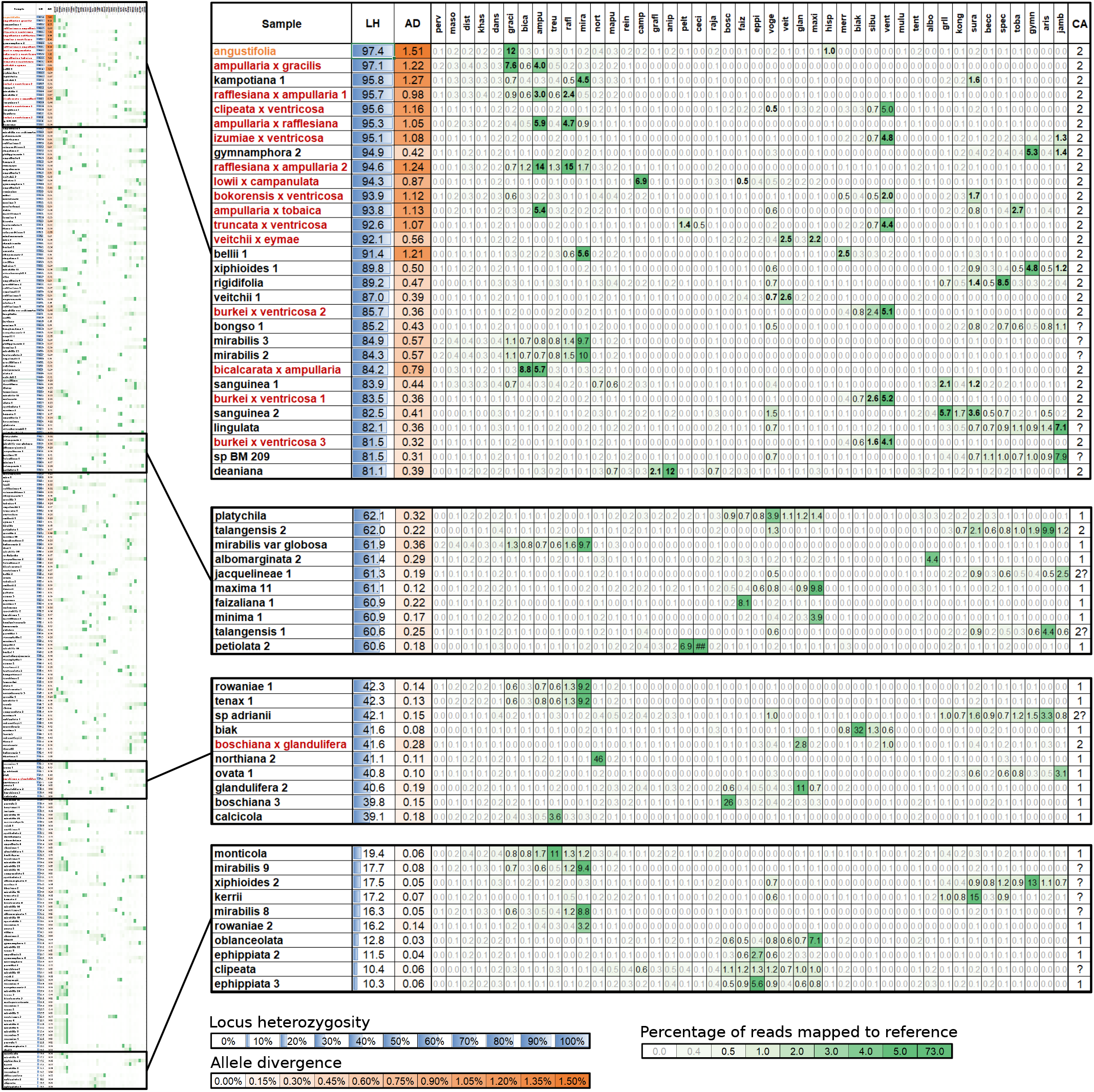
Clade association table and heatmap displaying the percentage of reads matching to each of the 44 clade references. Complete table on left with extracts of example rows shown on the right. Table includes locus heterozygosity (LH) and allele divergence (AD), percentages of reads matching to each reference, and number of clade associations (CA). Known hybrids in red, putative hybrid in orange.

The highest proportions of matching reads were found in phylogenetically distinct, basally diverging lineages (e.g., > 60% in *N. khasiana* Hook.f.), while multiple clade associations were often found in taxa with low LH and AD when they were not closely related to one or in between two clade references (e.g., *N. petiolata* Danser, *N. clipeata*). Few samples have reads associated with multiple divergent clades even though their LH and AD was very low (e.g., *N. spathulata* Danser, *N. talangensis* Nerz & Wistuba).

### Phasing of sequence reads

In mapping onto the selected references, an average of 4.9% (1.1–13.9%) of original sequence reads mapped unambiguously to a single reference. Thirteen phased accessions had approximately equal proportions of unambiguous mapped reads; six accessions had a proportion of approximately 3:2, nine accessions had proportions of approximately 2:1, and four accessions had a proportion of roughly 1:3 (Appendix 9).

### Processing of haplotype accessions

Most phased accessions had considerably lower heterozygosity and allele divergence compared to the non-phased accessions before phasing (Appendix 10). Both phased accessions of *N. boschiana* × *glandulifera* had very low sequence recovery (35.5% and 10.5%) and were removed from the dataset. The paralog detection for each sample individually resulted in flagging of on average 6.8 genes per sample as putative paralogs that were removed from the dataset (Appendix 10).

### Combined dataset of phased and non-phased accessions

The combined dataset consisted of 285 genes and 315 taxa, with an aligned sequence length of 400,685 base pairs (bp) and 32.6% missing data.

The topology of the combined phylogeny is congruent with the framework phylogeny in all nodes, but with stronger support (Fig. 4). The support for the backbone nodes and for the clades that contained hybrids increased compared to the framework phylogeny with non-phased accessions due to the reduction of conflicting phylogenetic signal. This was especially prominent regarding gene concordance factors showing that phasing hybrids reduced conflicting information and increased concordance in the dataset.

**Figure 4.**
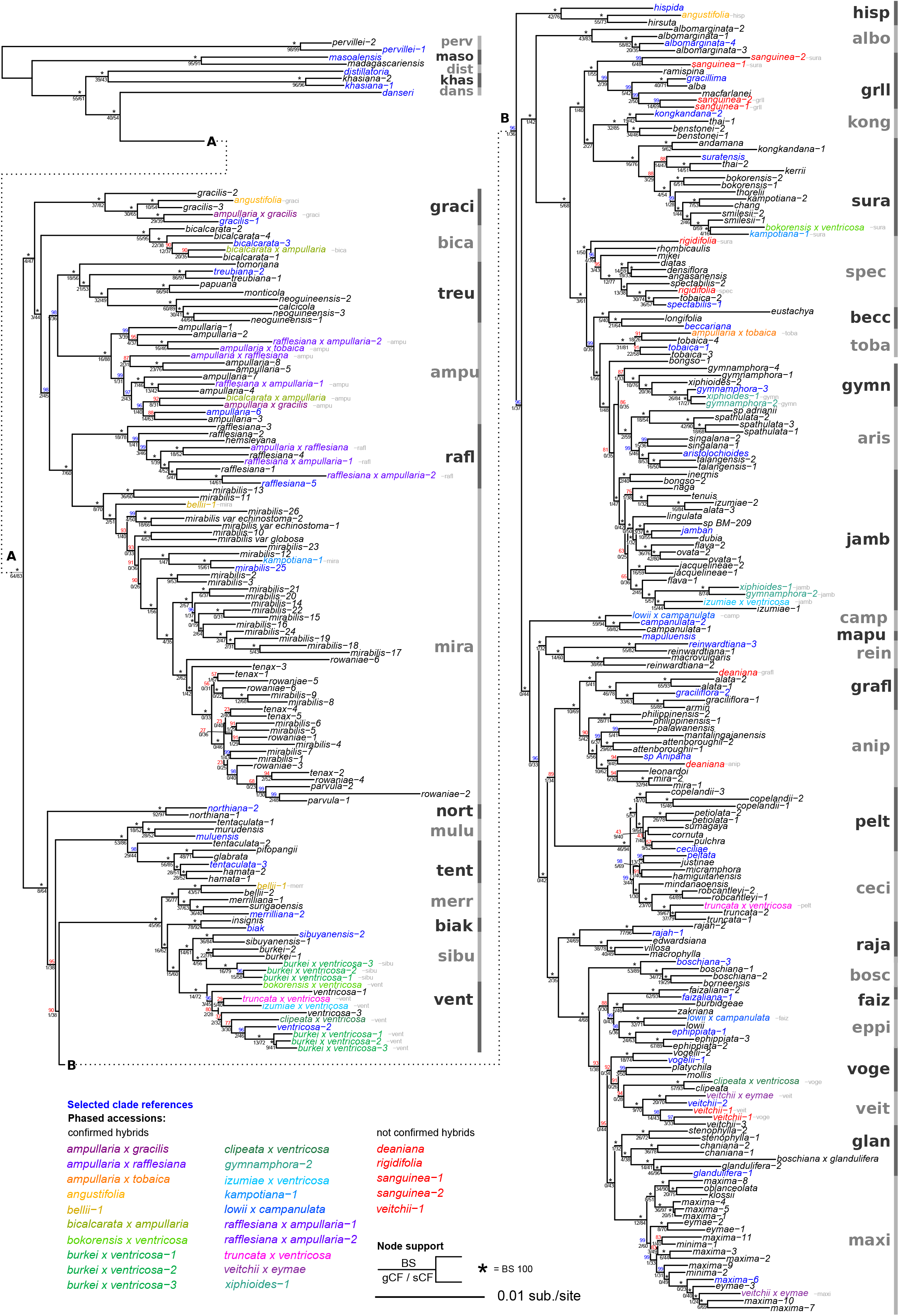
Phylogenetic tree of consensus supermatrix including phased haplotype accessions. Phased accessions are displayed in different colors. Clade references are displayed in blue. Node support is shown above the node in bootstrap (BS) (*=BS100), and below the node in gene and site concordance factors (gCF/sCF).

The phased haplotype accessions from all 16 known hybrids grouped in the clade of their respective parental species, mostly as sister to the parental species, e.g., ‘*N. ampullaria × gracilis* to graci grouped with *N. gracilis*-1 and ‘*N. ampullaria × gracilis* to ampu’ groups with *N. ampullaria*-6 and *N. ampullaria*-3. The phased accessions of the putative hybrid *N. angustifolia* grouped in divergent clades with *N. gracilis*-3 and *N. hirsuta* Hook.f.. Of the nine accessions previously considered to be non-hybrids but with high LH and multiple clade associations, five had phased accessions grouping in divergent clades, three of which with taxa that revealed the parentage, *N. kampotiana*-1 with *N. mirabilis*-25 and *N. smilesii*-1 Hemsl., *N. bellii*-1 with *N. mirabilis*-12 and *N. bellii*-2, and *N. gymnamphora*-2 Reinw. Ex Nees as well as *N. xiphioides*-1 with *N. gymnamphora*-3, and *N. izumiae*-1. The phased accessions of *N. deaniana* Macfarl. grouped more ambiguously with one clade containing *N*. sp-Anipaha and *N. leonardoi* and another clade of *N. alata*-1,2 and *N. graciflora*-1,2. The phased haplotypes of four more accessions, *N. rigidifolia, N. sanguinea*-1,2, and *N. veitchii*-1 did not separate into divergent clades.

## Discussion

### Read phasing with HybPhaser

With the *Nepenthes* dataset, we have shown that phasing reads prior to sequence assembly through HybPhaser is an effective way to detect and characterise haplotypes in hybrid accessions. Previous approaches perform phasing during or after the sequences assembly (Kates et al. 2018, Andermann et al. 2019); however linking reads through these methods using *de novo* requires frequent heterozygous sites to connect reads across the loci (Fig. 1A), which can be difficult to achieve even with read-backed phasing methods using paired-end reads assembly (Kates et al. 2018). Linking phased genes to generate a concatenated multi-gene dataset can be performed by assigning them to clades based on single locus phylogenies, which requires sufficient phylogenetic signal in each locus for successful clade assignment (Tripplet et al. 2012, Estep et al. 2014, Marcussen et al. 2015). In HybPhaser, sequence reads are phased by mapping to multiple reference sequences representing parental clades prior to their assembly, avoiding these issues.

Although this pre-assembly phasing approach of HybPhaser is advantageous, it must be emphasized that the phased haplotypes only represent an approximation of the parental haplotypes. BBSplit, as used in the HybPhaser workflow, assigns reads to a single phased accession that match unambiguously to its reference while all other reads are assigned to all phased accessions. Therefore, the differences between the references determine the phasing. Theoretically, haplotypes of hybrids will phase perfectly (100% unambiguous matches) if the references are the actual parents of the hybrid sample and no random mutations or cross-over events occurred. However, if the references selected diverge from the actual parental lineages, are heterozygous, or differ in the sequence length or locus coverage, haplotypes will not be phased perfectly and only approximate the parental haplotypes.

Furthermore, the efficacy of the HybPhaser phasing approach is sensitive to the selection of appropriate reference sequences. Firstly, successful phasing will be dependent on the phylogenetic divergence between references; when references are closely related more reads will be ambiguous and the number of successfully matched reads will be low. Secondly, the divergence between the clade reference and the parental lineage of the hybrid has an influence on clade association; with increasing divergence to the clade reference, fewer reads will match unambiguously. We therefore recommend that selected references represent samples that are evenly distributed across the phylogeny, have high loci coverage, and contain little allele divergence. In *Nepenthes*, we chose 44 clade references based on a phylogenetic framework and the assessment of heterozygosity across the dataset. This included relatively closely related references in order to resolve known hybrids with closely related parents (*N. burkei × ventricosa*, and *N. veitchii × eymae*) as well putative hybrids with similarly high LH but low AD (e.g., *N. sanguinea, N. gymnamphora-*2, or *N. xiphioides*-1). Choosing suitable reference sequences was crucial for successfully phasing *Nepenthes* hybrids with parents of differing relatedness using HybPhaser.

### Hybrid detection with HybPhaser

In this study we have shown the utility of HybPhaser in not only characterizing the parentage of known hybrids, but also in detecting previously unknown, putative hybrids. Almost all known hybrid accessions of *Nepenthes* had high LH (>80%) and an AD that reflected the divergence of the parental lineages (most had >0.6%). This contrasts with most samples, which had LH between 20% and 70% and an allele divergence of 0.1% and 0.3% reflecting the occurrence of allelic variation in species. Several samples had an allele divergence between 0.3% and 0.6%, which might indicate the existence of introgression through backcrossing of hybrids into one parental population or loss of duplicated genes after polyploidisation (Mallet et al. 2016, Soltis et al. 2016). Low coverage of sequence reads can lead to low LH and AD values due to the method of variance calling. Here the hybrid *N. boschiana × glandulifera* had by far the lowest LH (41.6%) and AD (0.42%) of all included hybrids. This might be an underestimate due to low sequence coverage, which is indicated by low sequence recovery of the original accession (50.5% of the target sequence length) and the very low sequence recovery of the phased accessions (10.6% and 35.5%). Although *N. borokensis × ventricosa* and *N. gymnamphora*-2 had similarly low sequence recovery in the original mapping, they had normal values of LH and AD and a sequence recovery of phased accessions similar to the normal accessions.

The assessment of heterozygous sites and thus the conflict between the gene variants or divergent haplotypes in the dataset provides a simple way to gain insights into the reticulation of samples. *Nepenthes* is known for a variety of horticultural and natural hybrids that exist (Clarke et al. 2018). We found a gradient of LH and AD across the genus with many samples having intermediate values between known hybrids and most other samples, which indicated introgression and the presence of unknown hybrid accessions. Fourteen samples had LH (>80%) and AD (>0.3%) similar to the closely related hybrids of which nine had reads associated with multiple clades and were phased. After phasing and inclusion of phased accession into the phylogenetic analyses, four samples had phased accessions clearly grouping in divergent clades with specific taxa confirming their hybrid status and revealing parental lineages, *N. bellii*-1 (*bellii* × *mirabilis*), *N. kampotiana*-1 (*smilesii* × *mirabilis*), *N. gymnamphora*-2 (*gymnamphora* × *izumiae*), *N. xiphioides*-1 (*gymnamphora* × *izumiae*). Further, the AD of phased accessions combined from each of the samples was lower than the AD of the non-phased accession. Four samples were not confirmed as hybrids based on the phasing, because the phased accessions did not group in divergent clades (*N. rigidifolia, N*.*sanguinea*-1,2, and *N. veitchii*-1). One sample was more ambiguous; *Nepenthes deaniana* had phased accessions grouping in divergent clades, but not clearly associated with other samples and reducing the phylogenetic support of the affected clades. Further, the AD of the phased accessions combined (0.36% + 0.21%) was much higher than the AD of the not phased accession (0.39%). Therefore, *N. deaniana* cannot be assumed to be a hybrid of these clades.

### Paralog detection with HybPhaser

Our results suggest that the paralog detection of HybPhaser greatly improves on existing methods. In approaches such as the paralog investigator of HybPiper, a paralog warning is only issued when multiple *de novo* contigs are recovered with comparable size, i.e., when multiple contigs have at least 85% of the whole target sequence length (Johnson et al. 2016). In order to successfully detect a paralogous gene, heterozygous sites must occur frequent enough to connect reads of all versions across most of the locus and because the target length is used as reference, the gene must not consist of multiple exons shorter than 85% of the gene. The paralog detection provided in HybPhaser is based on the idea that paralogous genes have higher rates of heterozygous sites than normal genes, and therefore can detect paralogs independent of exon size or distribution of heterozygous sites across the gene.

In *Nepenthes*, HybPiper issued warnings for only 0.16 of genes per sample, while the approach taken by HybPhaser flagged 31.5 genes per sample (c. 10% of the genes). This large difference is not only due to the methodology but also to the chosen conservative approach to remove rather more than less genes. Firstly, flagging genes that have unusual high proportions of SNPs across all samples (here 24 genes) will lead to removing those genes for samples without multiple variants. This might however be desired, as gene loss can lead to leaving only one version active, which is not guaranteed to be the ortholog. Secondly, the method flags genes that have unusual high rates of SNPs due to other reasons than paralog variants, e.g., sequencing or assembly artefacts, indels, or non-paralogous but similar sequences from other genes. HybPhaser provides graphs to assess the distribution of SNPs across all and each sample to set thresholds appropriately and further the option to investigate the origin of high SNP count in single genes by viewing the mapped reads in the generated BAM files.

### Reconciling phylogenetic conflict with HybPhaser

In this study we have demonstrated how HybPhaser can be used to handle phylogenetic conflict in target capture datasets and produce more rigorous phylogenies. Hybrids introduce phylogenetic conflict in the dataset, leading to poorly resolved clades or wrong topology (McDade 1992, Soltis et al. 2008). Depending on the method of sequence generation and dealing with heterozygous sites, this will have different effect. Hybrids might group together with one parent, basal to one, or in between multiple parental clades. Here we used ambiguity codes in the non-phased dataset and most hybrids were in basal position in the clade of one parent while having a shorter total branch length. Phasing of hybrid accession reduced the conflicting signal and improved the clade support of affected clades, especially of gene and site concordance factors. To further decrease the conflicting signal in the dataset one might consider removing all samples with high values of LH or AD. This can be especially useful to increase clade support for the framework phylogeny for clade association.

## Conclusions

HybPhaser provides a novel workflow to detect and phase hybrid accessions in target capture datasets. In this study, we have used HybPhaser to untangle the reticulate evolutionary history of *Nepenthes*, and demonstrated its utility for phasing reads into parental haplotypes, revealing and characterizing hybrids, detecting putative paralogous genes, and resolving phylogenetic conflict.

**Table 1.**
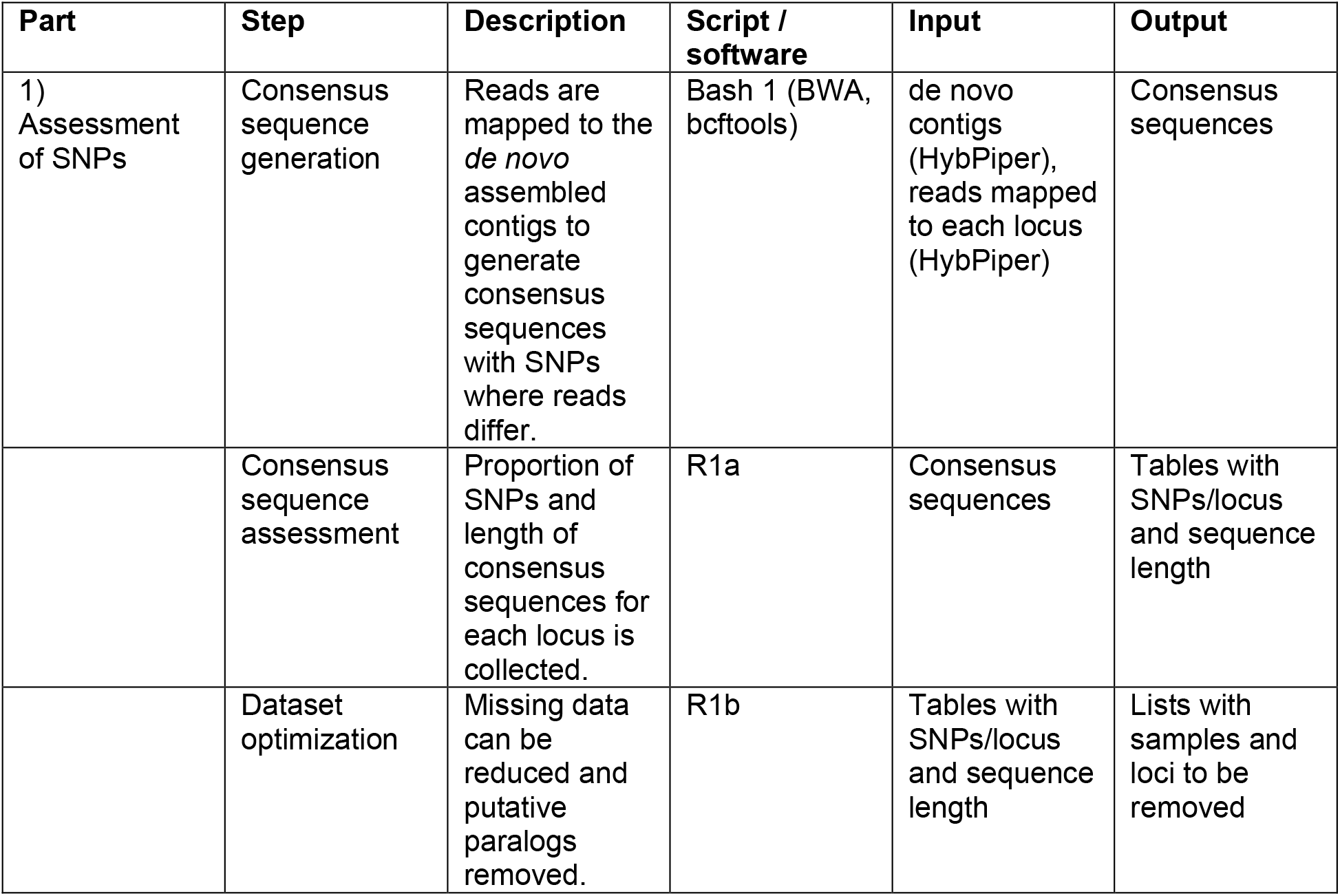
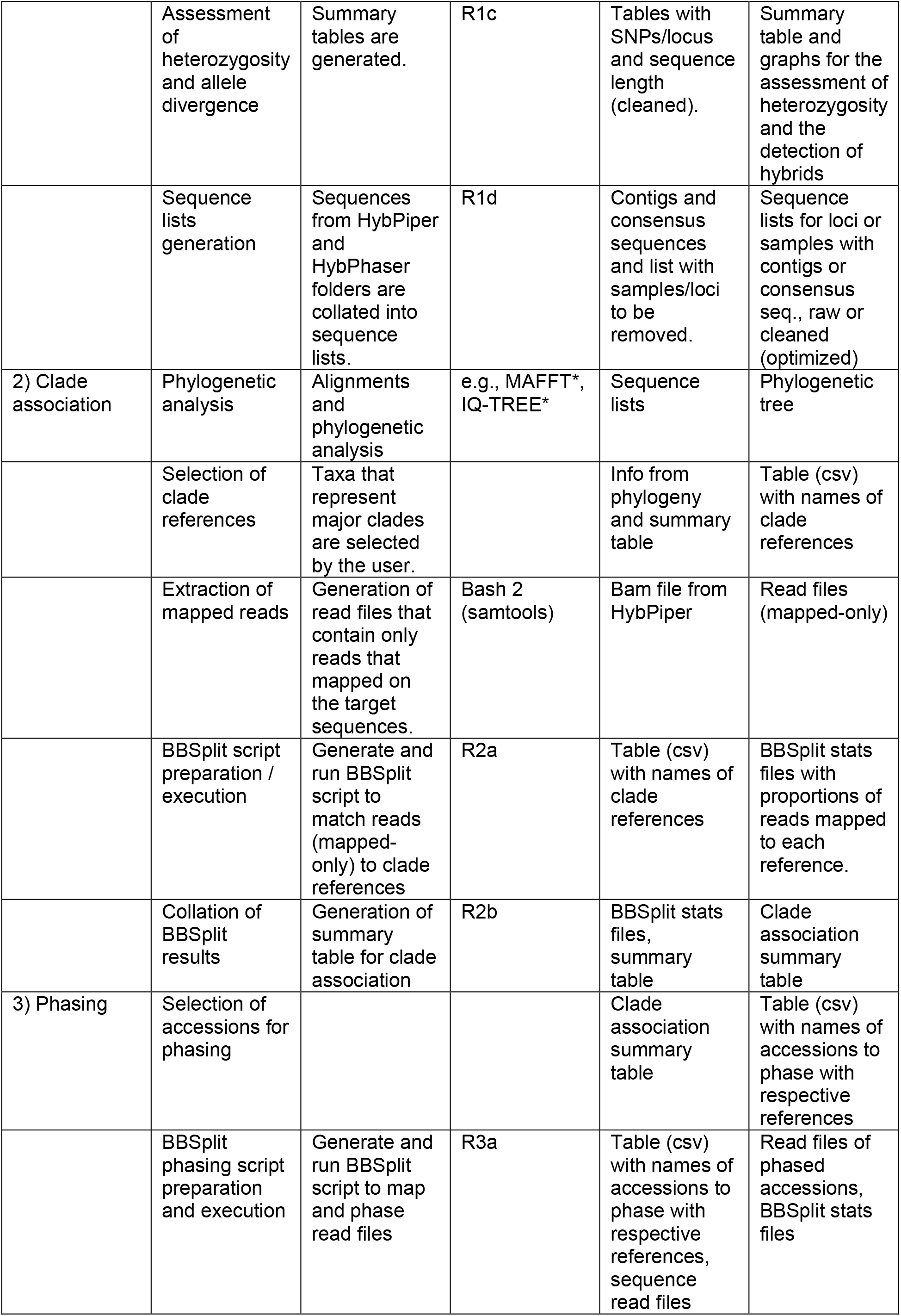
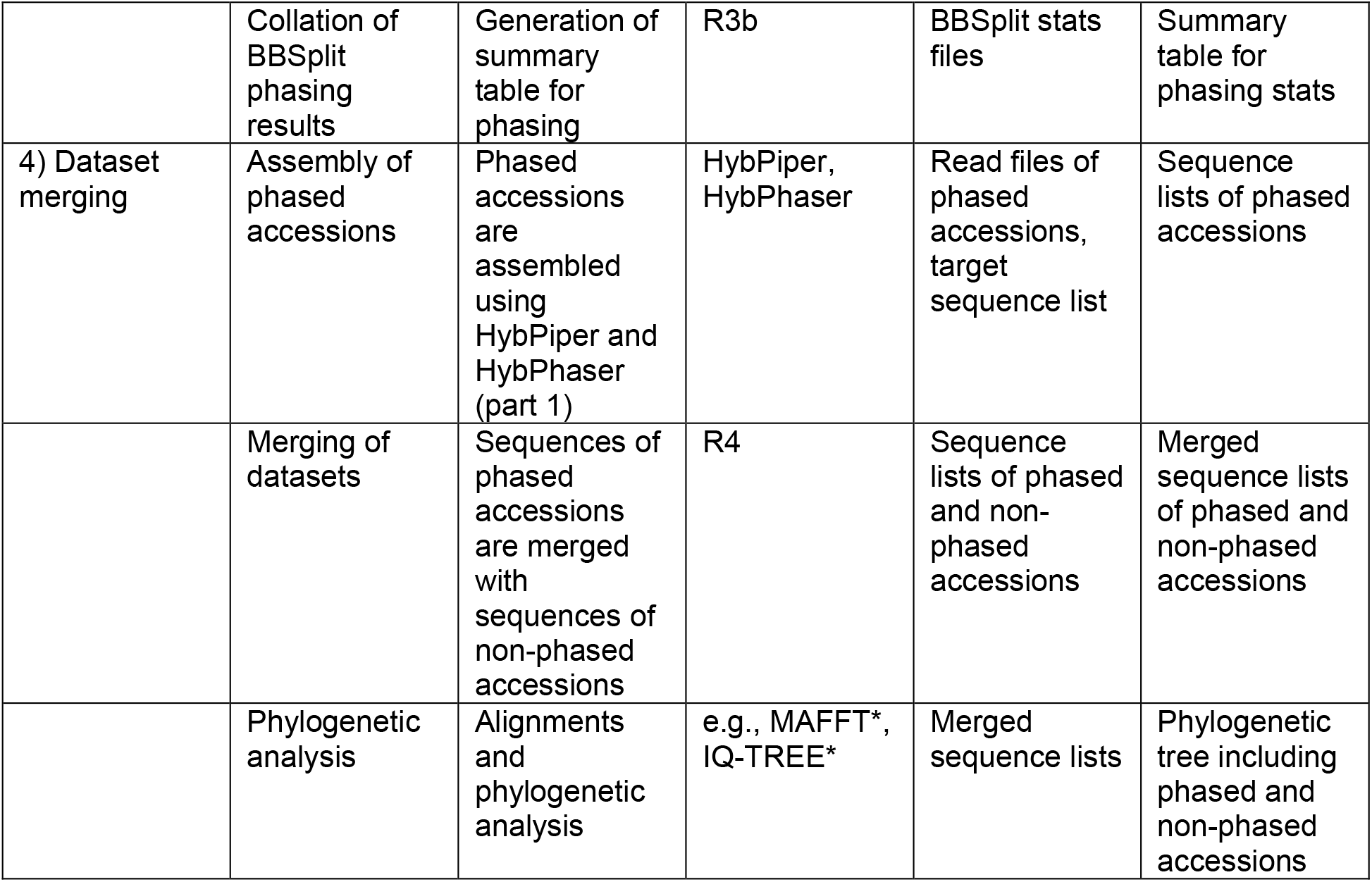
Workflow overview containing all steps with short description, script/software used, required input and generated output for each step. Software marked with an asterisk are not part of the workflow.

| Part | Step | Description | Script / software | Input | Output |
| --- | --- | --- | --- | --- | --- |
| 1)<br>Assessment<br>of SNPs | Consensus<br>sequence<br>generation | Reads are mapped to the *de novo* assembled contigs to generate consensus sequences with SNPs where reads differ. | Bash 1 (BWA, bcftools) | de novo contigs (HybPiper), reads mapped to each locus (HybPiper) | Consensus sequences |
|  | Consensus<br>sequence<br>assessment | Proportion of SNPs and length of consensus sequences for each locus is collected. | R1a | Consensus sequences | Tables with SNPs/locus and sequence length |
|  | Dataset<br>optimization | Missing data can be reduced and putative paralogs removed. | R1b | Tables with SNPs/locus and sequence length | Lists with samples and loci to be removed |
| HybPhaser workflow |  |  |  |  |  |
|  | Assessment of heterozygosity and allele divergence | Summary tables are generated. | R1c | Tables with SNPs/locus and sequence length (cleaned). | Summary table and graphs for the assessment of heterozygosity and the detection of hybrids |
|  | Sequence lists generation | Sequences from HybPiper and HybPhaser folders are collated into sequence lists. | R1d | Contigs and consensus sequences and list with samples/loci to be removed. | Sequence lists for loci or samples with contigs or consensus seq., raw or cleaned (optimized) |
| 2) Clade association | Phylogenetic analysis | Alignments and phylogenetic analysis | e.g., MAFFT*, IQ-TREE* | Sequence lists | Phylogenetic tree |
|  | Selection of clade references | Taxa that represent major clades are selected by the user. |  | Info from phylogeny and summary table | Table (csv) with names of clade references |
|  | Extraction of mapped reads | Generation of read files that contain only reads that mapped on the target sequences. | Bash 2 (samtools) | Bam file from HybPiper | Read files (mapped-only) |
|  | BBSplit script preparation / execution | Generate and run BBSplit script to match reads (mapped-only) to clade references | R2a | Table (csv) with names of clade references | BBSplit stats files with proportions of reads mapped to each reference. |
|  | Collation of BBSplit results | Generation of summary table for clade association | R2b | BBSplit stats files, summary table | Clade association summary table |
| 3) Phasing | Selection of accessions for phasing |  |  | Clade association summary table | Table (csv) with names of accessions to phase with respective references |
|  | BBSplit phasing script preparation and execution | Generate and run BBSplit script to map and phase read files | R3a | Table (csv) with names of accessions to phase with respective references, sequence read files | Read files of phased accessions, BBSplit stats files |
| HybPhaser workflow |  |  |  |  |  |
|  | Collation of BBSplit phasing results | Generation of summary table for phasing | R3b | BBSplit stats files | Summary table for phasing stats |
| 4) Dataset merging | Assembly of phased accessions | Phased accessions are assembled using HybPiper and HybPhaser (part 1) | HybPiper, HybPhaser | Read files of phased accessions, target sequence list | Sequence lists of phased accessions |
|  | Merging of datasets | Sequences of phased accessions are merged with sequences of non-phased accessions | R4 | Sequence lists of phased and non-phased accessions | Merged sequence lists of phased and non-phased accessions |
|  | Phylogenetic analysis | Alignments and phylogenetic analysis | e.g., MAFFT*, IQ-TREE* | Merged sequence lists | Phylogenetic tree including phased and non-phased accessions |

## Supporting information

Appendix 1 - Manual

Appendix 2 - Table of accessions

Appendix 3 - HybPiper results

Appendix 4 - Table SNPs

Appendix 5 - Dataset optimization

Appendix 6 - Summary table

Appendix 7 - Phylogeny normal

Appendix 8 - Clade association Table

Appendix 9 - Phasing

Appendix 10 - Processing of phased accessions

## Acknowledgements

This work was supported by the Australian Biological Resource Study (Department of the Environment, Australian Government, ABRS grant RFL214-62) as well as the Australian and Pacific Science Foundation (APSF 16/4). We are thankful to Melissa Harrison (Australian Tropical Herbarium) for support in the laboratory and Todd McLay and Chris Jackson from (Genomics for Australian Plants consortium and the Royal Botanic Gardens Victoria) for feedback on the HybPhaser workflow.

