## Appendix 1 - Manual for "HybPhaser: a workflow for the detection and phasing of hybrids in target capture datasets"

##### ****Appendix 1 - HybPhaser Manual****

##### ****Overview****

In the first part (SNPs assessment), HybPhaser is used to measure and assess heterozygous sites in the dataset to detect hybrid accessions as well as putative paralogous genes. It can further be used to optimize the dataset by reducing missing data and removing those putative paralogs. Finally, consensus sequences (HybPhaser) or de novo contigs (HybPiper) can be collated to generate sequence lists ready for alignment and phylogenetic analyses.

In the second part (Clade association), the association of read towards divergent clades is assessed to detect hybrid accessions that can be phased into multiple accessions. The framework phylogeny is used to select suitable references for all major clades across the studies group. The software BBSplit is then used to simultaneously map sequence reads to all clade references recording the proportion of reads matching unambiguously to a single reference. HybPhaser R scripts facilitate running the BBSplit analysis and collate the results into a summary table in order to select suitable accessions that can be phased.

In the third part (Phasing), BBSplit is used to simultaneously map the sequence reads of selected accessions to relevant clade references and distribute reads into new read files representing accessions of phased haplotypes. HybPhaser R scripts facilitate the running of BBSplit and generate a summery table of the mapping stats.

Finally, the newly generated read files can then be assembled using HybPiper and HybPhaser similar to the original read files. HybPhaser R scripts can then be used to combine sequence lists and generate a dataset of phased and non-phased accessions. Phylogenetic analysis of this combined dataset can then reveal the origin of hybrid taxa and reduce phylogenetic conflict for better resolved phylogenies.

##### Main Principles of the approach

**Mapping reads to de novo contigs to detect heterozygosity**

Target capture sequence assembly generally relies on de novo assembly, which can lead to chimeric sequences in the presence of divergence between gene variants. While reference mapping can capture heterozygous sites by assigning ambiguity codes, it relies on a reference for sequence assembly. HybPhaser maps the sequence reads back onto the de novo contig and generates a consensus sequence that codes heterozygous sites (SNPs) with ambiguity codes. The assessment of SNPs across samples and genes can then be used to detect hybrids (high proportions of SNPs in most genes compared to other samples) and paralogous genes (higher proportion of SNPs in a single gene compared to other genes).

**Phasing of reads by mapping to multiple references simultaneously**

Hybrid accessions contain sequence reads from both parental lineages. If there is sufficient divergence between the parental lineages, the reads can be associated with different clades by mapping them simultaneously to multiple references and recording to which they map best. HybPhaser applies this principle in two steps. First the association of reads with clades across the studied group is assessed by mapping reads from all accessions to several references representing clades. Samples that have reads associated with more than one clade can then be phased by mapping only to the relevant references and separating the read files accordingly.

**Installation**

HybPhaser scripts can be downloaded from GitHub

git clone https://github.com/larsnauheimer/HybPhaser.git

**Software dependencies**

- [R (v4.0)](https://cran.r-project.org/)
- R-packages:
  - ape (v5.4)
  - sequinR (v4.2)
  - stringR (v1.4)
- [BWA](http://bio-bwa.sourceforge.net/)
- [SAMtools](https://github.com/samtools/)
- [Bcftools](https://samtools.github.io/bcftools/)
- BBSplit ([BBMap](https://sourceforge.net/projects/bbmap/) v.38.87)
- [HybPiper](https://github.com/mossmatters/HybPiper) (which has additional dependencies)

**Data Preparation**

Prior to run HybPhaser, sequence assembly has to be performed using **HybPiper** [(Johnson et al. 2016)](https://bsapubs.onlinelibrary.wiley.com/doi/abs/10.3732/apps.1600016). The use of HybPiper is well-explained in the [HybPiper-Wiki](https://github.com/mossmatters/HybPiper/wiki).

HybPiper requires the sequence reads and a fasta file with the target sequences.

In short, HybPiper pre-selects reads that match to a target gene using BWA or BLAST and then performs a *de novo* assembly using Spades of the pre-selected read files. Extronerate is then used to extract and concatenate exon regions to generate gene sequences. Optionally intronerate can be used to recover any available intron regions and concatenate these with the exons to 'supercontigs'.

HybPiper generates one folder for each sample, which contains subfolder for each gene that contains assembly files as well as the assembled contig sequences. HybPhaser uses several of the output files of HybPiper for further analyses. The 'cleanup.py' script of HybPiper can be used before running HybPhaser.

**Output files from HybPiper used in HybPhaser:**

reads mapped to each locus:

- sample/gene/sample/gene_unpaired.fasta (for single-end reads)
- sample/gene/sample/gene_interleaved.fasta (for paired-end reads)

contig sequence as reference:

- sample/gene/sample/sequences/FNA/gene.FNA (for normal contigs)
- sample/gene/sample/sequences/intron/gene_supercontig.fasta (for supercontigs)

BAM file used to extract reads that mapped to the target sequences:

- sample/sample.BAM

**How to use HybPhaser**

HybPhaser consists of two bash scripts and several R scripts that can be executed from the main script. Four configuration scripts (one for each part) have to be adjusted before the remaining R-scripts can be run line by line from the main script.

- **Configuration scripts**: to adjust all necessary variables
- **Main script**: for the execution of all other R scripts
- **Sub-scripts**: to be left alone

**Data preparation**

1. assembly sequences in HybPiper

**SNPs assessment**

2. generate consensus sequences (bash)
3. adjust config script 1 (R)
4. run scripts for part 1 in main script (R)
5. perform alignments and phylogenetic analyses

**Clade association**

6. extract mapped reads (bash, optional)
7. adjust config script 2 for (R)
8. run scripts for part 2 in main script (R)

**Phasing**

9. adjust config script 3 (R)
10. run scripts for part 3 in main script (R)

**Combine datasets**

11. adjust config script 4 (R)
12. run scripts for part 4 in main script (R)
13. perform alignments and phylogenetic

### 1. SNPs assessment

The detection of hybrids (and putative paralogous genes) relies on the assessment of SNPs, which is performed in several steps.

Firstly, consensus sequences have to be generated by re-mapping the sequence reads using the de novo contigs (from HybPiper) as references, which is performed using the command line script "generate_consensus_sequences.sh". Then, the SNPs information coded in consensus sequences is collected using R to generating tables and graphs for dataset optimization, assessment of heterzygosity and allele divergence and finally to generate sequence lists that are ready for alignment and phylogenetic analyses.

All steps execpt the first one are perfomed in R using the main script to execute sub-scripts and the configuration script to set all variables.

#### 1.1. Consensus sequence generation

Consensus sequences summarize the information of assembled sequence reads and can contain ambiguity characters where reads differ. Differences in sequence reads can occur through divergent alleles, paralogous genes, sequencing errors, or contamination. The generation of consensus sequences allows to measure and assess the occurrence of heterozygous sites across the dataset and to identify hybrid taxa as well as paralogous genes. To generate consensus sequences, HybPhaser maps the reads mapped of a single locus to the de novo assembled contig of that locus. Both files are available in the HybPiper output. HybPhaser uses BWA for the mapping and bcftools for the variant calling and consensus sequence generation for which the settings can be adjusted. The consensus sequences contain ambiguity characters coding for SNPs.

The bash script “generate_consensus_sequences.sh” is run from the command line. Input: de novo contigs (HybPiper), reads mapped to each locus (HybPiper) Output: Consensus sequences and processing files for mapping and variant calling (written in HybPiper folder structure: /sample/gene/sample/sequences/remapping/)

generate_consensus_sequences.sh

**Options:**

- -n Name of sample (required)
- -p Path to HybPiper results folder. Default is current folder.
- -t Maximum number of threads used. Default is 1.
- -s (without argument) If chosen, supercontigs are used instead of normal contigs.
- -d Minimum coverage on site to be regarded for assigning ambiguity code. If read depth on that site is lower than chosen value, the site is not used for ambiguity code but the most frequent base is returned. Default is 10.
- -f Minimum allele frequency regarded for assigning ambiguity code. If the alternative allele is less frequent than the chosen value, it is not included in ambiguity coding. Default is 0.15.
- -a Minimum count of alleles to be regarded for assigning ambiguity code. If the alternative allele ocurrs less often than the chosen value, it is not included in ambiguity coding. Default is 4.

Examples for a single sample

generate_consensus_sequences.sh -n sample1 -p ~/hybpiper_results -t 4 -s

or using a list of samples (similar to the HybPiper nameslist)

while read NAME; do; generate_consensus_sequences.sh -n $NAME -p ~/hybpiper_results -t 4 -s; done < nameslist.txt

Note: The BAM files from the remapping step can be very useful for understanding why some samples or genes have high amount of heterozygous sites. Opening the BAM files in a visual viewer (e.g. Geneious) can give valuable insights into the data, e.g. help deciding whether a gene/exon is paralogous or whether a sample is contaminated.

#### 1.2. Consensus sequence assessment

The R script “Rscript_1a_count_snps_in_consensus_seqs.R” is used to generate a table with the proportions of SNPs in each locus and sample as well as a table with sequence length. Ambiguity characters that code for 3 different nucleotides are weighted twice of the ones coding for 2 different nucleotides.

The script requires several variables to be set in the configuration script “Configure_1_SNPs_assessment.R”:

- path_to_hybpiper_results: set path to the HybPiper base folder
- path_for_HybPhaser_output: set path for HybPhaser output`
- fasta_file_with_targets: direct to the file with target sequences (baits)
- txt_file_with_list_of_accessions: direct to a file that contains all sample names (one line per sample)
- contig: select contig type, either “normal” for standard HybPiper assembly or “supercontig“ for the supercontig assembly, if intronerate.py was used to generate concatenated exon sequenes in HybPiper.

#### 1.3. Dataset optimization

The R script “Rscript_1b_optimize_dataset.R” can be used to reduce the proportion of missing data by removing low quality samples and loci as well as removing putative paralog loci.

It is recommended to run the dataset optimisation script first, assess the results (graphs and tables) and then decide on thresholds that are relevant for the used dataset.

**Missing data:**

HybPhaser graphs and summary statistics for missing data in samples and loci, as well as recovered sequence length for samples as proportion of the target sequence (or the longest target sequence, if multiple exist for a gene). Thresholds can be set to determine which samples are to be excluded from the dataset.

These variables can be configured in the configuration script:

- remove_samples_with_less_than_this_propotion_of_loci_recovered: set threshold for samples to be removed that have a low proportion of the number of loci recovered (e.g. 0.8 will only keep samples that have a sequence recovered for at least 80% of loci)
- remove_samples_with_less_than_this_propotion_of_target_sequence_length_recovered: set threshold for samples to be removed that have short sequence length recovered (as proportion of the target sequence length). (0.6 will only keep samples that have on average 60% of the target sequence length recovered)
- remove_loci_with_less_than_this_propotion_of_samples_recovered: set threshold for loci to be removed that have a low proportion of the number of samples recovered (e.g. 0.8 will only keep loci that have a sequence recovered for at least 80% of samples)
- remove_loci_with_less_than_this_propotion_of_target_sequence_length_recovered: set threshold for loci to be removed that have short sequence length recovered (as proportion of the target sequence length). (e.g. 0.6 will only keep loci that have on average 60% of the target sequence length recovered)

**Putative paralogs:**

Genes with unusually high proportions of SNPs compared to other genes might have heterozygous sites due to the presence of paralogous genes or contamination. Recent paralogs might be restricted to a single species, while ancient paralogs can be shared across many taxa. Here loci can be removed from the dataset that have high proportion of SNPs across all samples as well as that are 'outlier' in a sample.

These variables can be configured in the configuration script:

- remove_loci_for_all_samples_with_more_than_this_mean_proportion_of_SNPs: set threshold to remove loci with high mean proportions of SNPs across all samples. (e.g. 0.02 will remove all loci with an average of 2% or SNPS across all samples, or "outliers" will remove all loci that have more than 1.5*IQR (interquartile range) above the 3rd quartile of mean SNPs, "none" or 0 will not remove any loci)
- file_with_putative_paralogs_to_remove_for_all_samples: set path to file that contains list of putative paralogs to be removed for all samples. Tthis can be used for the processing a subset of accessions and the list of paralogs from another analysis should be imported (e.g. of phased accessions). OPTIONAL! Using "" will result in applying previous selected mode or threshold.
- remove_outlier_loci_for_each_sample: set whether outlier loci are to be removed per sample. "yes" will remove for each samples loci that have a higher proportion of SNPs than 1.5*IQR (interquartile range) above the 3rd quartile.

#### 1.4. Assessment of heterozygosity and allele divergence

The script “Rscript_1c_summary _table.R” generates a table summarizing the results from the SNPs assessment after applying the dataset optimization steps. It can be used to assess the proportion of loci with any SNPs (heterozygosity) and the proportion of SNPs (allele divergence) in order to detect hybrid samples.

Hybrids are expected to have a high heterozygosity (>90%) as well as a considerable allele divergence (>1%). However, these values are relative and can vary depending on the dataset, targets, sequencing quality, study group and many other factors. They should be carefully evaluated.

High allele divergence can indicate a large population with diverse allele variants or in hybrids it can be the divergence between the parental alleles and correlate to the divergence of the parental species. A low allele divergence can indicate low allele diversity, e.g. occurring in isolated populations.

Be aware, that low values of heterozygosity and allele divergence can be due to low coverage in the remapping. It might be useful to adjust the thresholds for variance calling in step 1.1.

#### 1.5. Sequence lists generation

The R script “Rscript_1d_generate_sequence_lists.R” collects de novo assembled contig sequences (HybPiper) and remapped consensus sequences (HybPhaser) and creates sequence lists for each locus and for each sample as well as without or with the dataset optimization applied. In total eight folder are generated that contain the sequence lists in fasta format:

- consensus_sample_raw
- consensus_sample_clean
- consensus_loci_raw
- consensus_loci_clean
- contig_sample_raw
- contig_sample_clean
- contig_loci_raw
- contig_loci_clean

The loci sequence lists can be aligned in order to perform phylogenetic analyses. The sample sequence lists can be used as references for the clade association and the phasing steps.

There is a complication with the contig_sample_x sequence lists of normal contigs (not supercontigs), which makes their generation very slow: HybPiper does not write the gene name into the fasta file but only the sample name. The sequence list generation of HybPhaser uses the fast unix command cat, which does not work when the gene names are not in the sequence names. The work around takes much more time.

To save time you can turn off the slow work around by setting the variable write_gene_names_in_contig_sample_seqlist to "no". The generated sequence lists per sample for contigs (or normal contigs) will then not contain the gene names.

### 2. Clade association

Hybrid accessions contain sequence reads from divergent parents, which can be phased, if the parents are sufficiently divergent. The clade association step of HybPhaser maps all reads onto multiple references representing divergent clades and thus identify hybrids with divergent parents. The software BBSplit (BBMap) is used to simultaneously map the reads to the multiple references and record the proportion of reads mapped unambiguously to each reference.

The configuration script **Configure_2_Clade_association.R** is used to set required variables for sub-scripts to be run in the main script. They will generate a command line script that executes the BBSplit analysis (2a) and summarizes the results (2b).

#### 2.1. Selection of clade references

Based on a framework phylogeny (using the generated sequence lists), the user has to select suitable clade references. Ideally, clades should be selected that have similar phylogenetic distance across the phylogeny. Too many clades will result in less reads mapping unambiguously towards a single reference, and too few clades will result in accessions not mapping to any reference. Further accessions chosen to represent clades should have low heterozygosity and low allele divergence but high sequence recovery (number of loci sequenced and proportion of target sequence). A table (CSV format) listing the clade references as well as an abbreviated name needs to be generated by the user.

#### 2.2. Extraction of mapped reads

Read files vary in the proportion of reads that match the target sequences, which has impact on the proportion of reads mapping to the clade references. It is therefore recommended to only use reads for the clade association that mapped to the target sequences. HybPiper generates a bam file that marked all reads that mapped to the target sequences, which can be extracted using the command line script **extract_mapped_reads.sh**. The script generated new read files, which can be used for the clade association analysis.

extract_mapped_reads.sh [options]

**Options:**

- -n Text file with list of samples (one line per samples)
- -p Path to HybPiper results folder. Default is current folder.
- -o Path to output folder where new read files are saved. Default is a folder called _mapped_reads in the HybPiper results folder."

Note: For an even more precise clade association assessment one can use BBNorm (BBMap package), which can be used to normalize the coverage in the mapped reads. The coverage of mapped reads is generally higher in the center of the targeted exons and drop towards the margins. If the sequencing coverage is high enough, a normalization step can be beneficial.

#### 2.3. BBSplit script preparation / execution

The R script **Rscript_2a_prepare_bbsplit_results.R** can be used to generate the executable command line files that run the clade association mapping. It can also execute the script in R, although the mapping is time consuming and might be rather executed from the command line.

Variables to be set in the configuration script **Configure_2_Clade_association.R**:

- path_to_HybPhaser_results: path to HybPhaser result folder (e.g. .../HybPiper/_HybPhaser/)
- path_to_clade_association_folder: path to the output folder; best as subfolder in the HybPhaser folder (e.g. ".../hybpiper_results/HybPhaser_supercontig/clade_association_1")
- csv_file_with_clade_reference_names: set filename for the previously prepared text file with reference sample names
- path_to_read_files_cladeassociation: set folder that contains sequence read files (e.g. ".../HybPiper/_mapped_reads/")
- read_type_cladeassociation: set whether reads are single or paired-end reads (single end, if you use the mapped-only reads) ["single-end" or "paired-end"]
- ID_read_pair1: if reads are paird-end, set unique part of filename including the ending (e.g. "_R1.fastq", "_R2.fastq"). If reads are single-end, you can ignore this.
- ID_read_pair2: similar to “ID_read_pair2”
- path_to_reference_sequences: set folder that contains the sequences of samples to select the clade reference sequences (e.g., .../HybPiper_results/HybPhaser/sequences/consensus_samples_raw")
- path_to_bbmap: set folder to bbmap binaries, if bbmap is not in your path variable
- no_of_threads: set number of threads to be used for BBSplit [any number]
- run_bash_script_in_R: select whether the script is run directly in R [“yes”] or whether to run it manually from command line using the generated script

#### 2.4. Collation of BBSplit results

The script **Rscript_2b_collate_bbsplit_results.R** generates a summary table based on the BBSplit output files. It displays the proportions of reads from each sample mapping unambiguously to each of the clade references. This table is essential to select accessions that can be phased.

#### Notes

The proportions of associated reads vary with the references. If references are closely related, less reads are unambiguous for either, if they are too far apart reads may not map unambiguous at all.

It can therefore be useful to run multiple clade association analyses with more or less clades, or with different references. It might also be adventageous to run first a wide scale association and then a fine-scale association.

### 3. Phasing

Hybrids with reads associated to multiple clades can be phased accordingly into phased accessions that approximate the phased haplotype accessions. BBSplit has the option to map reads to multiple references and distribute reads to new files. Here it is used so reads that map unambiguously to a single reference are assigned to only the regarding read file, while reads that map ambiguously to multiple references are assigned to all read files. This generated phased accessions that differ in the relevant sequences where SNPs occur.

#### Configure_3_Phasing.R

HybPhaser scripts are used to facilitate the application of BBSplit by generating an executable command line script and generates a summary table of the BBSplit results. All variables for the phasing are set in the script **Configure_3_Phasing.R** and the relevant script is executed in the main script.

#### 3.1. Selection of accessions for phasing

The selection of suitable accessions for the phasing step has to be done by the user based on the clade association. A table (CSV format) listing the accession to be phased as well as the relevant references needs to be generated by the user. The file needs to be a comma separated file with the following header (columns): samples,ref1,abb1,ref2,abb2,ref3,abb3,...

- samples is the sample name that should be phased
- ref1 is the sample name of the 1st reference
- abb1 is the abbreviation for the 1st reference
- …

#### 3.2. BBSplit phasing script preparation and execution

The script **Rscript_3a_prepare_phasing_script.R** can be used to generate the command line script that executes the BBSplit phasing step, or it can be run inside the R script.

**Variables to set in the configure script:**

- path_to_HybPhaser_results: set HybPhaser basefolder
- path_to_read_files_phasing: set path to read files that should be phased
- read_type_phasing: set whether reads are paired-end or single-end ["paired-end" or “single-end"]
- ID_read_pair1: if reads are paird-end, set unique part of filename including the ending (e.g. "_R1.fastq")
- ID_read_pair2: if reads are paird-end, set unique part of filename including the ending (e.g. "_R2.fastq")
- path_to_reference_sequences: set folder for reference sequences (e.g. "/sequences/HybPhaser_samples_raw")
- path_to_bbmap_executables: set path to bbmap executables (if not in path)
- path_to_phasing_folder: set path to phasing output folder
- csv_file_with_phasing_prep_info: set CSV file with accessions to phase and relevant references
- no_of_threads_phasing: set number of maximum used threads [any number]
- run_bash_script_in_R: set whether to run the script to run the bbsplit analyses in R instead of in the terminal ["yes" or "no"]
- folder_for_phased_reads: set output folder for phased reads
- folder_for_phasing_stats: set output folder for phasing stats

#### 3.3. Collation of BBSplit phasing results

The script **Rscript_3b_collate_phasing_stats.R** generated a summary table that displays the proportion of reads that matched unambiguously to each of the references the accession was phased to. The ratio of reads mapped to different references can give insights into the ploidy level, e.g. a F1 hybrid between two parents should roughly have a 1:1 ratio of parental haplotypes, but a tetraploid might have a 3:1 ratio under certain circumstances.

#### 3.4. Assembly of phased accessions

The phased accessions can be assembled similar to the original accessions using HybPiper and HybPhaser (part1). The resulting sequence lists can then be used for phylogenetic analyses, ideally combined with the non-phased accessions.

### 4. Combining phased with normal sequences

The R script **Rscript_4a_combine_phased_with_normal_sequence_lists.R** can be used to combine the phased with the normal sequence lists. It is possible to make subsets of the combined dataset using text files listing accession or loci to be included or excluded.

In the configuration script **Configure_4_Combining_sequence_lists.R** all required variables can be set:

- path_to_hybpiper_results: set HybPiper base folder
- path_to_hybpiper_results_phased: set Hybpiper base folder of phased accessions
- fasta_file_with_targets: set file with sequence targets
- txt_file_with_list_of_accessions: set file with a list of normal accessions (e.g. ".../hybpiper/namelist.txt")
- txt_file_with_list_of_phased_accessions: set file with list of phased accessions (e.g. ".../hybpiper_phased/namelist_phased.txt")
- sequence_type: set sequence list name to be combined (referring to the folder name in the sequences subfolder, e.g. "consensus_loci_clean" or "contig_loci_clean")
- contig: set contig type ["normal" or "supercontig"]
- name_of_sequence_list_output: set name of the output sequence list
- file_with_samples_included: define subset by listing accession to include (“” will include all)
- file_with_samples_excluded: define subset by listing accession to exclude (“” will exclude none)
- file_with_loci_excluded: define subset by listing loci to exclude (“” will exclude none)
- exchange_phased_with_not_phased_samples: set whether the non-phased accessions of the phased samples will be exchanged. ("yes" will exclude normal accessions, "no" will include normal and phased accessions from the same sample)

The combined sequence lists are ready for alignment and phylogenetic analyses.
