## Appendix 2 - Table of accessions for "HybPhaser: a workflow for the detection and phasing of hybrids in target capture datasets"

Appendix 2 – List of accessions

Table 1. List of accessions included in this study with DNA or SRA number, short name used in manuscript, voucher information, geographic origin, and whether the sample has been used in Murphy et al. 2020 or was generated for this study.

| Species / hybrid | DNA number / SRA accession | Name in manuscript | Collector & Field number | Geographic origin | Study |
| --- | --- | --- | --- | --- | --- |
| Nepenthes adnata | ERR3662059 | NA (excluded from analyses) | Murphy 296 (K) | Kelog Sembilan, Sumatra | Murphy et al. 2020 |
| Nepenthes alata | G07767 | alata 1 | Clarke, C. & Bourke, G. 29 | Luzon | this study |
| Nepenthes alata | G07886 | alata 2 | Clarke, C. & Bourke, G. 33 | Mt. Data, Luzon | this study |
| Nepenthes alata | ERR3657191 | alata 3 | Murphy 306 (K) | Bontoc province, Luzon, Philippines | Murphy et al. 2020 |
| Nepenthes alba | ERR3657192 | alba |  | Gunung Tahan, Peninsular Malaysia | Murphy et al. 2020 |
| Nepenthes albomarginata | G08095 | albomarginata 1 | Crayn, D.M. 1632 | - | this study |
| Nepenthes albomarginata | G08129 | albomarginata 2 | Crayn, D.M. 1614 | - | this study |
| Nepenthes albomarginata | ERR3657194 | albomarginata 3 |  | Gunung Jerai, N.W. Pen. Malaysia | Murphy et al. 2020 |
| Nepenthes albomarginata | ERR3662876 | albomarginata 4 | Murphy 303 (K) | Labi Road, Brunei, Borneo | Murphy et al. 2020 |
| Nepenthes ampullaria | G07903 | ampullaria 1 | Clarke, C. & Bourke, G. 2 | Borneo | this study |
| Nepenthes ampullaria | G08093 | ampullaria 2 | Crayn, D.M. 1630 | - | this study |
| Nepenthes ampullaria | G08134 | ampullaria 3 | Crayn, D.M. 1619 | - | this study |
| Nepenthes ampullaria | G08137 | ampullaria 4 | Crayn, D.M. 1622 | - | this study |
| Nepenthes ampullaria | ERR3657200 | ampullaria 5 |  | Taiyeve, Mamberamo Regency, Papua | Murphy et al. 2020 |
| Nepenthes ampullaria | ERR3659452 | ampullaria 6 |  | Labi Road, Brunei | Murphy et al. 2020 |
| Nepenthes ampullaria | ERR3660988 | ampullaria 7 |  | Labi Road, Brunei | Murphy et al. 2020 |
| Nepenthes ampullaria | ERR3662930 | ampullaria 8 |  | Enaratoli-Nabiri, Papua, Indonesia | Murphy et al. 2020 |
| Nepenthes ampullaria × gracilis | G08101 | ampullaria × gracilis | Crayn, D.M. 1638 | - | this study |
| Nepenthes ampullaria × rafflesiana | G08104 | ampullaria × rafflesiana | Crayn, D.M. 1641 | - | this study |
| Nepenthes ampullaria × tobaica | G08098 | ampullaria × tobaica | Crayn, D.M. 1635 | - | this study |
| Nepenthes andamana | ERR3662060 | andamana | Murphy 365 (K) | Takuapa, Phang-nga, Thailand | Murphy et al. 2020 |
| Nepenthes angasanensis | ERR3712174 | angasanensis |  | Sumatra | Murphy et al. 2020 |
| Nepenthes angustifolia | ERR3712173 | angustifolia | Cheek 19060 (K) | Borneo | Murphy et al. 2020 |
| Nepenthes aristolochioides | ERR3662061 | aristolochioides |  | Jambi, Sumatra | Murphy et al. 2020 |
| Nepenthes armin | ERR3712097 | armin | Murphy 300 (K) | Sibuyan, Philippines | Murphy et al. 2020 |
| Nepenthes attenboroughii | G08117 | attenboroughii 1 | Crayn, D.M. 1602 | - | this study |
| Nepenthes attenboroughii | ERR3712098 | attenboroughii 2 |  | Palawan, Philippines | Murphy et al. 2020 |
| Nepenthes beccariana | ERR3660989 | beccariana | Cheek 17894 (K) | Sibolga, North Sumatra | Murphy et al. 2020 |
| Nepenthes bellii | G08102 | bellii 1 | Crayn, D.M. 1639 | - | this study |
| Nepenthes bellii | ERR3662062 | bellii 2 | Cheek 17884 (K) | Mindanao | Murphy et al. 2020 |
| Nepenthes benstonei | G07897 | benstonei 1 | Clarke, C. & Bourke, G. 38 | Bukit Bakar, Kelantan, Malaysia (type locality) | this study |
| Nepenthes benstonei | ERR3662142 | benstonei 2 |  | Bukit Bakar, Kelantan, Pen. Malaysia | Murphy et al. 2020 |
| Nepenthes biak | ERR3712175 | biak | Murphy 313 (K) | Biak, Papua, Indonesia | Murphy et al. 2020 |
| Nepenthes bicalcarata | G08130 | bicalcarata 1 | Crayn, D.M. 1615 | - | this study |
| Nepenthes bicalcarata | ERR3657202 | bicalcarata 2 |  | Sri Aman, Sarawak | Murphy et al. 2020 |
| Nepenthes bicalcarata | ERR3660990 | bicalcarata 3 |  | Sipitang, Sabah | Murphy et al. 2020 |
| Nepenthes bicalcarata | ERR3672060 | bicalcarata 4 | Murphy B. 325 (K) |  | Murphy et al. 2020 |
| Nepenthes bicalcarata × ampullaria | G08132 | bicalcarata × ampullaria | Crayn, D.M. 1617 | - | this study |
| Nepenthes bokorensis | G07856 | bokorensis 1 | Clarke, C. & Bourke, G. 35 | Mt. Bokor massif, Cambodia | this study |
| Nepenthes bokorensis | ERR3662143 | bokorensis 2 | Murphy 308 (K) | Bokor Falls, Cambodia | Murphy et al. 2020 |
| Nepenthes bokorensis × ventricosa | G07899 | bokorensis × ventricosa | Clarke, C. & Bourke, G. 54 | Horticultural hybrid | this study |
| Nepenthes bongso | G07898 | bongso 1 | Clarke, C. & Bourke, G. 19 | Sumatra | this study |
| Nepenthes bongso | ERR3662063 | bongso 2 | Cheek 17854 (K) | Gunung Merapi, Sumatra | Murphy et al. 2020 |
| Nepenthes borneensis | G07870 | borneensis | Clarke, C. & Bourke, G. 23 | Mt. Besar (type locality), Kalimantan Selatan | this study |
| Nepenthes boschiana | G07855 | boschiana 1 | Clarke, C. & Bourke, G. 32 | Mt Sakumpang (type locality), Kalimantan Selatan | this study |
| Nepenthes boschiana | G08112 | boschiana 2 | Crayn, D.M. 1597 | - | this study |
| Nepenthes boschiana | ERR3662144 | boschiana 4 |  | Gunung Besar, South Kalimantan | Murphy et al. 2020 |
| Nepenthes boschiana × glandulifera | G07862 | boschiana × glandulifera | Clarke, C. & Bourke, G. 52 | Horticultural hybrid | this study |
| Nepenthes burbidgeae | ERR3662064 | burbidgeae | Cheek 17827 (K) | Mt. Kinabalu, Borneo | Murphy et al. 2020 |
| Nepenthes burkei | G07875 | burkei 1 | Clarke, C. & Bourke, G. 5 | Mindoro, Philippines | this study |
| Nepenthes burkei | ERR3662065 | burkei 2 | Cheek 17851 (K) | Mt. Halcon, Mindoro, Philippines | Murphy et al. 2020 |
| Nepenthes burkei × ventricosa | G07783 | burkei × ventricosa 1 | Clarke, C. & Bourke, G. 53 | Horticultural hybrid | this study |
| Nepenthes burkei × ventricosa | G07854 | burkei × ventricosa 2 | Clarke, C. & Bourke, G. 53 | Horticultural hybrid | this study |
| Nepenthes burkei × ventricosa | G07877 | burkei × ventricosa 3 |  |  | this study |
| Nepenthes campanulata | G08136 | campanulata 1 | Crayn, D.M. 1621 | - | this study |
| Nepenthes campanulata | ERR3662145 | campanulata 2 |  | Gunung mulu, Sarawak | Murphy et al. 2020 |
| Nepenthes ceciliae | ERR3662066 | ceciliae | Murphy 212 (K) | Mt Kiamo, Mindanao, Philippines | Murphy et al. 2020 |
| Nepenthes chang | ERR3712005 | chang | Cheek 17897 (K) | Ko Chang Island | Murphy et al. 2020 |
| Nepenthes chaniana | G07882 | chaniana 1 | Clarke, C. & Bourke, G. 27 | Sarawak, Borneo | this study |
| Nepenthes chaniana | ERR3662277 | chaniana 2 | Cheek 17869 (K) | Gunung Batu Lawi, Sarawak, BE | Murphy et al. 2020 |
| Nepenthes clipeata | ERR3662278 | clipeata | Cheek 17887 (K) | Gunung Kelam, West Kalimantan | Murphy et al. 2020 |
| Nepenthes clipeata × rafflesiana | G08106 | clipeata × rafflesiana | Crayn, D.M. 1643 | Horticultural hybrid | this study |
| Nepenthes copelandii | G07863 | copelandii 1 | Clarke, C. & Bourke, G. 25 | Mt. Pasian, Mindanao | this study |
| Nepenthes copelandii | G07883 | copelandii 2 | Clarke, C. & Bourke, G. 24 | Mt. Apo, Mindanao | this study |
| Nepenthes copelandii | ERR3662279 | copelandii 3 |  | Mt Apo, Mindanao, Philippines | Murphy et al. 2020 |
| Nepenthes cornuta | ERR3712006 | cornuta |  | Mindanao, Philippines | Murphy et al. 2020 |
| Nepenthes danseri | ERR3672061 | danseri | Murphy B. 309 (K) |  | Murphy et al. 2020 |
| Nepenthes deaniana | ERR3712177 | deaniana |  | Mt Pulgar/Thumb's Peak, Palawan | Murphy et al. 2020 |
| Nepenthes densiflora | ERR3662067 | densiflora | Murphy 214 (K) | Aceh, Sumatra, Indonesia | Murphy et al. 2020 |
| Nepenthes diatas | ERR3662280 | diatas | Cheek 17888 (K) | Gunung Bandalhara, Aceh, Sumatra | Murphy et al. 2020 |
| Nepenthes distillatoria | ERR3662281 | distillatoria | Murphy 366 (K) | Sri Lanka | Murphy et al. 2020 |
| Nepenthes dubia | ERR3657203 | dubia |  | Gunung Talakamau, W. Sumatra | Murphy et al. 2020 |
| Nepenthes edwardsiana | ERR3660991 | edwardsiana |  | Mt Tambuyukon, Sabah, Borneo | Murphy et al. 2020 |
| Nepenthes ephippiata | G07857 | ephippiata 1 | Clarke, C. & Bourke, G. 37 | Bukit Batu, Hose Mtns, Sarawak. | this study |
| Nepenthes ephippiata | G07900 | ephippiata 2 | Clarke, C. & Bourke, G. 36 | Bukit Raya, Kalimantan Barat/Tengah | this study |
| Nepenthes ephippiata | ERR3660992 | ephippiata 3 | Cheek 17880 (K) | Gunung Raya, W. Kalimantan, Indonesia | Murphy et al. 2020 |
| Nepenthes eustachya | ERR3662068 | eustachya |  | Kelong Semblian, Sumatra | Murphy et al. 2020 |
| Nepenthes eymae | G07887 | eymae 1 | Clarke, C. & Bourke, G. 20 | Mt. Lumut, Sulawesi | this study |
| Nepenthes eymae | G08121 | eymae 2 | Crayn, D.M. 1606 | - | this study |
| Nepenthes eymae | ERR3662871 | eymae 3 |  | Sul Teng, Sulawesi | Murphy et al. 2020 |
| Nepenthes faizaliana | G08090 | faizaliana 1 | Crayn, D.M. 1627 | - | this study |
| Nepenthes faizaliana | ERR3662069 | faizaliana 2 |  | Gunung Mulu, Sarawak | Murphy et al. 2020 |
| Nepenthes flava | G08119 | flava 1 | Crayn, D.M. 1604 | - | this study |
| Nepenthes flava | ERR3712178 | flava 2 | Cheek 17845 (K) | Sumatra | Murphy et al. 2020 |
| Nepenthes glabrata | ERR3662070 | glabrata | Cheek 17832 (K) | Highlands, Central Sulawesi | Murphy et al. 2020 |
| Nepenthes glandulifera | G07830 | glandulifera 1 | Clarke, C. & Bourke, G. 30 | Bukit Batu, Hose Mtns, Sarawak. | this study |
| Nepenthes glandulifera | ERR3662282 | glandulifera 2 | Cheek 17862 (K) | Hose Mountains, Sarawak, Malaysia | Murphy et al. 2020 |
| Nepenthes graciliflora | ERR3661586 | graciliflora 1 | Murphy 362 (K) | Foothills Mt Guiting Guiting, Sibuyan | Murphy et al. 2020 |
| Nepenthes graciliflora | ERR3662284 | graciliflora 2 | Murphy 305 (K) |  | Murphy et al. 2020 |
| Nepenthes graciliflora | G08092 | NA (excluded from analyses) | Crayn, D.M. 1629 | - | this study |
| Nepenthes gracilis | G08091 | gracilis 1 | Crayn, D.M. 1628 | - | this study |
| Nepenthes gracilis | ERR3657205 | gracilis 2 |  | Kuche, Tak Bai, Narathiwat, S. Thailand | Murphy et al. 2020 |
| Nepenthes gracilis | ERR3662283 | gracilis 3 | Murphy 310 (K) | Labi Road, Brunei | Murphy et al. 2020 |
| Nepenthes gracillima | ERR3662285 | gracillima |  | Tahan Mnts., Pahang, Pen. Malaysia | Murphy et al. 2020 |
| Nepenthes gymnamphora | G07893 | gymnamphora 1 | Clarke, C. & Bourke, G. 12 | Sumatra | this study |
| Nepenthes gymnamphora | G07901 | gymnamphora 2 | Clarke, C. & Bourke, G. 8 | Sumatra | this study |
| Nepenthes gymnamphora | ERR3662071 | gymnamphora 3 | Cheek 17834 (K) | Mt Talakmau, Ophir District, Sumatra | Murphy et al. 2020 |
| Nepenthes gymnamphora | ERR3662923 | gymnamphora 4 | Cheek 17864 (K) | Wonosobo, Dieng Mts., C. Java | Murphy et al. 2020 |
| Nepenthes hamata | ERR3657206 | hamata 1 |  | Gunung Katopasa, Sulawesi | Murphy et al. 2020 |
| Nepenthes hamata | ERR3657207 | hamata 2 |  | Gunung Lumut, Sulawesi | Murphy et al. 2020 |
| Nepenthes hamata | G07884 | NA (excluded from analyses) | Clarke, C. & Bourke, G. 14 | Mt. Lumut, Sulawesi | this study |
| Nepenthes hamiguitanensis | ERR3657208 | hamiguitanensis |  | Mt Hamiguitan, Mindanao | Murphy et al. 2020 |
| Nepenthes hemsleyana | ERR3662286 | hemsleyana | Murphy 311 (K) | Labi Road, Brunei | Murphy et al. 2020 |
| Nepenthes hirsuta | ERR3662072 | hirsuta |  | Gunung Serapai, Sarawak | Murphy et al. 2020 |
| Nepenthes hispida | ERR3712007 | hispida | Murphy 314 (K) | Brunei | Murphy et al. 2020 |
| Nepenthes hurrelliana | G08258 | NA (excluded from analyses) | Jago, B. 1465 | - | this study |
| Nepenthes inermis | ERR3662073 | inermis | Cheek 17841 (K) | Gunung Gadut, Padang, W. Sumatra | Murphy et al. 2020 |
| Nepenthes insignis | ERR3661587 | insignis |  | Taiyeve, Mamberamo Regency, Papua | Murphy et al. 2020 |
| Nepenthes izumiae | G07891 | izumiae 1 | Clarke, C. & Bourke, G. 10 | Mt. Lesung Tungkut, Sumatra | this study |
| Nepenthes izumiae | ERR3662287 | izumiae 2 |  | Bukit barisian, West Sumatra | Murphy et al. 2020 |
| Nepenthes izumiae × ventricosa | G07866 | izumiae × ventricosa | Clarke, C. & Bourke, G. 51 | Horticultural hybrid | this study |
| Nepenthes jacquelineae | G07873 | jacquelineae 1 | Clarke, C. & Bourke, G. 9 | Mt. Lesung Tungkut, Sumatra | this study |
| Nepenthes jacquelineae | ERR3662074 | jacquelineae 2 | Cheek 17835 (K) | Gunung Gadung, W Sumatra | Murphy et al. 2020 |
| Nepenthes jamban | ERR3662075 | jamban | Cheek 17870 (K) | Bukit Barisan | Murphy et al. 2020 |
| Nepenthes justinae | ERR3712099 | justinae |  | Mt. Hamiguitan, Mindanao, Philippines | Murphy et al. 2020 |
| Nepenthes kampotiana | G07895 | kampotiana 1 | Clarke, C. & Bourke, G. 49 | Cambodia or Vietnam? | this study |
| Nepenthes kampotiana | ERR3657987 | kampotiana 2 |  | Trat Province, Thailand | Murphy et al. 2020 |
| Nepenthes kerrii | ERR3712008 | kerrii | Murphy 361 (K) | Satun Province, Thailand | Murphy et al. 2020 |
| Nepenthes khasiana | G07874 | khasiana 1 | Clarke, C. & Bourke, G. 15 | Horticulture, India | this study |
| Nepenthes khasiana | ERR3662076 | khasiana 2 | Cheek 17896 (K) | India | Murphy et al. 2020 |
| Nepenthes klossii | ERR3661588 | klossii |  | Papua | Murphy et al. 2020 |
| Nepenthes kongkandana | G07869 | kongkandana 1 | Clarke, C. & Bourke, G. 48 | Narathiwat Province, Thailand (type locality) | this study |
| Nepenthes kongkandana | ERR3662288 | kongkandana 2 | Cheek 17875 (K) | Chana, Hat Yali, Thailand | Murphy et al. 2020 |
| Nepenthes lamii | ERR3661589 | NA (excluded from analyses) |  | Mount Dorman, Puncak Regency, Papua | Murphy et al. 2020 |
| Nepenthes lavicola | ERR3662077 | NA (excluded from analyses) |  | Gunung Geureadong, Sumatra | Murphy et al. 2020 |
| Nepenthes leonardoi | ERR3712100 | leonardoi |  | Palawan, Philippines | Murphy et al. 2020 |
| Nepenthes lingulata | ERR3662078 | lingulata | Cheek 17865 (K) | Bukit Barisan, Sumatra | Murphy et al. 2020 |
| Nepenthes longifolia | ERR3662289 | longifolia | Murphy 363 (K) | Kelong Semblian, Sumatra | Murphy et al. 2020 |
| Nepenthes longifolia | G07763 | NA (excluded from analyses) | Clarke, C. & Bourke, G. 45 | Kelok Sembilan, Sumatra Barat, Indonesia | this study |
| Nepenthes lowii | ERR3662085 | lowii |  | Gunung Mulu, Sarawak, Malaysia | Murphy et al. 2020 |
| Nepenthes lowii × campanulata | G08109 | lowii × campanulata | Crayn, D.M. 1646 | - | this study |
| Nepenthes macfarlanei | ERR3662290 | macfarlanei |  | Cameron Highlands, Pen. Malaysia | Murphy et al. 2020 |
| Nepenthes macfarlanei | G08257 | NA (excluded from analyses) | Chua et al. FRI 39048 | Gunung Batu Putih, Malaysia | this study |
| Nepenthes macrophylla | ERR3662086 | macrophylla |  | Mt. Trus Madi, Sabah, Borneo | Murphy et al. 2020 |
| Nepenthes macrovulgaris | ERR3657988 | macrovulgaris |  | Gunung Silam, Sabah, Malaysia | Murphy et al. 2020 |
| Nepenthes madagascariensis | ERR3672062 | madagascariensis | Murphy B. 302 (K) |  | Murphy et al. 2020 |
| Nepenthes mantalingajanensis | ERR3712101 | mantalingajanensis |  | Mount Mantalingajan, Palawan | Murphy et al. 2020 |
| Nepenthes mapuluensis | ERR3662087 | mapuluensis |  | Sankulirang Range, East Kalimantan | Murphy et al. 2020 |
| Nepenthes masoalensis | ERR3662088 | masoalensis | Cheek 17895 (K) | Mt Ambato, Masoala Pen., Madagascar | Murphy et al. 2020 |
| Nepenthes maxima | G07859 | maxima 1 | Clarke, C. & Bourke, G. 44 | Sulawesi | this study |
| Nepenthes maxima | ERR3661608 | maxima 10 | Murphy 222 (K) | Mt. Sessian, Sulawesi | Murphy et al. 2020 |
| Nepenthes maxima | ERR3662875 | maxima 11 | Cheek 17879 (K) | Central Sulawesi | Murphy et al. 2020 |
| Nepenthes maxima | G07861 | maxima 2 | Clarke, C. & Bourke, G. 43 | Sulawesi, lowlands | this study |
| Nepenthes maxima | G07864 | maxima 3 | Clarke, C. & Bourke, G. 21 | Mt. Sesean, Sulawesi | this study |
| Nepenthes maxima | G08103 | maxima 4 | Crayn, D.M. 1640 | - | this study |
| Nepenthes maxima | G08107 | maxima 5 | Crayn, D.M. 1644 | - | this study |
| Nepenthes maxima | ERR3661602 | maxima 6 | Cheek 17842 (K) | Maluku, Sulawesi, 1600m | Murphy et al. 2020 |
| Nepenthes maxima | ERR3661604 | maxima 7 |  | Napu valley, 350m, Sulawesi | Murphy et al. 2020 |
| Nepenthes maxima | ERR3661605 | maxima 8 | Murphy 1003 (K) | Anggi, Manokwari, W. Papua | Murphy et al. 2020 |
| Nepenthes maxima | ERR3661607 | maxima 9 | Cheek 17891 (K) | Mt.Tinombala road, Sulawesi | Murphy et al. 2020 |
| Nepenthes maxima | ERR3661606 | NA (excluded from analyses) | Murphy 1015 (K) | Sopnyai, Hingk, Manokwari, W.Papua | Murphy et al. 2020 |
| Nepenthes maxima | G07894 | NA (excluded from analyses) | Clarke, C. & Bourke, G. 40 | New Guinea highlands (Papua, Indonesia) | this study |
| Nepenthes merrilliana | G08089 | merrilliana 1 | Crayn, D.M. 1626 | - | this study |
| Nepenthes merrilliana | ERR3662291 | merrilliana 2 | Murphy 312 (K) | Foothills of Mount Legaspi, Mindanao | Murphy et al. 2020 |
| Nepenthes micramphora | ERR3657989 | micramphora |  | Mount Hamiguitan, Mindanao | Murphy et al. 2020 |
| Nepenthes mikei | ERR3662089 | mikei | Cheek 17857 (K) | Mt.Bandahara, Aceh, Sumatra | Murphy et al. 2020 |
| Nepenthes mindanaoensis | ERR3661609 | mindanaoensis |  | Dinagat | Murphy et al. 2020 |
| Nepenthes minima | G07872 | minima 1 | Clarke, C. & Bourke, G. 50 | Lake Poso, Sulawesi | this study |
| Nepenthes minima | ERR3662292 | minima 2 |  | Lake Poso area, Sulteng., Sulawesi | Murphy et al. 2020 |
| Nepenthes mira | G07902 | mira 1 | Clarke, C. & Bourke, G. 4 | Cleopatra's Needle, Palawan, Philippines (type locality) | this study |
| Nepenthes mira | ERR3662090 | mira 2 | Cheek 17824 (K) | Palawan, Philippines | Murphy et al. 2020 |
| Nepenthes mirabilis | G08086 | mirabilis 1 | Crayn, D.M. 1623 | - | this study |
| Nepenthes mirabilis | ERR3657198 | mirabilis 10 |  | Trat, Thailand | Murphy et al. 2020 |
| Nepenthes mirabilis | ERR3657199 | mirabilis 11 | Trethowan 527 (K) | Wawonii, South East Sulawesi | Murphy et al. 2020 |
| Nepenthes mirabilis | ERR3661610 | mirabilis 12 |  | South of Samarinda, East Kalimantan | Murphy et al. 2020 |
| Nepenthes mirabilis | ERR3661611 | mirabilis 13 |  | Gunung Lampia, Sulawesi | Murphy et al. 2020 |
| Nepenthes mirabilis | ERR3661613 | mirabilis 14 | Murphy 1054 (K) | Gunung Botak, Manokwari, W. Papua | Murphy et al. 2020 |
| Nepenthes mirabilis | ERR3661615 | mirabilis 15 | Murhpy 1048 (K) | Gunung Botak, Manokwari, W. Papua | Murphy et al. 2020 |
| Nepenthes mirabilis | ERR3661616 | mirabilis 16 | Murphy 1051 (K) | Gunung Botak, Manokwari, W. Papua | Murphy et al. 2020 |
| Nepenthes mirabilis | ERR3661617 | mirabilis 17 | Murphy 1044 (K) | Gunung Botak, Manokwari, W. Papua | Murphy et al. 2020 |
| Nepenthes mirabilis | ERR3661618 | mirabilis 18 | Murphy 1053 (K) | Gunung Botak, Manokwari, W. Papua | Murphy et al. 2020 |
| Nepenthes mirabilis | ERR3661619 | mirabilis 19 | Murphy 1055 (K) | Gunung Botak, Manokwari, W. Papua | Murphy et al. 2020 |
| Nepenthes mirabilis | G08094 | mirabilis 2 | Crayn, D.M. 1631 | - | this study |
| Nepenthes mirabilis | ERR3661620 | mirabilis 20 | Murhpy 1050 (K) | Gunung Botak, Manokwari, W. Papua | Murphy et al. 2020 |
| Nepenthes mirabilis | ERR3661621 | mirabilis 21 | Murphy 1059 (K) | Gunung Botak, Manokwari, W. Papua | Murphy et al. 2020 |
| Nepenthes mirabilis | ERR3661622 | mirabilis 22 | Murphy 1059 (K) | Gunung Botak, Manokwari, W. Papua | Murphy et al. 2020 |
| Nepenthes mirabilis | ERR3662877 | mirabilis 23 | Murphy 304 (K) | Pasir Panjang, W. Kalimantan | Murphy et al. 2020 |
| Nepenthes mirabilis | ERR3662928 | mirabilis 24 | Murphy 1045 (K) | Gunung Botak, Manokwari, W. Papua | Murphy et al. 2020 |
| Nepenthes mirabilis | ERR3662929 | mirabilis 25 | Murphy 364 (K) | Trang, Thailand | Murphy et al. 2020 |
| Nepenthes mirabilis | ERR3672063 | mirabilis 26 | Murphy B. 324 (K) |  | Murphy et al. 2020 |
| Nepenthes mirabilis | G08099 | mirabilis 3 | Crayn, D.M. 1636 | - | this study |
| Nepenthes mirabilis | G08236 | mirabilis 4 | Wilson, G.W. 652 | Cape York, Jardine River NP | this study |
| Nepenthes mirabilis | G08237 | mirabilis 5 | Wilson, G.W. 660 | Cape York, Jardine River NP | this study |
| Nepenthes mirabilis | G08244 | mirabilis 6 | Wilson, G.W. 734 | Cape York, Heathlands NP | this study |
| Nepenthes mirabilis | G08245 | mirabilis 7 | Wilson, G.W. 737 | Cape York, Steve Irwin Scientific Reserve | this study |
| Nepenthes mirabilis | G08246 | mirabilis 8 | Wilson, G.W. 741 | Bramston Beach, QLD | this study |
| Nepenthes mirabilis | G08247 | mirabilis 9 | Wilson, G.W. 744 | Bramston Beach, QLD | this study |
| Nepenthes mirabilis | G08131 | mirabilis var echinostoma 1 | Crayn, D.M. 1616 | - | this study |
| Nepenthes mirabilis | ERR3657990 | NA (excluded from analyses) | Merello 3295 (K) | Halmahera, Maluku | Murphy et al. 2020 |
| Nepenthes mirabilis | ERR3657991 | NA (excluded from analyses) | Trethowan 214 (K) | Morowali, Central Sulawes | Murphy et al. 2020 |
| Nepenthes mirabilis | ERR3661612 | NA (excluded from analyses) | Murphy 1052 (K) | Gunung Botak, Manokwari, W. Papua | Murphy et al. 2020 |
| Nepenthes mirabilis | ERR3661614 | NA (excluded from analyses) | Murphy 1043 (K) | Gunung Botak, Manokwari, W. Papua | Murphy et al. 2020 |
| Nepenthes mirabilis var. echinostoma | ERR3657204 | mirabilis var echinostoma 2 |  | Lambi Hills, Sarawak | Murphy et al. 2020 |
| Nepenthes mirabilis var. globosa | G08128 | mirabilis var globosa | Crayn, D.M. 1612 | - | this study |
| Nepenthes mollis | ERR3712179 | mollis |  | Borneo | Murphy et al. 2020 |
| Nepenthes monticola | ERR3712102 | monticola | Willis 147 (K) | Freeport, Mimika Regency, Papua | Murphy et al. 2020 |
| Nepenthes muluensis | ERR3657371 | muluensis |  | Gunung Murud, Sarawak, Borneo | Murphy et al. 2020 |
| Nepenthes murudensis | ERR3662293 | murudensis |  | Gunung Murud, Sarawak, Borneo | Murphy et al. 2020 |
| Nepenthes naga | ERR3662294 | naga | Cheek 17860 (K) | Bukit Bansan, Sumatra | Murphy et al. 2020 |
| Nepenthes neoguineensis | G08125 | neoguineensis 1 | Crayn, D.M. 1610 | - | this study |
| Nepenthes neoguineensis | ERR3657378 | neoguineensis 2 | Argent 521 (K) | Freeport, Mimika Regency, Papua | Murphy et al. 2020 |
| Nepenthes neoguineensis | ERR3662053 | neoguineensis 3 |  | Jayapura, Papua | Murphy et al. 2020 |
| Nepenthes northiana | G08135 | northiana 1 | Crayn, D.M. 1620 | - | this study |
| Nepenthes northiana | ERR3662295 | northiana 2 |  | Near Bau, S. Kuching, Sarawak | Murphy et al. 2020 |
| Nepenthes oblanceolata | ERR3712103 | oblanceolata |  | Wamena area, Papua, Indonesia | Murphy et al. 2020 |
| Nepenthes ovata | G08118 | ovata 1 | Crayn, D.M. 1603 | - | this study |
| Nepenthes ovata | ERR3662296 | ovata 2 | Cheek 17830 (K) | Gunung Pangulubao, Sumatra | Murphy et al. 2020 |
| Nepenthes palawanensis | ERR3712010 | palawanensis |  | Sultan Peak, Palawan, Philippines | Murphy et al. 2020 |
| Nepenthes papuana | ERR3657379 | papuana |  | Mt. Doorman, Puncak Regency, Papua | Murphy et al. 2020 |
| Nepenthes parvula | G08240 | parvula 1 | Wilson, G.W. 723 | Cape York, South Jardine Swamp | this study |
| Nepenthes parvula | G08248 | parvula 2 | Wilson, G.W. 746 | Cape York, Jardine River NP | this study |
| Nepenthes peltata | ERR3662297 | peltata | Cheek 17849 (K) | Mount Hamiguitan, Mindanao | Murphy et al. 2020 |
| Nepenthes pervillei | G08123 | pervillei 1 | Crayn, D.M. 1608 | - | this study |
| Nepenthes pervillei | ERR3657380 | pervillei 2 |  | Seychelles | Murphy et al. 2020 |
| Nepenthes petiolata | G07881 | petiolata 1 | Clarke, C. & Bourke, G. 22 | Mt. Urdaneta, Mindanao, Philippines | this study |
| Nepenthes petiolata | ERR3662091 | petiolata 2 | Murphy 320 (K) | Mount Hilong-Hilong, Mindanao | Murphy et al. 2020 |
| Nepenthes philippinensis | G07878 | philippinensis 1 | Clarke, C. & Bourke, G. 1 | Mt. Victoria, Palawan | this study |
| Nepenthes philippinensis | ERR3662299 | philippinensis 2 | Cheek 17872 (K) | Palawan | Murphy et al. 2020 |
| Nepenthes pitopangii | ERR3712011 | pitopangii |  | Wistuba | Murphy et al. 2020 |
| Nepenthes platychila | ERR3662300 | platychila | Cheek 17873 (K) | Bukit Batu, Hose Mountains, Sarawak | Murphy et al. 2020 |
| Nepenthes pulchra | ERR3657381 | pulchra |  | Mindanao, Philippines | Murphy et al. 2020 |
| Nepenthes rafflesiana | G08097 | rafflesiana 1 | Crayn, D.M. 1634 | - | this study |
| Nepenthes rafflesiana | G08127 | rafflesiana 2 | Crayn, D.M. 1613 | - | this study |
| Nepenthes rafflesiana | ERR3657382 | rafflesiana 3 |  | Kuching, Sarawak | Murphy et al. 2020 |
| Nepenthes rafflesiana | ERR3657383 | rafflesiana 4 |  | Jahore Bahru, Peninsular Malaysia | Murphy et al. 2020 |
| Nepenthes rafflesiana | ERR3662092 | rafflesiana 5 |  | Mt Kinabalu, Sabah | Murphy et al. 2020 |
| Nepenthes rafflesiana × ampullaria | G08126 | rafflesiana × ampullaria 1 | Crayn, D.M. 1611 | - | this study |
| Nepenthes rafflesiana × ampullaria (× hookeriana) | ERR3662102 | rafflesiana × ampullaria 2 |  | Labi Road, Brunei | Murphy et al. 2020 |
| Nepenthes rajah | G08114 | rajah 1 | Crayn, D.M. 1599 | - | this study |
| Nepenthes rajah | ERR3657984 | rajah 2 |  | Gunung Tamboyokan, Kinabalu, Sabah | Murphy et al. 2020 |
| Nepenthes ramispina | ERR3662301 | ramispina | Cheek 17859 (K) | Gunung Ulu Kali, Pen. Malaysia | Murphy et al. 2020 |
| Nepenthes reinwardtiana | G08100 | reinwardtiana 1 | Crayn, D.M. 1637 | - | this study |
| Nepenthes reinwardtiana | G08133 | reinwardtiana 2 | Crayn, D.M. 1618 | - | this study |
| Nepenthes reinwardtiana | ERR3662093 | reinwardtiana 3 | Murphy 319 (K) | Tambunan Road, Crocker Range, Sabah | Murphy et al. 2020 |
| Nepenthes rhombicaulis | ERR3662302 | rhombicaulis | Cheek 17831 (K) | Gunung Pangulabao, Sumatra | Murphy et al. 2020 |
| Nepenthes rigidifolia | ERR3712180 | rigidifolia |  | Mt. Sidikalang, Karo Regncy, Sumatra | Murphy et al. 2020 |
| Nepenthes robcantleyi | G08111 | robcantleyi 1 | Crayn, D.M. 1596 | - | this study |
| Nepenthes robcantleyi | ERR3712181 | robcantleyi 2 |  | Mindanao | Murphy et al. 2020 |
| Nepenthes rowaniae | G08241 | rowaniae 6 | Wilson, G.W. 725 | Cape York, East Jardine Swamp | this study |
| Nepenthes rowaniae | G08259 | NA (excluded from analyses) | Field, A.R. living collection | - | this study |
| Nepenthes rowaniae | G05999 | rowaniae 1 | Schulte, K. 259 |  | this study |
| Nepenthes rowaniae | G08233 | rowaniae 2 | Gray, B. 9126 | Cape York, Shelburne, QLD | this study |
| Nepenthes rowaniae | G08234 | rowaniae 3 | Jensen, R. 3365 | Cape York, Jardine River NP | this study |
| Nepenthes rowaniae | G08239 | rowaniae 4 | Wilson, G.W. 666 | Cape York, South Jardine Swamp | this study |
| Nepenthes rowaniae | G08249 | rowaniae 5 | Wilson, G.W. 747 | Cape York, Jardine River NP | this study |
| Nepenthes rowaniae | ERR3662094 | rowaniae 6 | Cheek 17886 (K) | Cape York Pen., Queensland, Australia | Murphy et al. 2020 |
| Nepenthes sanguinea | G07858 | sanguinea 1 | Clarke, C. & Bourke, G. 26 | Peninsular Malaysia | this study |
| Nepenthes sanguinea | ERR3662095 | sanguinea 2 | Murphy 317 (K) | Cameron Highlands, Pen. Malaysia | Murphy et al. 2020 |
| Nepenthes sibuyanensis | G08113 | sibuyanensis 1 | Crayn, D.M. 1598 | - | this study |
| Nepenthes sibuyanensis | ERR3662096 | sibuyanensis 2 | Murphy 316 (K) | Mount Guiting-Guiting, Sibuyan Island | Murphy et al. 2020 |
| Nepenthes singalana | G07880 | singalana 1 | Clarke, C. & Bourke, G. 7 | Sumatra | this study |
| Nepenthes singalana | ERR3662303 | singalana 2 |  | Gunung Belirang, Sumatra | Murphy et al. 2020 |
| Nepenthes smilesii | ERR3662924 | smilesii 1 |  | Kiriom, Cambodia | Murphy et al. 2020 |
| Nepenthes smilesii | ERR3662925 | smilesii 2 |  | Phu Kradung, Loei, Thailand | Murphy et al. 2020 |
| Nepenthes sp. 'adrianii' | ERR3712004 | sp adrianii | Cheek 17867 (K) | Gunung Salak | Murphy et al. 2020 |
| Nepenthes sp. Anipahan | ERR3712104 | sp Anipaha |  | Mt. Anipahan, Palawan, Philippines | Murphy et al. 2020 |
| Nepenthes sp. BM-2019 | ERR3712119 | sp BM 209 | Murphy 1089 (K) | Sumatra | Murphy et al. 2020 |
| Nepenthes sp. calcicola | G08235 | calcicola | Venter, S. 14170 | Kikori, Papua New Guinea | this study |
| Nepenthes spathulata | G07896 | spathulata 1 | Clarke, C. & Bourke, G. 28 | Mt. Tanggamus, Lampung, Sumatra | this study |
| Nepenthes spathulata | G08116 | spathulata 2 | Crayn, D.M. 1601 | - | this study |
| Nepenthes spathulata | ERR3662304 | spathulata 3 | Cheek 17829 (K) | Sumatra | Murphy et al. 2020 |
| Nepenthes spectabilis | ERR3662055 | NA (excluded from analyses) | Cheek 17848 (K) | Gunung Pangulubao, N. Sumatra | Murphy et al. 2020 |
| Nepenthes spectabilis | G07890 | spectabilis 1 | Clarke, C. & Bourke, G. 16 | North Sumatra | this study |
| Nepenthes spectabilis | ERR3657985 | spectabilis 2 | Cheek 17881 (K) | Gunung Bandahara, Sumatra | Murphy et al. 2020 |
| Nepenthes stenophylla | G08115 | stenophylla 1 | Crayn, D.M. 1600 | - | this study |
| Nepenthes stenophylla | ERR3662305 | stenophylla 2 |  | Bareo, Sarawak | Murphy et al. 2020 |
| Nepenthes sumagaya | ERR3712012 | sumagaya |  | Mount Sumagaya, Mindanao | Murphy et al. 2020 |
| Nepenthes suratensis | ERR3712013 | suratensis | Cheek 17899 (K) | Suratthani Province, Thailand | Murphy et al. 2020 |
| Nepenthes surigaoensis | ERR3662315 | surigaoensis |  | Mt Masay Elliot, Mindanao, Philippines | Murphy et al. 2020 |
| Nepenthes talangensis | G07888 | talangensis 1 | Clarke, C. & Bourke, G. 11 | Mt. Talang, Sumatra | this study |
| Nepenthes talangensis | ERR3662326 | talangensis 2 | Cheek 17861 (K) | Gunung Talang, Sumatra | Murphy et al. 2020 |
| Nepenthes tenax | G08243 | tenax 4 | Wilson, G.W. 732 | Cape York, South Jardine Swamp | this study |
| Nepenthes tenax | G08242 | tenax 5 | Wilson, G.W. 729 | Cape York, South Jardine Swamp | this study |
| Nepenthes tenax | G08087 | NA (excluded from analyses) | Crayn, D.M. 1624 | - | this study |
| Nepenthes tenax | G06000 | tenax 1 | Schulte, K. 260 |  | this study |
| Nepenthes tenax | G08238 | tenax 2 | Wilson, G.W. 661 | Cape York, Jardine River NP | this study |
| Nepenthes tenax | ERR3662327 | tenax 3 | Cheek 17866 (K) | Jardine Swamp, Cape York, Australia | Murphy et al. 2020 |
| Nepenthes tentaculata | G07885 | tentaculata 1 | Clarke, C. & Bourke, G. 17 | Mt. Kinabalu, Sabah | this study |
| Nepenthes tentaculata | ERR3662056 | tentaculata 2 |  | Gunung Raja, South Kalimantan | Murphy et al. 2020 |
| Nepenthes tentaculata | ERR3662328 | tentaculata 3 | Cheek 17844 (K) | Gunung Murud, Sarawa, Malaysia | Murphy et al. 2020 |
| Nepenthes tenuis | ERR3662329 | tenuis | Cheek 17863 (K) | Mt.Taram,Tjampo River, W.Sumatra | Murphy et al. 2020 |
| Nepenthes thai | G07879 | thai 1 | Clarke, C. & Bourke, G. 39 | Narathiwat Province, Thailand (type locality) | this study |
| Nepenthes thai | ERR3657986 | thai 2 | Cheek 17874 (K) | Kao Aidang, Hala Bala N.P, Thailand | Murphy et al. 2020 |
| Nepenthes thorelii | ERR3662097 | thorelii |  | Bin Chau Vietnam “Mr Son” | Murphy et al. 2020 |
| Nepenthes tobaica | G07871 | tobaica 1 | Clarke, C. & Bourke, G. 34 | Lake Toba, North Sumatra | this study |
| Nepenthes tobaica | G08122 | tobaica 2 | Crayn, D.M. 1607 | - | this study |
| Nepenthes tobaica | ERR3662098 | tobaica 3 | Murphy 321 (K) | Lake Toba, Sumatra | Murphy et al. 2020 |
| Nepenthes tobaica | ERR3662099 | tobaica 4 | Cheek 17833 (K) | Doloksangul | Murphy et al. 2020 |
| Nepenthes tomoriana | ERR3662330 | tomoriana |  | Gunung Lampia, Tomoroi bay, Sulawesi | Murphy et al. 2020 |
| Nepenthes treubiana | ERR3657197 | treubiana 1 |  | Raja Amput Regency, W. Papua | Murphy et al. 2020 |
| Nepenthes treubiana | ERR3662100 | treubiana 2 | Cheek 17871 (K) | Fakfak, Papua | Murphy et al. 2020 |
| Nepenthes truncata | G07889 | NA (excluded from analyses) | Clarke, C. & Bourke, G. 41 | Surigao del Norte, Mindanao | this study |
| Nepenthes truncata | G08124 | truncata 1 | Crayn, D.M. 1609 | NE Mindanao | this study |
| Nepenthes truncata | ERR3662057 | truncata 2 |  | Gunung Pasian, Mindanao | Murphy et al. 2020 |
| Nepenthes truncata × ventricosa | G08105 | truncata × ventricosa | Crayn, D.M. 1642 | Horticultural hybrid | this study |
| Nepenthes veitchii | G08088 | veitchii 1 | Crayn, D.M. 1625 |  | this study |
| Nepenthes veitchii | G08096 | veitchii 2 | Crayn, D.M. 1633 |  | this study |
| Nepenthes veitchii | ERR3662101 | veitchii 3 | Murphy 323 (K) | Bario Highlands, Sarawak | Murphy et al. 2020 |
| Nepenthes veitchii × eymae | G08108 | veitchii × eymae | Crayn, D.M. 1645 | Horticultural hybrid | this study |
| Nepenthes ventricosa | G07860 | ventricosa 1 | Clarke, C. & Bourke, G. 46 | Luzon | this study |
| Nepenthes ventricosa | G07865 | ventricosa 2 | Clarke, C. & Bourke, G. 47 | Luzon | this study |
| Nepenthes ventricosa | ERR3672064 | ventricosa 3 | Murphy B. 323 (K) |  | Murphy et al. 2020 |
| Nepenthes vieillardii | ERR3662926 | NA (excluded from analyses) | Baumann 15241 (K) | New Caledonia | Murphy et al. 2020 |
| Nepenthes vieillardii | ERR3662927 | NA (excluded from analyses) | Buckholz 1463 (K) | New Caledonia | Murphy et al. 2020 |
| Nepenthes villosa | ERR3662058 | villosa |  | Mount Kinabalu | Murphy et al. 2020 |
| Nepenthes vogelii | G08110 | vogelii 1 | Crayn, D.M. 1595 |  | this study |
| Nepenthes vogelii | ERR3672065 | vogelii 2 | Murphy B. 322 (K) |  | Murphy et al. 2020 |
| Nepenthes xiphioides | G07876 | xiphioides 1 | Clarke, C. & Bourke, G. 13 | Mt. Sinabung, Sumatra (type locality) | this study |
| Nepenthes xiphioides | ERR3662331 | xiphioides 2 | Cheek 17868 (K) | Toba Region | Murphy et al. 2020 |
| Nepenthes zakriana | G08120 | zakriana | Crayn, D.M. 1605 | - | this study |
