## Appendix 3 - HybPiper results for "HybPhaser: a workflow for the detection and phasing of hybrids in target capture datasets"

Table 1. Statistics retrieved from HybPiper script “hybpiper_stats.py” for the normal contigs (not the ones including intron regions).

| Name | Num Reads | Reads Mapped | Pct On Target | Genes Mapped | Genes With Contigs | Genes With Seqs | Genes At 25 pct | Genes At 50 pct | Genes At 75 pct | Genes at 150 pct | Paralog Warnings |
| --- | --- | --- | --- | --- | --- | --- | --- | --- | --- | --- | --- |
| Nepenthes_ rowaniae_G08241 | 1892702 | 135509 | 0.072 | 346 | 315 | 294 | 255 | 171 | 80 | 0 | 0 |
| Nepenthes_ tenax_G08242 | 3737193 | 248125 | 0.066 | 348 | 331 | 318 | 288 | 214 | 121 | 0 | 0 |
| Nepenthes_ tenax_G08243 | 3164816 | 194157 | 0.061 | 352 | 326 | 310 | 280 | 210 | 114 | 0 | 0 |
| Nepenthes_alata_ERR3657191 | 277369 | 26967 | 0.097 | 336 | 203 | 186 | 154 | 100 | 52 | 0 | 0 |
| Nepenthes_alata_G07767 | 2969360 | 236425 | 0.08 | 344 | 321 | 305 | 267 | 189 | 107 | 0 | 0 |
| Nepenthes_alata_G07886 | 2134048 | 175630 | 0.082 | 345 | 316 | 301 | 254 | 171 | 86 | 0 | 0 |
| Nepenthes_alba_ERR3657192 | 1544007 | 183056 | 0.119 | 349 | 316 | 305 | 279 | 219 | 130 | 0 | 1 |
| Nepenthes_albomarginata_ERR3657194 | 311674 | 38691 | 0.124 | 340 | 260 | 249 | 216 | 133 | 68 | 0 | 0 |
| Nepenthes_albomarginata_ERR3662876 | 994758 | 109170 | 0.11 | 348 | 304 | 293 | 263 | 199 | 108 | 0 | 0 |
| Nepenthes_albomarginata_G08095 | 3900045 | 240518 | 0.062 | 347 | 308 | 289 | 259 | 181 | 91 | 0 | 0 |
| Nepenthes_albomarginata_G08129 | 4706910 | 249938 | 0.053 | 349 | 325 | 312 | 286 | 213 | 120 | 0 | 0 |
| Nepenthes_ampullaria_ERR3657200 | 527238 | 57774 | 0.11 | 349 | 290 | 284 | 240 | 169 | 90 | 0 | 0 |
| Nepenthes_ampullaria_ERR3659452 | 1126364 | 138836 | 0.123 | 350 | 312 | 299 | 262 | 191 | 109 | 0 | 0 |
| Nepenthes_ampullaria_ERR3660988 | 1182641 | 84219 | 0.071 | 350 | 282 | 262 | 232 | 160 | 88 | 0 | 0 |
| Nepenthes_ampullaria_ERR3662930 | 475967 | 54538 | 0.115 | 347 | 285 | 275 | 239 | 166 | 87 | 0 | 0 |
| Nepenthes_ampullaria_G07903 | 4485673 | 427943 | 0.095 | 350 | 333 | 317 | 292 | 218 | 119 | 0 | 0 |
| Nepenthes_ampullaria_G08093 | 4616350 | 467338 | 0.101 | 347 | 332 | 314 | 281 | 203 | 118 | 0 | 0 |
| Nepenthes_ampullaria_G08134 | 3394505 | 279999 | 0.082 | 348 | 326 | 309 | 274 | 200 | 110 | 0 | 0 |
| Nepenthes_ampullaria_G08137 | 2439992 | 203438 | 0.083 | 347 | 321 | 302 | 266 | 192 | 95 | 0 | 0 |
| Nepenthes_ampullaria_x_gracilis_G08101 | 4524136 | 279892 | 0.062 | 348 | 314 | 300 | 270 | 183 | 95 | 0 | 0 |
| Nepenthes_ampullaria_x_rafflesiana_G08104 | 4852496 | 283109 | 0.058 | 352 | 316 | 303 | 264 | 191 | 107 | 0 | 0 |
| Nepenthes_ampullaria_x_tobaica_G08098 | 2340333 | 136964 | 0.059 | 346 | 302 | 281 | 235 | 146 | 76 | 0 | 0 |
| Nepenthes_andamana_ERR3662060 | 1070189 | 205050 | 0.192 | 343 | 317 | 301 | 272 | 204 | 113 | 0 | 0 |
| Nepenthes_angasanensis_ERR3712174 | 734344 | 128750 | 0.175 | 349 | 315 | 304 | 276 | 206 | 114 | 0 | 0 |
| Nepenthes_angustifolia_ERR3712173 | 9813935 | 404190 | 0.041 | 349 | 328 | 312 | 278 | 215 | 115 | 0 | 0 |
| Nepenthes_aristolochioides_ERR3662061 | 601658 | 108097 | 0.18 | 344 | 302 | 287 | 251 | 179 | 90 | 0 | 0 |
| Nepenthes_armin_ERR3712097 | 357674 | 42880 | 0.12 | 341 | 267 | 259 | 224 | 142 | 73 | 0 | 0 |
| Nepenthes_attenboroughii_ERR3712098 | 822525 | 85233 | 0.104 | 347 | 302 | 285 | 250 | 176 | 104 | 0 | 0 |
| Nepenthes_attenboroughii_G08117 | 3707598 | 288994 | 0.078 | 350 | 324 | 307 | 276 | 209 | 119 | 0 | 0 |
| Nepenthes_beccariana_ERR3660989 | 596268 | 59750 | 0.1 | 346 | 246 | 231 | 199 | 138 | 74 | 0 | 0 |
| Nepenthes_bellii_ERR3662062 | 644082 | 114330 | 0.178 | 341 | 297 | 287 | 251 | 183 | 104 | 0 | 0 |
| Nepenthes_bellii_G08102 | 1858762 | 110685 | 0.06 | 349 | 298 | 280 | 238 | 138 | 65 | 0 | 0 |
| Nepenthes_benstonei_ERR3662142 | 1638435 | 290141 | 0.177 | 353 | 330 | 315 | 286 | 223 | 132 | 0 | 0 |
| Nepenthes_benstonei_G07897 | 5192566 | 393868 | 0.076 | 353 | 325 | 310 | 277 | 210 | 113 | 0 | 0 |
| Nepenthes_biak_ERR3712175 | 340039 | 67646 | 0.199 | 347 | 293 | 283 | 248 | 181 | 89 | 0 | 0 |
| Nepenthes_bicalcarata_ERR3657202 | 1754376 | 131010 | 0.075 | 352 | 311 | 299 | 265 | 183 | 102 | 0 | 0 |
| Nepenthes_bicalcarata_ERR3660990 | 795063 | 44887 | 0.056 | 345 | 250 | 241 | 210 | 130 | 59 | 0 | 0 |
| Nepenthes_bicalcarata_ERR3672060 | 183008 | 25043 | 0.137 | 340 | 216 | 211 | 182 | 99 | 41 | 0 | 0 |
| Nepenthes_bicalcarata_G08130 | 2003851 | 101223 | 0.051 | 331 | 290 | 271 | 215 | 112 | 52 | 0 | 0 |
| Nepenthes_bicalcarata_x_ampullaria_G08132 | 1835190 | 116542 | 0.064 | 332 | 304 | 283 | 228 | 125 | 64 | 0 | 0 |
| Nepenthes_bokorensis_ERR3662143 | 761680 | 133151 | 0.175 | 348 | 310 | 297 | 265 | 199 | 111 | 0 | 0 |
| Nepenthes_bokorensis_G07856 | 2941174 | 128122 | 0.044 | 347 | 291 | 264 | 221 | 132 | 74 | 0 | 0 |
| Nepenthes_bokorensis_x_ventricosa_G07899 | 3127797 | 308393 | 0.099 | 348 | 324 | 311 | 271 | 191 | 103 | 0 | 0 |
| Nepenthes_bongso_ERR3662063 | 740513 | 128803 | 0.174 | 343 | 296 | 284 | 255 | 191 | 108 | 0 | 0 |
| Nepenthes_bongso_G07898 | 2319507 | 152144 | 0.066 | 333 | 308 | 286 | 244 | 153 | 75 | 0 | 0 |
| Nepenthes_borneensis_G07870 | 3919147 | 355303 | 0.091 | 344 | 324 | 307 | 280 | 198 | 112 | 0 | 0 |
| Nepenthes_boschiana_ERR3662144 | 511979 | 94959 | 0.185 | 345 | 309 | 300 | 273 | 204 | 107 | 0 | 0 |
| Nepenthes_boschiana_G07855 | 4815323 | 481911 | 0.1 | 345 | 330 | 314 | 280 | 208 | 120 | 0 | 0 |
| Nepenthes_boschiana_G08112 | 4032884 | 168753 | 0.042 | 347 | 320 | 306 | 277 | 193 | 100 | 0 | 0 |
| Nepenthes_boschiana_x_glandulifera_G07862 | 1327785 | 39617 | 0.03 | 335 | 250 | 228 | 170 | 69 | 20 | 0 | 0 |
| Nepenthes_burbidgeae_ERR3662064 | 982312 | 177814 | 0.181 | 345 | 312 | 302 | 270 | 209 | 122 | 0 | 1 |
| Nepenthes_burkei_ERR3662065 | 641187 | 101091 | 0.158 | 345 | 295 | 282 | 259 | 194 | 107 | 0 | 0 |
| Nepenthes_burkei_G07875 | 1820162 | 73711 | 0.04 | 349 | 295 | 276 | 237 | 151 | 67 | 0 | 0 |
| Nepenthes_burkei_x_ventricosa_G07783 | 3435735 | 291192 | 0.085 | 347 | 326 | 309 | 268 | 193 | 95 | 0 | 0 |
| Nepenthes_burkei_x_ventricosa_G07854 | 3007822 | 259767 | 0.086 | 346 | 324 | 304 | 270 | 187 | 88 | 0 | 0 |
| Nepenthes_burkei_x_ventricosa_G07877 | 2501786 | 118560 | 0.047 | 347 | 309 | 291 | 253 | 173 | 90 | 0 | 0 |
| Nepenthes_campanulata_ERR3662145 | 719131 | 122073 | 0.17 | 346 | 316 | 303 | 274 | 200 | 113 | 0 | 0 |
| Nepenthes_campanulata_G08136 | 2115008 | 135332 | 0.064 | 343 | 311 | 295 | 258 | 164 | 89 | 0 | 0 |
| Nepenthes_ceciliae_ERR3662066 | 936129 | 195321 | 0.209 | 346 | 313 | 304 | 271 | 203 | 119 | 0 | 0 |
| Nepenthes_chang_ERR3712005 | 845716 | 164212 | 0.194 | 343 | 305 | 294 | 263 | 196 | 109 | 0 | 0 |
| Nepenthes_chaniana_ERR3662277 | 809479 | 132422 | 0.164 | 345 | 313 | 299 | 270 | 193 | 112 | 0 | 0 |
| Nepenthes_chaniana_G07882 | 3484160 | 144954 | 0.042 | 350 | 317 | 303 | 273 | 199 | 103 | 0 | 0 |
| Nepenthes_clipeata_ERR3662278 | 452777 | 71166 | 0.157 | 341 | 293 | 284 | 252 | 182 | 96 | 0 | 0 |
| Nepenthes_clipeata_x_ventricosa_G08106 | 3040797 | 213238 | 0.07 | 347 | 317 | 303 | 272 | 190 | 100 | 0 | 0 |
| Nepenthes_copelandii_ERR3662279 | 604438 | 106978 | 0.177 | 344 | 308 | 300 | 277 | 212 | 110 | 0 | 0 |
| Nepenthes_copelandii_G07863 | 1409632 | 111249 | 0.079 | 336 | 287 | 263 | 211 | 122 | 63 | 0 | 1 |
| Nepenthes_copelandii_G07883 | 3634839 | 182731 | 0.05 | 352 | 326 | 304 | 255 | 176 | 90 | 0 | 0 |
| Nepenthes_cornuta_ERR3712006 | 680795 | 126670 | 0.186 | 339 | 301 | 291 | 261 | 191 | 108 | 0 | 0 |
| Nepenthes_danseri_ERR3672061 | 230457 | 40921 | 0.178 | 343 | 270 | 260 | 218 | 133 | 73 | 0 | 0 |
| Nepenthes_deaniana_ERR3712177 | 1336806 | 254808 | 0.191 | 349 | 327 | 315 | 287 | 219 | 134 | 0 | 0 |
| Nepenthes_densiflora_ERR3662067 | 715746 | 148122 | 0.207 | 345 | 309 | 299 | 273 | 195 | 112 | 0 | 0 |
| Nepenthes_diatas_ERR3662280 | 1069545 | 195019 | 0.182 | 350 | 320 | 310 | 289 | 225 | 129 | 0 | 0 |
| Nepenthes_distillatoria_ERR3662281 | 1516024 | 208352 | 0.137 | 350 | 323 | 311 | 289 | 227 | 135 | 0 | 1 |
| Nepenthes_dubia_ERR3657203 | 365649 | 44028 | 0.12 | 341 | 264 | 259 | 226 | 145 | 76 | 0 | 0 |
| Nepenthes_edwardsiana_ERR3660991 | 509493 | 55550 | 0.109 | 345 | 282 | 270 | 236 | 160 | 86 | 0 | 0 |
| Nepenthes_ephippiata_ERR3660992 | 376506 | 53721 | 0.143 | 343 | 286 | 273 | 240 | 153 | 80 | 0 | 0 |
| Nepenthes_ephippiata_G07857 | 4250937 | 265169 | 0.062 | 349 | 320 | 308 | 276 | 206 | 111 | 0 | 0 |
| Nepenthes_ephippiata_G07900 | 2413742 | 147753 | 0.061 | 344 | 307 | 297 | 262 | 189 | 95 | 0 | 0 |
| Nepenthes_eustachya_ERR3662068 | 945043 | 128430 | 0.136 | 344 | 284 | 271 | 239 | 165 | 90 | 0 | 0 |
| Nepenthes_eymae_ERR3662871 | 1110453 | 114068 | 0.103 | 346 | 306 | 290 | 262 | 183 | 106 | 0 | 2 |
| Nepenthes_eymae_G07887 | 3057588 | 225016 | 0.074 | 341 | 321 | 301 | 255 | 174 | 92 | 0 | 0 |
| Nepenthes_eymae_G08121 | 1743728 | 131038 | 0.075 | 337 | 301 | 286 | 241 | 157 | 72 | 0 | 0 |
| Nepenthes_faizaliana_ERR3662069 | 840507 | 171409 | 0.204 | 344 | 312 | 301 | 272 | 203 | 115 | 0 | 0 |
| Nepenthes_faizaliana_G08090 | 2142254 | 221083 | 0.103 | 341 | 320 | 305 | 269 | 187 | 100 | 0 | 0 |
| Nepenthes_flava_ERR3712178 | 535728 | 84354 | 0.157 | 342 | 291 | 283 | 255 | 188 | 104 | 0 | 0 |
| Nepenthes_flava_G08119 | 3340025 | 212617 | 0.064 | 350 | 312 | 300 | 267 | 188 | 103 | 0 | 0 |
| Nepenthes_glabrata_ERR3662070 | 1052501 | 210121 | 0.2 | 345 | 317 | 303 | 277 | 211 | 121 | 0 | 0 |
| Nepenthes_glandulifera_ERR3662282 | 849220 | 137237 | 0.162 | 347 | 307 | 291 | 252 | 180 | 101 | 0 | 1 |
| Nepenthes_glandulifera_G07830 | 5411128 | 288658 | 0.053 | 350 | 331 | 315 | 296 | 226 | 119 | 0 | 0 |
| Nepenthes_graciliflora_ERR3661586 | 1281482 | 89483 | 0.07 | 351 | 281 | 264 | 228 | 157 | 88 | 0 | 0 |
| Nepenthes_graciliflora_ERR3662284 | 602117 | 101027 | 0.168 | 347 | 303 | 297 | 267 | 202 | 111 | 0 | 0 |
| Nepenthes_graciliflora_G08092 | 4122883 | 421073 | 0.102 | 346 | 326 | 312 | 279 | 205 | 115 | 0 | 0 |
| Nepenthes_gracilis_ERR3657205 | 373848 | 46916 | 0.125 | 345 | 274 | 264 | 225 | 144 | 76 | 0 | 0 |
| Nepenthes_gracilis_ERR3662283 | 1222957 | 229852 | 0.188 | 350 | 325 | 313 | 286 | 214 | 131 | 0 | 0 |
| Nepenthes_gracilis_G08091 | 5109324 | 316090 | 0.062 | 351 | 331 | 320 | 293 | 221 | 127 | 0 | 0 |
| Nepenthes_gracillima_ERR3662285 | 1184230 | 190372 | 0.161 | 349 | 317 | 309 | 274 | 214 | 126 | 0 | 2 |
| Nepenthes_gymnamphora_ERR3662071 | 707309 | 123839 | 0.175 | 343 | 293 | 283 | 257 | 187 | 114 | 0 | 0 |
| Nepenthes_gymnamphora_ERR3662923 | 6814916 | 236274 | 0.035 | 348 | 315 | 297 | 275 | 203 | 110 | 0 | 0 |
| Nepenthes_gymnamphora_G07893 | 1860840 | 99044 | 0.053 | 341 | 276 | 258 | 210 | 130 | 73 | 0 | 0 |
| Nepenthes_gymnamphora_G07901 | 3019763 | 292722 | 0.097 | 347 | 323 | 308 | 283 | 208 | 122 | 0 | 1 |
| Nepenthes_hamata_ERR3657206 | 579180 | 49797 | 0.086 | 344 | 276 | 265 | 230 | 159 | 75 | 0 | 0 |
| Nepenthes_hamata_ERR3657207 | 331827 | 38417 | 0.116 | 342 | 269 | 258 | 222 | 148 | 74 | 0 | 0 |
| Nepenthes_hamata_G07884 | 3729433 | 440478 | 0.118 | 348 | 333 | 314 | 288 | 217 | 121 | 0 | 0 |
| Nepenthes_hamiguitanensis_ERR3657208 | 608369 | 55521 | 0.091 | 348 | 280 | 265 | 234 | 161 | 83 | 0 | 0 |
| Nepenthes_hemsleyana_ERR3662286 | 714908 | 117221 | 0.164 | 346 | 316 | 304 | 276 | 202 | 96 | 0 | 0 |
| Nepenthes_hirsuta_ERR3662072 | 710123 | 130541 | 0.184 | 346 | 300 | 286 | 257 | 194 | 113 | 0 | 1 |
| Nepenthes_hispida_ERR3712007 | 903255 | 148515 | 0.164 | 340 | 297 | 291 | 267 | 196 | 109 | 0 | 0 |
| Nepenthes_hurrelliana_G08258 | 2260754 | 78057 | 0.035 | 348 | 258 | 232 | 158 | 59 | 15 | 0 | 0 |
| Nepenthes_inermis_ERR3662073 | 864963 | 153220 | 0.177 | 342 | 302 | 293 | 265 | 208 | 115 | 0 | 1 |
| Nepenthes_insignis_ERR3661587 | 678821 | 79205 | 0.117 | 350 | 305 | 292 | 258 | 191 | 102 | 0 | 0 |
| Nepenthes_izumiae_ERR3662287 | 768603 | 117771 | 0.153 | 349 | 302 | 290 | 255 | 179 | 95 | 0 | 1 |
| Nepenthes_izumiae_G07891 | 3127038 | 144177 | 0.046 | 348 | 317 | 301 | 265 | 183 | 91 | 0 | 0 |
| Nepenthes_izumiae_x_ventricosa_G07866 | 3026660 | 137582 | 0.045 | 349 | 311 | 295 | 255 | 169 | 89 | 0 | 0 |
| Nepenthes_jacquelineae_ERR3662074 | 984992 | 171708 | 0.174 | 340 | 306 | 296 | 266 | 199 | 116 | 0 | 0 |
| Nepenthes_jacquelineae_G07873 | 2258600 | 206031 | 0.091 | 344 | 312 | 299 | 267 | 188 | 105 | 0 | 0 |
| Nepenthes_jamban_ERR3662075 | 1268839 | 237587 | 0.187 | 347 | 310 | 297 | 277 | 211 | 122 | 0 | 2 |
| Nepenthes_justinae_ERR3712099 | 445529 | 50699 | 0.114 | 344 | 281 | 272 | 242 | 166 | 92 | 0 | 0 |
| Nepenthes_kampotiana_ERR3657987 | 585119 | 65013 | 0.111 | 346 | 277 | 265 | 237 | 152 | 91 | 0 | 1 |
| Nepenthes_kampotiana_G07895 | 4937630 | 325792 | 0.066 | 351 | 329 | 313 | 281 | 194 | 107 | 0 | 0 |
| Nepenthes_kerrii_ERR3712008 | 936796 | 127745 | 0.136 | 339 | 278 | 255 | 214 | 148 | 85 | 0 | 0 |
| Nepenthes_khasiana_ERR3662076 | 1261220 | 273721 | 0.217 | 349 | 327 | 313 | 282 | 215 | 131 | 0 | 1 |
| Nepenthes_khasiana_G07874 | 6099320 | 685051 | 0.112 | 349 | 334 | 321 | 297 | 234 | 141 | 1 | 0 |
| Nepenthes_klossii_ERR3661588 | 1354669 | 123498 | 0.091 | 351 | 299 | 279 | 251 | 183 | 103 | 0 | 0 |
| Nepenthes_kongkandana_ERR3662288 | 805452 | 153286 | 0.19 | 349 | 316 | 306 | 273 | 210 | 122 | 0 | 0 |
| Nepenthes_kongkandana_G07869 | 3869926 | 193920 | 0.05 | 349 | 318 | 299 | 257 | 167 | 84 | 0 | 0 |
| Nepenthes_lamii_ERR3661589 | 262695 | 11629 | 0.044 | 317 | 65 | 40 | 33 | 20 | 13 | 0 | 0 |
| Nepenthes_leonardoi_ERR3712100 | 482667 | 56074 | 0.116 | 347 | 285 | 273 | 240 | 159 | 89 | 0 | 0 |
| Nepenthes_lingulata_ERR3662078 | 815767 | 157026 | 0.192 | 345 | 304 | 292 | 259 | 191 | 111 | 0 | 0 |
| Nepenthes_longifolia_ERR3662289 | 872438 | 155697 | 0.178 | 350 | 320 | 302 | 274 | 202 | 117 | 0 | 0 |
| Nepenthes_longifolia_G07763 | 4117512 | 195554 | 0.047 | 352 | 309 | 292 | 259 | 175 | 96 | 0 | 0 |
| Nepenthes_lowii_ERR3662085 | 827675 | 155418 | 0.188 | 343 | 309 | 295 | 266 | 200 | 117 | 0 | 0 |
| Nepenthes_lowii_x_campanulata_G08109 | 4422100 | 351430 | 0.079 | 350 | 327 | 315 | 273 | 201 | 108 | 0 | 0 |
| Nepenthes_macfarlanei_ERR3662290 | 1638289 | 302496 | 0.185 | 352 | 327 | 312 | 288 | 227 | 143 | 0 | 0 |
| Nepenthes_macfarlanei_G08257 | 2757282 | 163103 | 0.059 | 344 | 228 | 155 | 95 | 30 | 7 | 0 | 0 |
| Nepenthes_macrophylla_ERR3662086 | 815735 | 150140 | 0.184 | 343 | 308 | 298 | 270 | 198 | 110 | 0 | 0 |
| Nepenthes_macrovulgaris_ERR3657988 | 688975 | 55960 | 0.081 | 348 | 274 | 264 | 227 | 159 | 83 | 0 | 0 |
| Nepenthes_madagascariensis_ERR3672062 | 189045 | 41291 | 0.218 | 339 | 280 | 266 | 227 | 141 | 71 | 0 | 0 |
| Nepenthes_mantalingajanensis_ERR3712101 | 754602 | 60053 | 0.08 | 347 | 276 | 260 | 234 | 150 | 85 | 0 | 0 |
| Nepenthes_mapuluensis_ERR3662087 | 1198042 | 242658 | 0.203 | 346 | 321 | 303 | 276 | 220 | 129 | 0 | 0 |
| Nepenthes_masoalensis_ERR3662088 | 2439341 | 482518 | 0.198 | 348 | 331 | 319 | 300 | 227 | 136 | 0 | 1 |
| Nepenthes_maxima_ERR3661602 | 1108241 | 127660 | 0.115 | 350 | 308 | 289 | 260 | 185 | 99 | 0 | 1 |
| Nepenthes_maxima_ERR3661604 | 804533 | 66263 | 0.082 | 343 | 250 | 223 | 189 | 125 | 66 | 0 | 0 |
| Nepenthes_maxima_ERR3661605 | 676305 | 93406 | 0.138 | 351 | 299 | 287 | 257 | 189 | 100 | 0 | 0 |
| Nepenthes_maxima_ERR3661606 | 707721 | 36751 | 0.052 | 335 | 146 | 105 | 79 | 44 | 25 | 0 | 0 |
| Nepenthes_maxima_ERR3661607 | 822352 | 104170 | 0.127 | 349 | 307 | 297 | 272 | 205 | 112 | 0 | 0 |
| Nepenthes_maxima_ERR3661608 | 781241 | 86238 | 0.11 | 348 | 293 | 281 | 245 | 179 | 101 | 0 | 2 |
| Nepenthes_maxima_ERR3662875 | 762686 | 79606 | 0.104 | 346 | 305 | 294 | 263 | 191 | 99 | 0 | 0 |
| Nepenthes_maxima_G07859 | 3914929 | 228093 | 0.058 | 349 | 320 | 309 | 278 | 202 | 111 | 0 | 0 |
| Nepenthes_maxima_G07861 | 4261653 | 267946 | 0.063 | 348 | 330 | 319 | 290 | 214 | 118 | 0 | 0 |
| Nepenthes_maxima_G07864 | 4045755 | 394641 | 0.098 | 342 | 333 | 312 | 282 | 203 | 108 | 0 | 1 |
| Nepenthes_maxima_G07894 | 3204065 | 208917 | 0.065 | 346 | 318 | 307 | 280 | 197 | 106 | 0 | 0 |
| Nepenthes_maxima_G08103 | 4241673 | 251572 | 0.059 | 347 | 315 | 302 | 265 | 187 | 99 | 0 | 0 |
| Nepenthes_maxima_G08107 | 3661653 | 287166 | 0.078 | 345 | 325 | 312 | 283 | 210 | 125 | 0 | 1 |
| Nepenthes_merrilliana_ERR3662291 | 534384 | 85520 | 0.16 | 348 | 295 | 289 | 260 | 191 | 104 | 0 | 0 |
| Nepenthes_merrilliana_G08089 | 2881445 | 246658 | 0.086 | 344 | 319 | 303 | 272 | 187 | 95 | 0 | 0 |
| Nepenthes_micramphora_ERR3657989 | 815229 | 70906 | 0.087 | 348 | 284 | 276 | 248 | 170 | 94 | 0 | 0 |
| Nepenthes_mikei_ERR3662089 | 845761 | 154752 | 0.183 | 342 | 302 | 295 | 268 | 202 | 109 | 0 | 1 |
| Nepenthes_mindanaoensis_ERR3661609 | 1046043 | 138062 | 0.132 | 349 | 310 | 300 | 277 | 205 | 119 | 0 | 0 |
| Nepenthes_minima_ERR3662292 | 1169861 | 223946 | 0.191 | 350 | 327 | 317 | 292 | 221 | 129 | 0 | 0 |
| Nepenthes_minima_G07872 | 4674596 | 275507 | 0.059 | 348 | 323 | 311 | 277 | 202 | 113 | 0 | 0 |
| Nepenthes_mira_ERR3662090 | 1309453 | 231454 | 0.177 | 346 | 312 | 301 | 279 | 212 | 129 | 0 | 0 |
| Nepenthes_mira_G07902 | 4501431 | 454250 | 0.101 | 350 | 328 | 312 | 291 | 219 | 132 | 0 | 0 |
| Nepenthes_mirabilis_ERR3657198 | 291340 | 37693 | 0.129 | 347 | 250 | 242 | 209 | 131 | 66 | 0 | 0 |
| Nepenthes_mirabilis_ERR3657199 | 437207 | 50881 | 0.116 | 344 | 284 | 273 | 234 | 157 | 81 | 0 | 0 |
| Nepenthes_mirabilis_ERR3657990 | 52173 | 5722 | 0.11 | 309 | 33 | 21 | 10 | 5 | 2 | 0 | 0 |
| Nepenthes_mirabilis_ERR3657991 | 534171 | 22587 | 0.042 | 327 | 110 | 80 | 65 | 39 | 26 | 0 | 0 |
| Nepenthes_mirabilis_ERR3661610 | 1027921 | 82918 | 0.081 | 349 | 303 | 288 | 252 | 182 | 104 | 0 | 0 |
| Nepenthes_mirabilis_ERR3661611 | 895165 | 101167 | 0.113 | 350 | 306 | 298 | 260 | 183 | 103 | 0 | 0 |
| Nepenthes_mirabilis_ERR3661612 | 234851 | 14770 | 0.063 | 320 | 58 | 44 | 35 | 20 | 11 | 0 | 0 |
| Nepenthes_mirabilis_ERR3661613 | 570779 | 60549 | 0.106 | 346 | 258 | 244 | 196 | 140 | 78 | 0 | 0 |
| Nepenthes_mirabilis_ERR3661614 | 614990 | 38108 | 0.062 | 339 | 192 | 161 | 132 | 84 | 47 | 0 | 0 |
| Nepenthes_mirabilis_ERR3661615 | 477119 | 32787 | 0.069 | 342 | 193 | 171 | 142 | 88 | 45 | 0 | 0 |
| Nepenthes_mirabilis_ERR3661616 | 654433 | 61799 | 0.094 | 349 | 288 | 277 | 245 | 171 | 96 | 0 | 0 |
| Nepenthes_mirabilis_ERR3661617 | 456725 | 30118 | 0.066 | 342 | 186 | 167 | 136 | 92 | 46 | 0 | 0 |
| Nepenthes_mirabilis_ERR3661618 | 532065 | 37929 | 0.071 | 338 | 209 | 193 | 160 | 103 | 61 | 0 | 0 |
| Nepenthes_mirabilis_ERR3661619 | 568122 | 45323 | 0.08 | 345 | 240 | 219 | 179 | 119 | 66 | 0 | 0 |
| Nepenthes_mirabilis_ERR3661620 | 865086 | 85961 | 0.099 | 350 | 304 | 296 | 264 | 179 | 103 | 0 | 0 |
| Nepenthes_mirabilis_ERR3661621 | 692408 | 65024 | 0.094 | 348 | 304 | 285 | 244 | 165 | 100 | 0 | 0 |
| Nepenthes_mirabilis_ERR3661622 | 690228 | 69568 | 0.101 | 348 | 295 | 283 | 251 | 168 | 90 | 0 | 0 |
| Nepenthes_mirabilis_ERR3662877 | 1012921 | 88368 | 0.087 | 346 | 275 | 243 | 201 | 131 | 73 | 0 | 0 |
| Nepenthes_mirabilis_ERR3662928 | 675149 | 65104 | 0.096 | 346 | 299 | 288 | 252 | 171 | 96 | 0 | 0 |
| Nepenthes_mirabilis_ERR3662929 | 808297 | 87594 | 0.108 | 351 | 302 | 290 | 262 | 188 | 105 | 0 | 0 |
| Nepenthes_mirabilis_ERR3672063 | 137976 | 26678 | 0.193 | 343 | 237 | 230 | 191 | 96 | 44 | 0 | 0 |
| Nepenthes_mirabilis_G08086 | 3489362 | 334876 | 0.096 | 349 | 329 | 310 | 279 | 195 | 101 | 0 | 0 |
| Nepenthes_mirabilis_G08094 | 3142346 | 181576 | 0.058 | 349 | 307 | 292 | 259 | 173 | 94 | 0 | 0 |
| Nepenthes_mirabilis_G08099 | 3423605 | 209209 | 0.061 | 349 | 310 | 298 | 262 | 180 | 95 | 0 | 0 |
| Nepenthes_mirabilis_G08236 | 2927267 | 168037 | 0.057 | 348 | 315 | 300 | 254 | 157 | 77 | 0 | 0 |
| Nepenthes_mirabilis_G08237 | 2476255 | 143371 | 0.058 | 350 | 315 | 299 | 245 | 154 | 70 | 0 | 0 |
| Nepenthes_mirabilis_G08244 | 2536953 | 174726 | 0.069 | 349 | 327 | 310 | 279 | 199 | 112 | 0 | 0 |
| Nepenthes_mirabilis_G08245 | 3891446 | 207544 | 0.053 | 350 | 325 | 310 | 284 | 213 | 121 | 0 | 0 |
| Nepenthes_mirabilis_G08246 | 2211549 | 154299 | 0.07 | 349 | 324 | 305 | 276 | 193 | 100 | 0 | 0 |
| Nepenthes_mirabilis_G08247 | 1948539 | 115349 | 0.059 | 348 | 318 | 299 | 270 | 187 | 104 | 0 | 0 |
| Nepenthes_mirabilis_var_echinostoma_ERR3657204 | 1774998 | 202447 | 0.114 | 352 | 326 | 308 | 284 | 209 | 127 | 0 | 0 |
| Nepenthes_mirabilis_var_echinostoma_G08131 | 2788251 | 255280 | 0.092 | 340 | 323 | 303 | 270 | 171 | 85 | 0 | 0 |
| Nepenthes_mirabilis_var_globosa_G08128 | 2275620 | 222723 | 0.098 | 342 | 322 | 300 | 258 | 168 | 84 | 0 | 0 |
| Nepenthes_mollis_ERR3712179 | 570413 | 81441 | 0.143 | 346 | 298 | 290 | 255 | 176 | 90 | 0 | 0 |
| Nepenthes_monticola_ERR3712102 | 152122 | 21265 | 0.14 | 336 | 207 | 196 | 152 | 77 | 36 | 0 | 0 |
| Nepenthes_muluensis_ERR3657371 | 1206595 | 133553 | 0.111 | 349 | 313 | 301 | 272 | 205 | 113 | 0 | 2 |
| Nepenthes_murudensis_ERR3662293 | 680292 | 103927 | 0.153 | 349 | 304 | 299 | 265 | 200 | 105 | 0 | 1 |
| Nepenthes_naga_ERR3662294 | 517758 | 85044 | 0.164 | 346 | 300 | 294 | 258 | 193 | 102 | 0 | 0 |
| Nepenthes_neoguineensis_ERR3657378 | 210997 | 19475 | 0.092 | 335 | 176 | 164 | 137 | 67 | 34 | 0 | 0 |
| Nepenthes_neoguineensis_ERR3662053 | 610629 | 36136 | 0.059 | 340 | 193 | 169 | 148 | 94 | 55 | 0 | 0 |
| Nepenthes_neoguineensis_G08125 | 2692447 | 217001 | 0.081 | 338 | 316 | 296 | 248 | 167 | 95 | 0 | 0 |
| Nepenthes_northiana_ERR3662295 | 779409 | 131546 | 0.169 | 347 | 313 | 305 | 270 | 197 | 113 | 0 | 0 |
| Nepenthes_northiana_G08135 | 2739114 | 224907 | 0.082 | 347 | 323 | 304 | 272 | 186 | 103 | 0 | 0 |
| Nepenthes_oblanceolata_ERR3712103 | 257440 | 26413 | 0.103 | 335 | 213 | 199 | 172 | 110 | 57 | 0 | 0 |
| Nepenthes_ovata_ERR3662296 | 677568 | 108488 | 0.16 | 344 | 306 | 294 | 266 | 199 | 113 | 0 | 0 |
| Nepenthes_ovata_G08118 | 3720387 | 242607 | 0.065 | 348 | 318 | 301 | 272 | 206 | 113 | 0 | 0 |
| Nepenthes_palawanensis_ERR3712010 | 1132668 | 166729 | 0.147 | 346 | 296 | 277 | 245 | 172 | 94 | 0 | 0 |
| Nepenthes_papuana_ERR3657379 | 319436 | 33359 | 0.104 | 344 | 249 | 237 | 202 | 128 | 69 | 0 | 0 |
| Nepenthes_parvula_G08240 | 3097062 | 171843 | 0.055 | 349 | 319 | 301 | 249 | 156 | 74 | 0 | 0 |
| Nepenthes_parvula_G08248 | 2236301 | 128599 | 0.058 | 345 | 324 | 307 | 270 | 179 | 94 | 0 | 0 |
| Nepenthes_peltata_ERR3662297 | 640644 | 123511 | 0.193 | 348 | 311 | 300 | 276 | 209 | 120 | 0 | 0 |
| Nepenthes_pervillei_ERR3657380 | 730331 | 80629 | 0.11 | 349 | 300 | 292 | 261 | 181 | 97 | 0 | 1 |
| Nepenthes_pervillei_G08123 | 2906040 | 409683 | 0.141 | 344 | 329 | 308 | 275 | 193 | 105 | 0 | 1 |
| Nepenthes_petiolata_ERR3662091 | 620204 | 109039 | 0.176 | 342 | 302 | 292 | 261 | 194 | 106 | 0 | 1 |
| Nepenthes_petiolata_G07881 | 1869390 | 131376 | 0.07 | 343 | 307 | 289 | 255 | 163 | 78 | 0 | 0 |
| Nepenthes_philippinensis_ERR3662299 | 926433 | 173015 | 0.187 | 351 | 319 | 308 | 281 | 217 | 125 | 0 | 1 |
| Nepenthes_philippinensis_G07878 | 4812380 | 447914 | 0.093 | 350 | 329 | 309 | 284 | 216 | 128 | 0 | 0 |
| Nepenthes_pitopangii_ERR3712011 | 1405236 | 263110 | 0.187 | 345 | 309 | 294 | 267 | 198 | 124 | 0 | 1 |
| Nepenthes_platychila_ERR3662300 | 660206 | 122190 | 0.185 | 348 | 317 | 305 | 272 | 203 | 108 | 0 | 0 |
| Nepenthes_pulchra_ERR3657381 | 680733 | 56529 | 0.083 | 346 | 279 | 271 | 235 | 160 | 86 | 0 | 0 |
| Nepenthes_rafflesiana_ERR3657382 | 292457 | 32913 | 0.113 | 344 | 244 | 232 | 201 | 126 | 64 | 0 | 0 |
| Nepenthes_rafflesiana_ERR3657383 | 390207 | 44747 | 0.115 | 344 | 270 | 259 | 224 | 144 | 79 | 0 | 0 |
| Nepenthes_rafflesiana_ERR3662092 | 729538 | 106221 | 0.146 | 340 | 297 | 290 | 256 | 185 | 97 | 0 | 1 |
| Nepenthes_rafflesiana_G08097 | 3392543 | 183487 | 0.054 | 344 | 309 | 296 | 258 | 178 | 96 | 0 | 0 |
| Nepenthes_rafflesiana_G08127 | 2054762 | 173990 | 0.085 | 338 | 318 | 297 | 249 | 153 | 74 | 0 | 0 |
| Nepenthes_rafflesiana_x_ampullaria_ERR3662102 | 1331983 | 235679 | 0.177 | 349 | 320 | 306 | 269 | 197 | 108 | 0 | 5 |
| Nepenthes_rafflesiana_x_ampullaria_G08126 | 3903412 | 199255 | 0.051 | 350 | 326 | 310 | 277 | 198 | 108 | 0 | 0 |
| Nepenthes_rajah_ERR3657984 | 1575112 | 147895 | 0.094 | 350 | 303 | 277 | 239 | 165 | 101 | 0 | 0 |
| Nepenthes_rajah_G08114 | 5538480 | 473827 | 0.086 | 351 | 325 | 315 | 285 | 217 | 125 | 0 | 0 |
| Nepenthes_ramispina_ERR3662301 | 902229 | 148564 | 0.165 | 347 | 306 | 296 | 273 | 208 | 122 | 0 | 3 |
| Nepenthes_reinwardtiana_ERR3662093 | 924752 | 189406 | 0.205 | 348 | 313 | 302 | 272 | 217 | 121 | 0 | 2 |
| Nepenthes_reinwardtiana_G08100 | 3937174 | 257170 | 0.065 | 348 | 313 | 298 | 266 | 193 | 101 | 0 | 0 |
| Nepenthes_reinwardtiana_G08133 | 1366297 | 134392 | 0.098 | 335 | 307 | 283 | 236 | 139 | 71 | 0 | 0 |
| Nepenthes_rhombicaulis_ERR3662302 | 524999 | 80742 | 0.154 | 348 | 292 | 286 | 254 | 186 | 96 | 0 | 0 |
| Nepenthes_rigidifolia_ERR3712180 | 571497 | 94054 | 0.165 | 347 | 302 | 293 | 263 | 183 | 103 | 0 | 2 |
| Nepenthes_robcantleyi_ERR3712181 | 635443 | 126903 | 0.2 | 345 | 310 | 300 | 273 | 204 | 112 | 0 | 0 |
| Nepenthes_robcantleyi_G08111 | 2953211 | 266242 | 0.09 | 348 | 317 | 301 | 273 | 199 | 101 | 0 | 0 |
| Nepenthes_rowaniae_ERR3662094 | 812199 | 108667 | 0.134 | 342 | 286 | 271 | 225 | 149 | 85 | 0 | 1 |
| Nepenthes_rowaniae_G05999 | 2940278 | 273422 | 0.093 | 345 | 325 | 310 | 269 | 192 | 106 | 0 | 0 |
| Nepenthes_rowaniae_G08233 | 1924143 | 95779 | 0.05 | 352 | 294 | 274 | 208 | 118 | 48 | 0 | 0 |
| Nepenthes_rowaniae_G08234 | 1882869 | 132953 | 0.071 | 349 | 315 | 302 | 266 | 163 | 85 | 0 | 0 |
| Nepenthes_rowaniae_G08239 | 3040778 | 223519 | 0.074 | 349 | 324 | 306 | 260 | 164 | 82 | 0 | 0 |
| Nepenthes_rowaniae_G08249 | 2506171 | 176788 | 0.071 | 348 | 331 | 313 | 277 | 199 | 109 | 0 | 0 |
| Nepenthes_sanguinea_ERR3662095 | 807648 | 164480 | 0.204 | 342 | 307 | 294 | 263 | 198 | 112 | 0 | 0 |
| Nepenthes_sanguinea_G07858 | 3447958 | 332791 | 0.097 | 343 | 321 | 301 | 267 | 194 | 101 | 0 | 0 |
| Nepenthes_sibuyanensis_ERR3662096 | 462562 | 69150 | 0.149 | 337 | 277 | 269 | 239 | 166 | 86 | 0 | 0 |
| Nepenthes_sibuyanensis_G08113 | 3164475 | 199412 | 0.063 | 352 | 319 | 302 | 262 | 187 | 95 | 0 | 0 |
| Nepenthes_singalana_ERR3662303 | 431954 | 75366 | 0.174 | 344 | 293 | 284 | 243 | 166 | 97 | 0 | 0 |
| Nepenthes_singalana_G07880 | 1785699 | 192757 | 0.108 | 346 | 312 | 295 | 266 | 182 | 104 | 0 | 0 |
| Nepenthes_smilesii_ERR3662924 | 1.20E+07 | 512521 | 0.041 | 350 | 336 | 313 | 284 | 229 | 133 | 0 | 0 |
| Nepenthes_smilesii_ERR3662925 | 1.40E+07 | 438414 | 0.032 | 352 | 330 | 304 | 281 | 216 | 115 | 0 | 0 |
| Nepenthes_sp_adrianii_ERR3712004 | 243711 | 44482 | 0.183 | 326 | 256 | 251 | 218 | 142 | 69 | 0 | 0 |
| Nepenthes_sp_Anipahan_ERR3712104 | 1077945 | 120395 | 0.112 | 349 | 316 | 302 | 275 | 202 | 119 | 0 | 1 |
| Nepenthes_sp_BM-2019_ERR3712119 | 367328 | 39807 | 0.108 | 343 | 254 | 242 | 214 | 130 | 74 | 0 | 1 |
| Nepenthes_sp_calcicola_G08235 | 4718340 | 327607 | 0.069 | 349 | 336 | 306 | 264 | 174 | 80 | 0 | 0 |
| Nepenthes_spathulata_ERR3662304 | 299786 | 49709 | 0.166 | 337 | 267 | 261 | 227 | 156 | 76 | 0 | 1 |
| Nepenthes_spathulata_G07896 | 2877227 | 244525 | 0.085 | 343 | 316 | 296 | 253 | 169 | 88 | 0 | 0 |
| Nepenthes_spathulata_G08116 | 2027411 | 145835 | 0.072 | 341 | 303 | 291 | 252 | 172 | 83 | 0 | 0 |
| Nepenthes_spectabilis_ERR3657985 | 487816 | 38607 | 0.079 | 348 | 254 | 243 | 203 | 123 | 67 | 0 | 0 |
| Nepenthes_spectabilis_ERR3662055 | 520635 | 26777 | 0.051 | 339 | 132 | 104 | 81 | 40 | 24 | 0 | 0 |
| Nepenthes_spectabilis_G07890 | 3086893 | 276823 | 0.09 | 345 | 318 | 304 | 275 | 200 | 115 | 0 | 0 |
| Nepenthes_stenophylla_ERR3662305 | 713657 | 125358 | 0.176 | 347 | 315 | 307 | 276 | 203 | 114 | 0 | 1 |
| Nepenthes_stenophylla_G08115 | 4407154 | 140223 | 0.032 | 348 | 317 | 304 | 265 | 192 | 98 | 0 | 0 |
| Nepenthes_sumagaya_ERR3712012 | 804764 | 146373 | 0.182 | 340 | 304 | 292 | 259 | 188 | 101 | 0 | 0 |
| Nepenthes_suratensis_ERR3712013 | 899615 | 160008 | 0.178 | 344 | 308 | 298 | 265 | 208 | 116 | 0 | 0 |
| Nepenthes_surigaoensis_ERR3662315 | 571253 | 97118 | 0.17 | 348 | 305 | 296 | 262 | 198 | 109 | 0 | 0 |
| Nepenthes_talangensis_ERR3662326 | 388573 | 67007 | 0.172 | 341 | 290 | 281 | 243 | 167 | 86 | 0 | 0 |
| Nepenthes_talangensis_G07888 | 1854001 | 186836 | 0.101 | 343 | 314 | 297 | 263 | 180 | 95 | 0 | 0 |
| Nepenthes_tenax_ERR3662327 | 635800 | 93159 | 0.147 | 345 | 298 | 287 | 252 | 171 | 100 | 0 | 0 |
| Nepenthes_tenax_G06000 | 3511078 | 329806 | 0.094 | 347 | 331 | 316 | 279 | 192 | 109 | 0 | 0 |
| Nepenthes_tenax_G08087 | 3125698 | 289428 | 0.093 | 345 | 326 | 311 | 276 | 198 | 110 | 0 | 0 |
| Nepenthes_tenax_G08238 | 3291689 | 207012 | 0.063 | 349 | 325 | 309 | 268 | 167 | 78 | 0 | 0 |
| Nepenthes_tentaculata_ERR3662056 | 816851 | 104594 | 0.128 | 350 | 304 | 294 | 266 | 193 | 105 | 0 | 0 |
| Nepenthes_tentaculata_ERR3662328 | 1008980 | 192335 | 0.191 | 349 | 324 | 312 | 282 | 219 | 128 | 0 | 1 |
| Nepenthes_tentaculata_G07885 | 3142858 | 310011 | 0.099 | 345 | 331 | 312 | 280 | 194 | 98 | 0 | 0 |
| Nepenthes_tenuis_ERR3662329 | 565273 | 102494 | 0.181 | 343 | 304 | 297 | 268 | 202 | 108 | 0 | 0 |
| Nepenthes_thai_ERR3657986 | 318443 | 32183 | 0.101 | 337 | 237 | 225 | 192 | 124 | 52 | 0 | 0 |
| Nepenthes_thai_G07879 | 4613549 | 185267 | 0.04 | 350 | 319 | 299 | 269 | 195 | 113 | 0 | 0 |
| Nepenthes_thorelii_ERR3662097 | 829019 | 146772 | 0.177 | 343 | 304 | 294 | 265 | 202 | 107 | 0 | 0 |
| Nepenthes_tobaica_ERR3662098 | 497336 | 89910 | 0.181 | 340 | 295 | 285 | 250 | 174 | 87 | 0 | 0 |
| Nepenthes_tobaica_ERR3662099 | 547126 | 103006 | 0.188 | 339 | 298 | 288 | 249 | 171 | 89 | 0 | 0 |
| Nepenthes_tobaica_G07871 | 3472469 | 153544 | 0.044 | 345 | 306 | 289 | 258 | 173 | 90 | 0 | 0 |
| Nepenthes_tobaica_G08122 | 2608163 | 208501 | 0.08 | 339 | 311 | 288 | 247 | 154 | 73 | 0 | 0 |
| Nepenthes_tomoriana_ERR3662330 | 625140 | 77699 | 0.124 | 349 | 280 | 263 | 223 | 145 | 81 | 0 | 0 |
| Nepenthes_treubiana_ERR3657197 | 1033955 | 68755 | 0.066 | 350 | 291 | 273 | 247 | 162 | 90 | 0 | 0 |
| Nepenthes_treubiana_ERR3662100 | 1041811 | 185528 | 0.178 | 347 | 314 | 306 | 275 | 211 | 122 | 0 | 0 |
| Nepenthes_truncata_ERR3662057 | 373123 | 46222 | 0.124 | 346 | 279 | 263 | 226 | 147 | 78 | 0 | 0 |
| Nepenthes_truncata_G07889 | 3686589 | 197968 | 0.054 | 351 | 325 | 309 | 273 | 197 | 95 | 0 | 0 |
| Nepenthes_truncata_G08124 | 2842204 | 248136 | 0.087 | 341 | 321 | 298 | 254 | 166 | 88 | 0 | 0 |
| Nepenthes_truncata_x_ventricosa_G08105 | 2325690 | 131522 | 0.057 | 348 | 300 | 283 | 239 | 151 | 76 | 0 | 0 |
| Nepenthes_veitchii_ERR3662101 | 628861 | 122551 | 0.195 | 343 | 304 | 291 | 258 | 195 | 99 | 0 | 0 |
| Nepenthes_veitchii_G08088 | 3350816 | 304879 | 0.091 | 345 | 319 | 305 | 273 | 195 | 104 | 0 | 0 |
| Nepenthes_veitchii_G08096 | 4857314 | 228284 | 0.047 | 345 | 310 | 294 | 261 | 180 | 96 | 0 | 0 |
| Nepenthes_veitchii_x_eymae_G08108 | 4429547 | 316064 | 0.071 | 350 | 320 | 308 | 284 | 208 | 113 | 0 | 0 |
| Nepenthes_ventricosa_ERR3672064 | 150864 | 25693 | 0.17 | 338 | 232 | 226 | 193 | 118 | 57 | 0 | 0 |
| Nepenthes_ventricosa_G07860 | 4119380 | 165408 | 0.04 | 350 | 322 | 293 | 257 | 173 | 83 | 0 | 0 |
| Nepenthes_ventricosa_G07865 | 3342732 | 189463 | 0.057 | 351 | 316 | 300 | 268 | 188 | 102 | 0 | 0 |
| Nepenthes_vieillardii_ERR3662926 | 1014659 | 72568 | 0.072 | 330 | 122 | 59 | 15 | 4 | 2 | 0 | 0 |
| Nepenthes_vieillardii_ERR3662927 | 1331318 | 111204 | 0.084 | 320 | 125 | 68 | 24 | 1 | 1 | 0 | 0 |
| Nepenthes_villosa_ERR3662058 | 964033 | 109830 | 0.114 | 349 | 310 | 297 | 267 | 197 | 114 | 0 | 0 |
| Nepenthes_vogelii_ERR3672065 | 167305 | 32607 | 0.195 | 338 | 253 | 243 | 207 | 123 | 65 | 0 | 0 |
| Nepenthes_vogelii_G08110 | 4192910 | 303496 | 0.072 | 348 | 325 | 307 | 282 | 205 | 113 | 0 | 0 |
| Nepenthes_xiphioides_ERR3662331 | 702110 | 107249 | 0.153 | 347 | 307 | 294 | 267 | 195 | 110 | 0 | 0 |
| Nepenthes_xiphioides_G07876 | 2943160 | 296285 | 0.101 | 345 | 325 | 304 | 277 | 209 | 111 | 0 | 0 |
| Nepenthes_zakriana_G08120 | 2789919 | 242538 | 0.087 | 343 | 327 | 302 | 259 | 169 | 80 | 0 | 0 |
