## Appendix 5 - Dataset optimization for "HybPhaser: a workflow for the detection and phasing of hybrids in target capture datasets"

**Appendix 5: Dataset Optimization – Missing Data**

Results from the script ‘Rscript2__dataset_optimization.R’.

**1) Dataset optimisation: Samples and loci removed to reduce missing data**

Graphical overview:

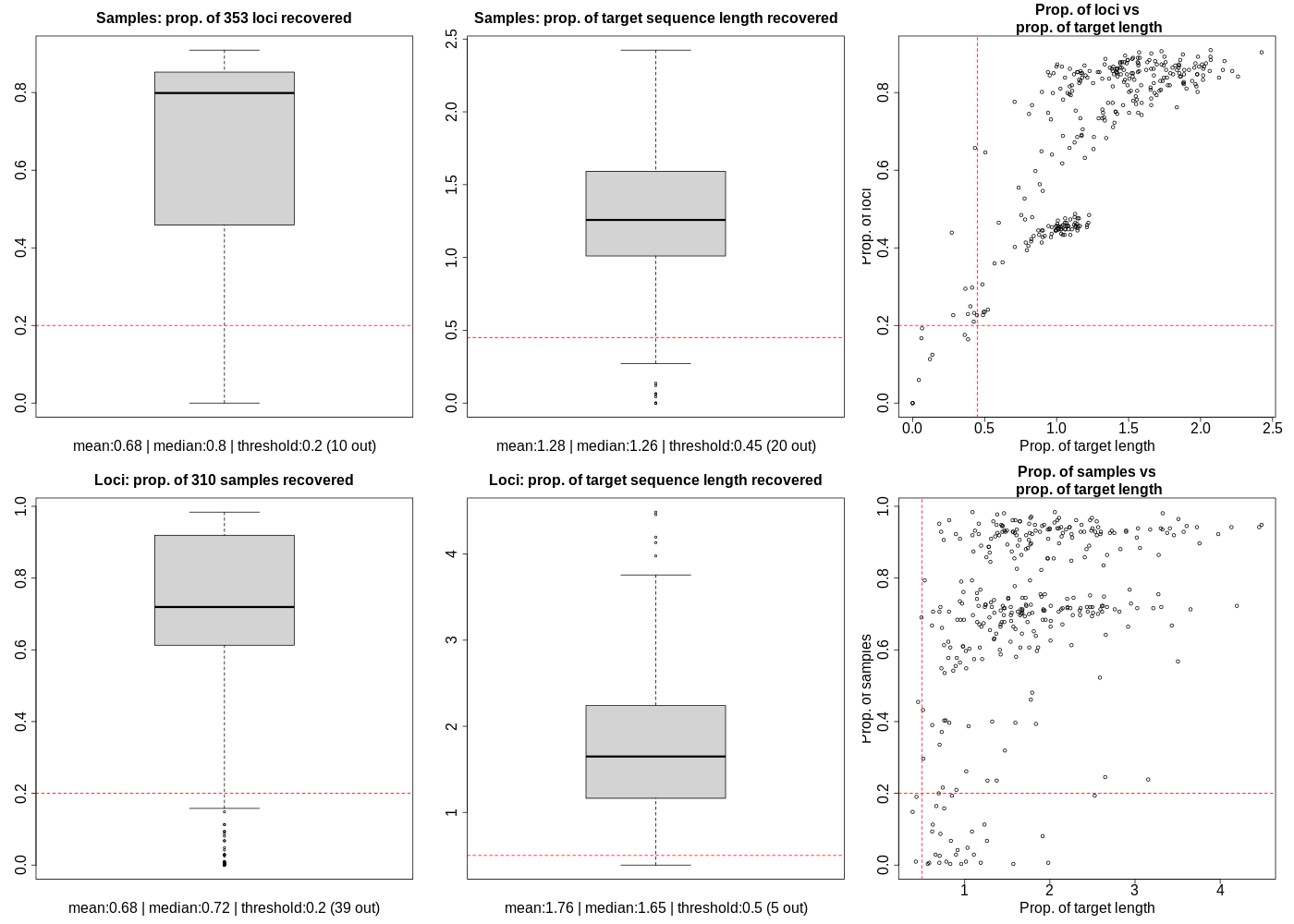

**Samples**

10 samples are below the threshold (0.2) for proportion of recovered loci:

| sample | proportion of loci recovered |
| --- | --- |
| Nepenthes_adnata_ERR3662059 | 0 |
| Nepenthes_lamii_ERR3661589 | 0.113 |
| Nepenthes_lavicola_ERR3662077 | 0 |
| Nepenthes_mirabilis_ERR3657990 | 0.059 |
| Nepenthes_mirabilis_ERR3661612 | 0.125 |
| Nepenthes_vieillardii_ERR3662926 | 0.167 |
| Nepenthes_vieillardii_ERR3662927 | 0.193 |
| Nepenthes_hamata_G07884 | 0.164 |
| Nepenthes_maxima_G07894 | 0.176 |
| Nepenthes_rowaniae_G08259 | 0 |

20 samples are below the threshold (0.45) for recovered target sequence length

| sample | recovered length as proportion of target sequence length |
| --- | --- |
| Nepenthes_adnata_ERR3662059 | 0 |
| Nepenthes_lamii_ERR3661589 | 0.12 |
| Nepenthes_lavicola_ERR3662077 | 0 |
| Nepenthes_maxima_ERR3661606 | 0.413 |
| Nepenthes_mirabilis_ERR3657990 | 0.043 |
| Nepenthes_mirabilis_ERR3657991 | 0.282 |
| Nepenthes_mirabilis_ERR3661612 | 0.137 |
| Nepenthes_mirabilis_ERR3661614 | 0.401 |
| Nepenthes_spectabilis_ERR3662055 | 0.367 |
| Nepenthes_vieillardii_ERR3662926 | 0.061 |
| Nepenthes_vieillardii_ERR3662927 | 0.065 |
| Nepenthes_graciliflora_G08092 | 0.447 |
| Nepenthes_hamata_G07884 | 0.385 |
| Nepenthes_hurrelliana_G08258 | 0.433 |
| Nepenthes_longifolia_G07763 | 0.425 |
| Nepenthes_macfarlanei_G08257 | 0.272 |
| Nepenthes_maxima_G07894 | 0.362 |
| Nepenthes_rowaniae_G08259 | 0 |
| Nepenthes_tenax_G08087 | 0.427 |
| Nepenthes_truncata_G07889 | 0.384 |

In total 20 samples were removed:

Nepenthes_adnata_ERR3662059, Nepenthes_lamii_ERR3661589, Nepenthes_lavicola_ERR3662077, Nepenthes_mirabilis_ERR3657990, Nepenthes_mirabilis_ERR3661612, Nepenthes_vieillardii_ERR3662926, Nepenthes_vieillardii_ERR3662927, Nepenthes_hamata_G07884, Nepenthes_maxima_G07894, Nepenthes_rowaniae_G08259, Nepenthes_maxima_ERR3661606, Nepenthes_mirabilis_ERR3657991, Nepenthes_mirabilis_ERR3661614, Nepenthes_spectabilis_ERR3662055, Nepenthes_graciliflora_G08092, Nepenthes_hurrelliana_G08258, Nepenthes_longifolia_G07763, Nepenthes_macfarlanei_G08257, Nepenthes_tenax_G08087, Nepenthes_truncata_G07889

**Loci**

39 loci are below the threshold (0.2) for proportion of recovered samples:

| Locus | proportion of samples recovered for |
| --- | --- |
| 5034 | 0 |
| 5260 | 0.006 |
| 5271 | 0.165 |
| 5328 | 0.081 |
| 5422 | 0.029 |
| 5660 | 0.026 |
| 5802 | 0.003 |
| 5941 | 0.158 |
| 5944 | 0.113 |
| 5990 | 0.094 |
| 6148 | 0.068 |
| 6150 | 0 |
| 6164 | 0.042 |
| 6270 | 0.006 |
| 6398 | 0.048 |
| 6401 | 0 |
| 6406 | 0.19 |
| 6430 | 0 |
| 6448 | 0.148 |
| 6457 | 0.01 |
| 6506 | 0.087 |
| 6514 | 0 |
| 6557 | 0.113 |
| 6565 | 0.029 |
| 6705 | 0.003 |
| 6732 | 0.003 |
| 6780 | 0.068 |
| 6785 | 0.01 |
| 6791 | 0.094 |
| 6864 | 0.003 |
| 6886 | 0.194 |
| 6893 | 0.006 |
| 6969 | 0 |
| 6977 | 0.029 |
| 7024 | 0.006 |
| 7194 | 0.194 |
| 7296 | 0 |
| 7325 | 0.01 |
| 7361 | 0 |

5 loci are below the threshold (0.5) for proportion of recovered target sequence length:

| Locus | proportion of target sequence length |
| --- | --- |
| 5339 | 0.453 |
| 5670 | 0.492 |
| 6406 | 0.433 |
| 6448 | 0.388 |
| 6457 | 0.426 |

In total 41 loci were removed:

5034, 5260, 5271, 5328, 5422, 5660, 5802, 5941, 5944, 5990, 6148, 6150, 6164, 6270, 6398, 6401, 6406, 6430, 6448, 6457, 6506, 6514, 6557, 6565, 6705, 6732, 6780, 6785, 6791, 6864, 6886, 6893, 6969, 6977, 7024, 7194, 7296, 7325, 7361, 5339, 5670
