## Appendix 6 - Summary table for "HybPhaser: a workflow for the detection and phasing of hybrids in target capture datasets"

**Appendix 5: Dataset Optimization – Putative Paralogs**

**Paralogous genes**

Information gathered from R script “2_optimize_dataset.R”

**Putative paralogs removed for all samples**

Figure 1: Bar graph and boxplot of mean proportion of SNPs across all samples for each locus.

Red line shows the threshold. Red bars show outlier loci that are removed from the dataset.

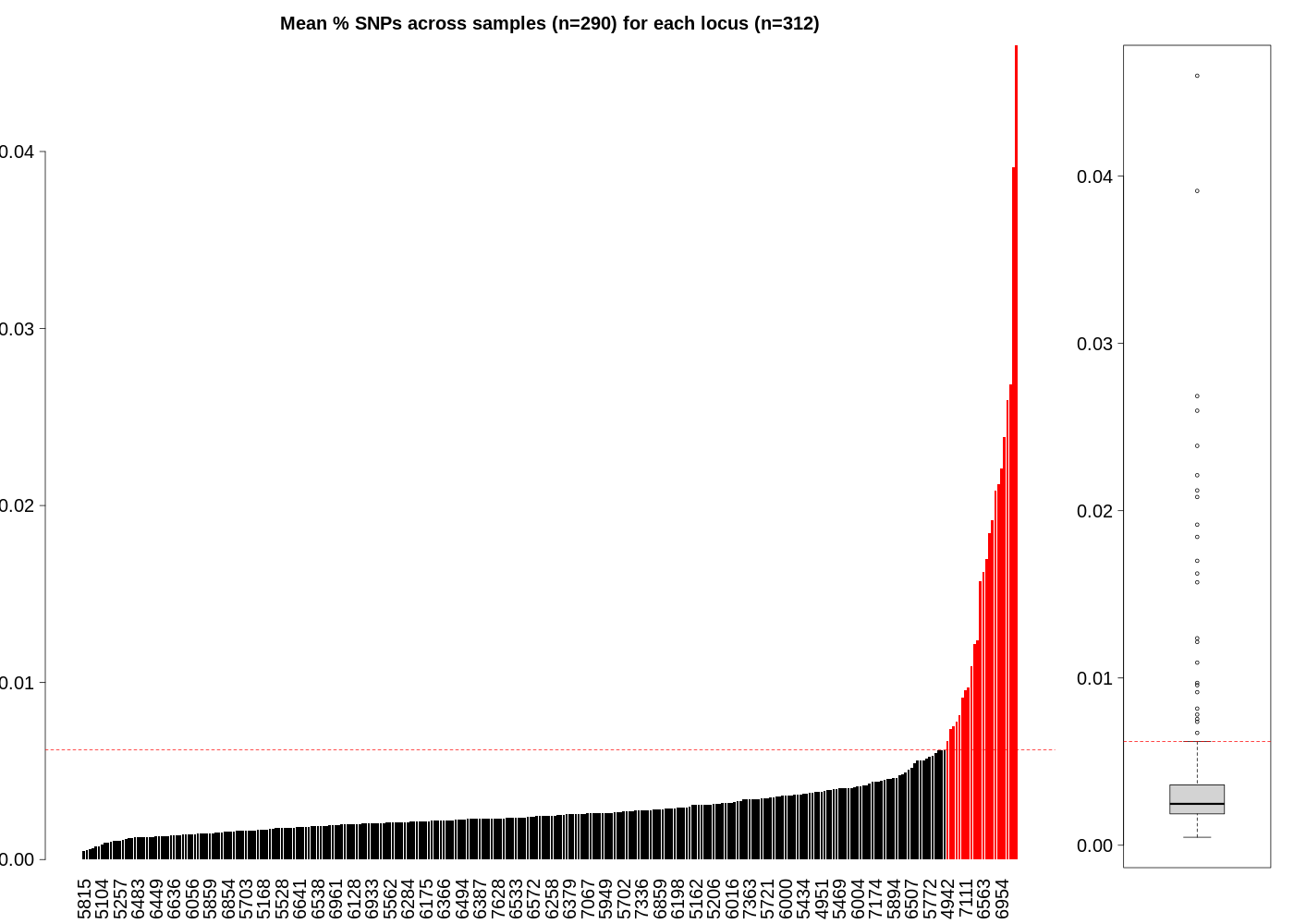

Variable 'remove_loci_for_all_samples_with_more_than_this_mean_proportion_of_SNPs' set to: outliers. Resulting threshold value (mean proportion of SNPs) was 0.00621 and 24 loci were removed.

Table 1: Removed loci for all species.

| locus | mean_prop_SNPs |
| --- | --- |
| 6412 | 0.046 |
| 6227 | 0.0391 |
| 5968 | 0.0269 |
| 4471 | 0.026 |
| 6679 | 0.0239 |
| 6954 | 0.0221 |
| 4757 | 0.0212 |
| 5899 | 0.0208 |
| 5865 | 0.0192 |
| 4806 | 0.0184 |
| 6487 | 0.017 |
| 6563 | 0.0162 |
| 6404 | 0.0157 |
| 7367 | 0.0124 |
| 6038 | 0.0122 |
| 6274 | 0.0109 |
| 5870 | 0.0097 |
| 7111 | 0.0096 |
| 6883 | 0.0092 |
| 5347 | 0.0082 |
| 5943 | 0.0078 |
| 6373 | 0.0075 |
| 6488 | 0.0074 |
| 4942 | 0.0067 |

**Paralogs removed for each sample**

Figure 2. Boxplots of the distribution of values for SNPs of each locus for each sample. Outlier loci are shown as dots.

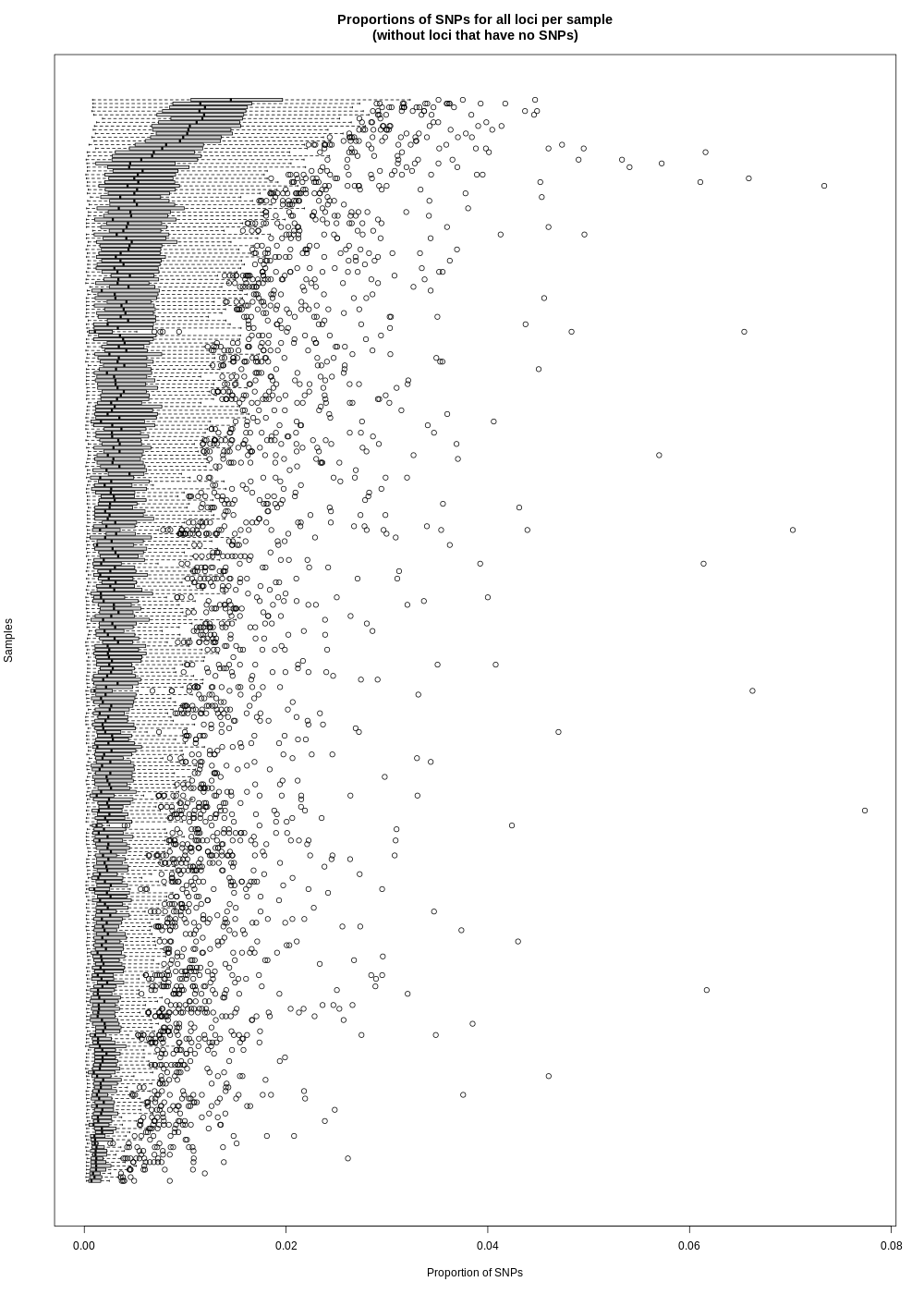

Table 2: List of putative paralog removal for each sample including threshold value (proportion of SNPs), number of loci removed, and names or removed loci.

| Sample | Threshold | # removed | Name of removed loci |
| --- | --- | --- | --- |
| Nepenthes_alata_G07767 | 0.00918 | 10 | 5502, 5822, 5921, 5945, 5974, 6114, 6797, 6854, 7021, 7628 |
| Nepenthes_alata_G07886 | 0.0094 | 10 | 4724, 5464, 5644, 5945, 5974, 6004, 6746, 6779, 6797, 6962 |
| Nepenthes_albomarginata_G08129 | 0.01322 | 12 | 4989, 5502, 5772, 5926, 5942, 6130, 6216, 6420, 6460, 6779, 6792, 6947 |
| Nepenthes_albomarginata_G08095 | 0.01351 | 5 | 5926, 6507, 6792, 6924, 7174 |
| Nepenthes_ampullaria_G07903 | 0.02397 | 7 | 5463, 5513, 6048, 6130, 6527, 6995, 7331 |
| Nepenthes_ampullaria_G08093 | 0.02134 | 9 | 5531, 5620, 6048, 6130, 6432, 6528, 6782, 6848, 7028 |
| Nepenthes_ampullaria_G08134 | 0.01938 | 9 | 6048, 6130, 6393, 6407, 6527, 6848, 6909, 6962, 7028 |
| Nepenthes_ampullaria_G08137 | 0.01885 | 6 | 4890, 5513, 6527, 6782, 6913, 7331 |
| Nepenthes_ampullaria_x_gracilis_G08101 | 0.02843 | 8 | 4848, 6048, 6462, 6528, 6962, 7067, 7273, 7331 |
| Nepenthes_ampullaria_x_tobaica_G08098 | 0.02876 | 7 | 4848, 5469, 5772, 5921, 6733, 6909, 7029 |
| Nepenthes_ampullaria_x_rafflesiana_G08104 | 0.02406 | 12 | 4989, 5513, 6004, 6130, 6420, 6432, 6527, 6746, 6848, 6962, 7028, 7331 |
| Nepenthes_attenboroughii_G08117 | 0.00757 | 16 | 5333, 5644, 5772, 5821, 5849, 6036, 6064, 6303, 6405, 6462, 6532, 6649, 6733, 6746, 6865, 6955 |
| Nepenthes_bellii_G08102 | 0.02872 | 7 | 5220, 6299, 6420, 6462, 6733, 6797, 6909 |
| Nepenthes_benstonei_G07897 | 0.01631 | 5 | 4796, 5469, 6528, 7028, 7135 |
| Nepenthes_bicalcarata_G08130 | 0.01247 | 6 | 5138, 5206, 5531, 6072, 6797, 7363 |
| Nepenthes_bicalcarata_x_ampullaria_G08132 | 0.02264 | 8 | 4989, 5090, 5138, 5842, 7028, 7067, 7174, 7331 |
| Nepenthes_bokorensis_G07856 | 0.00757 | 10 | 4802, 4951, 5398, 5460, 5536, 5921, 6284, 6528, 6909, 7583 |
| Nepenthes_bokorensis_x_ventricosa_G07899 | 0.02618 | 2 | 6713, 6733 |
| Nepenthes_bongso_G07898 | 0.0128 | 9 | 4724, 4890, 4951, 5032, 5162, 5489, 5562, 5921, 7174 |
| Nepenthes_borneensis_G07870 | 0.00626 | 18 | 4691, 5138, 5188, 5639, 5936, 5950, 6068, 6130, 6258, 6299, 6405, 6492, 6792, 6865, 6913, 6962, 7279, 7336 |
| Nepenthes_boschiana_G07855 | 0.00821 | 10 | 4691, 5188, 5463, 5816, 5950, 6072, 6383, 6492, 6779, 6913 |
| Nepenthes_burkei_x_ventricosa_G07877 | 0.00939 | 20 | 4848, 5032, 5131, 5138, 5162, 5463, 5477, 5822, 5910, 5940, 6064, 6068, 6216, 6299, 6459, 6570, 6825, 6860, 7363, 7577 |
| Nepenthes_boschiana_G08112 | 0.01129 | 4 | 4691, 5859, 6746, 6792 |
| Nepenthes_boschiana_x_glandulifera_G07862 | 0.01527 | 8 | 5427, 5502, 5531, 5821, 6298, 6507, 6865, 7273 |
| Nepenthes_burkei_G07875 | 0.00793 | 8 | 5842, 5858, 5910, 5945, 5958, 6198, 6216, 6459 |
| Nepenthes_burkei_x_ventricosa_G07783 | 0.01145 | 13 | 4848, 5032, 5822, 5842, 5910, 5936, 5940, 6299, 6389, 6459, 6620, 6792, 6825 |
| Nepenthes_burkei_x_ventricosa_G07854 | 0.01174 | 13 | 5463, 5578, 5822, 5894, 5910, 5936, 6064, 6299, 6459, 6620, 6649, 6792, 7136 |
| Nepenthes_sp_calcicola_G08235 | 0.01225 | 9 | 5427, 5702, 5721, 5936, 6056, 6130, 6685, 6860, 7028 |
| Nepenthes_campanulata_G08136 | 0.00674 | 6 | 4796, 4802, 5469, 6449, 6500, 6733 |
| Nepenthes_chaniana_G07882 | 0.01121 | 7 | 4691, 5335, 5656, 6130, 6528, 6792, 6955 |
| Nepenthes_clipeata_x_ventricosa_G08106 | 0.02745 | 6 | 5343, 5531, 6459, 6733, 6933, 7136 |
| Nepenthes_copelandii_G07863 | 0.00784 | 3 | 5463, 6034, 6860 |
| Nepenthes_copelandii_G07883 | 0.00814 | 8 | 4796, 5123, 5477, 6540, 6649, 6797, 6913, 6962 |
| Nepenthes_alata_ERR3657191 | 0.01001 | 6 | 5138, 5620, 5949, 6041, 6216, 6528 |
| Nepenthes_alba_ERR3657192 | 0.01436 | 4 | 5366, 5949, 6004, 6962 |
| Nepenthes_albomarginata_ERR3657194 | 0.01045 | 3 | 5926, 6072, 6528 |
| Nepenthes_albomarginata_ERR3662876 | 0.01171 | 6 | 5449, 5770, 5816, 5926, 6282, 6420 |
| Nepenthes_ampullaria_ERR3657200 | 0.02082 | 4 | 5477, 6130, 6420, 6528 |
| Nepenthes_ampullaria_ERR3659452 | 0.01945 | 12 | 5477, 6004, 6130, 6432, 6848, 6913, 6962, 6992, 7028, 7135, 7174, 7331 |
| Nepenthes_ampullaria_ERR3660988 | 0.01828 | 9 | 5528, 5620, 6130, 6527, 6528, 6848, 6913, 7028, 7331 |
| Nepenthes_ampullaria_ERR3662930 | 0.01761 | 2 | 6528, 7028 |
| Nepenthes_andamana_ERR3662060 | 0.01435 | 8 | 4527, 5502, 5821, 5921, 6258, 6298, 6492, 7324 |
| Nepenthes_angasanensis_ERR3712174 | 0.00962 | 11 | 4889, 5162, 5206, 5460, 5464, 5620, 5642, 5744, 5894, 6004, 6620 |
| Nepenthes_angustifolia_ERR3712173 | 0.03327 | 3 | 5772, 6000, 6462 |
| Nepenthes_aristolochioides_ERR3662061 | 0.00976 | 10 | 5188, 5843, 6068, 6216, 6284, 6383, 6527, 6572, 6848, 6992 |
| Nepenthes_armin_ERR3712097 | 0.00744 | 12 | 4932, 5200, 5463, 5502, 5921, 6003, 6462, 6528, 6825, 6924, 6979, 7324 |
| Nepenthes_attenboroughii_ERR3712098 | 0.00804 | 12 | 5299, 5644, 5945, 5950, 6003, 6303, 6389, 6528, 6955, 7174, 7363, 7371 |
| Nepenthes_beccariana_ERR3660989 | 0.00968 | 8 | 5428, 5464, 6376, 6384, 6454, 6962, 7028, 7279 |
| Nepenthes_bellii_ERR3662062 | 0.00879 | 5 | 5469, 5594, 6216, 6528, 6909 |
| Nepenthes_benstonei_ERR3662142 | 0.01505 | 6 | 5463, 5594, 5744, 6450, 7028, 7128 |
| Nepenthes_biak_ERR3712175 | 0.00366 | 10 | 5463, 5551, 5744, 5921, 5940, 6098, 6284, 6717, 6913, 7363 |
| Nepenthes_bicalcarata_ERR3657202 | 0.01092 | 6 | 5138, 5428, 5463, 6175, 6532, 6797 |
| Nepenthes_bicalcarata_ERR3660990 | 0.01076 | 5 | 4724, 5138, 5531, 5772, 6072 |
| Nepenthes_bicalcarata_ERR3672060 | 0.00775 | 5 | 5138, 5162, 5206, 5594, 6072 |
| Nepenthes_bokorensis_ERR3662143 | 0.009 | 5 | 5502, 5821, 5949, 7028, 7324 |
| Nepenthes_bongso_ERR3662063 | 0.00923 | 10 | 5299, 5477, 5502, 5594, 5721, 5960, 6216, 6383, 6860, 6968 |
| Nepenthes_boschiana_ERR3662144 | 0.01278 | 3 | 4691, 4989, 6439 |
| Nepenthes_burbidgeae_ERR3662064 | 0.00671 | 7 | 4954, 5404, 5421, 5594, 5974, 6000, 6379 |
| Nepenthes_burkei_ERR3662065 | 0.01044 | 3 | 5469, 7028, 7577 |
| Nepenthes_campanulata_ERR3662145 | 0.00674 | 5 | 5454, 5469, 5578, 6946, 7279 |
| Nepenthes_ceciliae_ERR3662066 | 0.00884 | 6 | 5318, 6299, 6660, 6713, 6746, 6860 |
| Nepenthes_chang_ERR3712005 | 0.00927 | 5 | 4951, 5421, 5477, 6004, 7028 |
| Nepenthes_chaniana_ERR3662277 | 0.01575 | 7 | 4989, 5427, 5463, 6439, 6462, 6528, 7135 |
| Nepenthes_clipeata_ERR3662278 | 0.02114 | 0 |  |
| Nepenthes_copelandii_ERR3662279 | 0.00732 | 7 | 4848, 5463, 5721, 5791, 5919, 6034, 6746 |
| Nepenthes_cornuta_ERR3712006 | 0.0091 | 6 | 5919, 6034, 6299, 6552, 6660, 7602 |
| Nepenthes_danseri_ERR3672061 | 0.01268 | 5 | 4691, 5188, 5821, 6979, 6992 |
| Nepenthes_deaniana_ERR3712177 | 0.0122 | 11 | 5469, 5821, 6003, 6454, 6528, 6531, 6909, 6962, 7136, 7324, 7577 |
| Nepenthes_densiflora_ERR3662067 | 0.01066 | 8 | 5038, 5162, 5594, 5821, 5926, 6004, 6532, 7324 |
| Nepenthes_diatas_ERR3662280 | 0.00901 | 8 | 5469, 5620, 5949, 6110, 6320, 6717, 6860, 7324 |
| Nepenthes_distillatoria_ERR3662281 | 0.00655 | 16 | 4890, 4951, 5299, 5578, 5821, 6034, 6050, 6130, 6216, 6450, 6507, 6528, 6639, 6685, 6860, 7371 |
| Nepenthes_dubia_ERR3657203 | 0.00985 | 5 | 4796, 5206, 6458, 6620, 7331 |
| Nepenthes_edwardsiana_ERR3660991 | 0.01098 | 2 | 6216, 6450 |
| Nepenthes_ephippiata_ERR3660992 | 0.01025 | 4 | 5772, 6383, 6393, 6913 |
| Nepenthes_eustachya_ERR3662068 | 0.01018 | 8 | 5404, 5464, 5536, 5642, 6198, 6552, 6689, 7324 |
| Nepenthes_eymae_ERR3662871 | 0.00421 | 7 | 5578, 6379, 6860, 6909, 7028, 7363, 7583 |
| Nepenthes_faizaliana_ERR3662069 | 0.01427 | 4 | 5772, 5919, 6528, 6962 |
| Nepenthes_flava_ERR3712178 | 0.01072 | 2 | 4890, 6746 |
| Nepenthes_glabrata_ERR3662070 | 0.00659 | 7 | 4989, 5428, 5551, 5840, 6003, 6492, 6848 |
| Nepenthes_glandulifera_ERR3662282 | 0.01824 | 7 | 4691, 5772, 5822, 6439, 6552, 6848, 7067 |
| Nepenthes_graciliflora_ERR3661586 | 0.00958 | 9 | 5463, 5502, 5721, 6003, 6462, 6825, 6924, 7324, 7371 |
| Nepenthes_graciliflora_ERR3662284 | 0.00723 | 10 | 4848, 5463, 5536, 5721, 5772, 6114, 6366, 6376, 6450, 6717 |
| Nepenthes_gracilis_ERR3657205 | 0.02481 | 3 | 6110, 6462, 6528 |
| Nepenthes_gracilis_ERR3662283 | 0.02122 | 3 | 6000, 6962, 6968 |
| Nepenthes_gracillima_ERR3662285 | 0.01614 | 3 | 5064, 5090, 6909 |
| Nepenthes_gymnamphora_ERR3662071 | 0.00543 | 12 | 5326, 5366, 5434, 5857, 5949, 5960, 6000, 6003, 6492, 6507, 6660, 7628 |
| Nepenthes_gymnamphora_ERR3662923 | 0.00587 | 3 | 4989, 5656, 7602 |
| Nepenthes_hamata_ERR3657206 | 0.00676 | 4 | 5469, 5620, 6282, 6860 |
| Nepenthes_hamata_ERR3657207 | 0.00415 | 4 | 5551, 5594, 5620, 5821 |
| Nepenthes_hamiguitanensis_ERR3657208 | 0.00575 | 6 | 4691, 5702, 5772, 6284, 6462, 6528 |
| Nepenthes_hemsleyana_ERR3662286 | 0.01584 | 14 | 5123, 5430, 5513, 5772, 5960, 6003, 6072, 6299, 6303, 6420, 6492, 6649, 6924, 7174 |
| Nepenthes_hirsuta_ERR3662072 | 0.0168 | 6 | 6000, 6492, 6532, 6746, 6962, 7324 |
| Nepenthes_hispida_ERR3712007 | 0.0073 | 5 | 6176, 6528, 6746, 6992, 7324 |
| Nepenthes_inermis_ERR3662073 | 0.00955 | 2 | 5721, 6320 |
| Nepenthes_insignis_ERR3661587 | 0.00354 | 3 | 5816, 5821, 6717 |
| Nepenthes_izumiae_ERR3662287 | 0.01233 | 5 | 4796, 5206, 5620, 5949, 6041 |
| Nepenthes_jacquelineae_ERR3662074 | 0.01227 | 4 | 5960, 5974, 6462, 7577 |
| Nepenthes_jamban_ERR3662075 | 0.00945 | 3 | 5138, 5406, 6051 |
| Nepenthes_justinae_ERR3712099 | 0.00636 | 6 | 4691, 4848, 5335, 6004, 6439, 6860 |
| Nepenthes_kampotiana_ERR3657987 | 0.00701 | 12 | 4951, 5188, 5702, 5744, 6003, 6639, 6667, 6733, 6913, 6962, 7174, 7279 |
| Nepenthes_kerrii_ERR3712008 | 0.0117 | 1 |  |
| Nepenthes_khasiana_ERR3662076 | 0.00557 | 9 | 4951, 5162, 5428, 5463, 5469, 5849, 5922, 5977, 6685 |
| Nepenthes_klossii_ERR3661588 | 0.00315 | 2 | 6003, 7028 |
| Nepenthes_kongkandana_ERR3662288 | 0.0157 | 11 | 5463, 5469, 5733, 5744, 5894, 5974, 6004, 6378, 6462, 6792, 6962 |
| Nepenthes_leonardoi_ERR3712100 | 0.00861 | 8 | 5188, 5469, 5772, 5849, 6216, 6717, 7324, 7577 |
| Nepenthes_lingulata_ERR3662078 | 0.01262 | 3 | 5974, 6320, 6492 |
| Nepenthes_longifolia_ERR3662289 | 0.00964 | 5 | 5428, 5642, 6492, 6528, 6685 |
| Nepenthes_lowii_ERR3662085 | 0.01514 | 6 | 4989, 5299, 6947, 6962, 7313, 7363 |
| Nepenthes_macfarlanei_ERR3662290 | 0.01411 | 4 | 5642, 5821, 6175, 7135 |
| Nepenthes_macrophylla_ERR3662086 | 0.00863 | 13 | 5463, 5477, 5721, 5772, 6068, 6216, 6258, 6462, 6492, 6933, 6992, 7313, 7363 |
| Nepenthes_macrovulgaris_ERR3657988 | 0.01084 | 4 | 5460, 5644, 6003, 6860 |
| Nepenthes_madagascariensis_ERR3672062 | 0.00787 | 5 | 4724, 4951, 5138, 6652, 6924 |
| Nepenthes_mantalingajanensis_ERR3712101 | 0.00662 | 13 | 4691, 4932, 5434, 5464, 5477, 5644, 5702, 6528, 6532, 6685, 6909, 6913, 7628 |
| Nepenthes_mapuluensis_ERR3662087 | 0.01462 | 12 | 4848, 5018, 5464, 5469, 5894, 6393, 6528, 6660, 6746, 6962, 7324, 7336 |
| Nepenthes_masoalensis_ERR3662088 | 0.00653 | 11 | 4691, 4724, 4889, 4951, 5064, 5138, 5721, 6041, 6130, 6685, 6913 |
| Nepenthes_maxima_ERR3661602 | 0.0062 | 11 | 4691, 5123, 5355, 5578, 5772, 6303, 6439, 6507, 6572, 6733, 6909 |
| Nepenthes_maxima_ERR3661604 | 0.00544 | 10 | 4691, 5578, 5620, 6299, 6405, 6572, 6882, 6909, 6978, 7583 |
| Nepenthes_maxima_ERR3661605 | 0.00343 | 5 | 5348, 6717, 6860, 7028, 7363 |
| Nepenthes_maxima_ERR3661607 | 0.0062 | 12 | 5299, 5366, 5464, 5502, 5578, 5620, 6000, 6051, 6379, 6860, 6909, 7336 |
| Nepenthes_maxima_ERR3661608 | 0.00464 | 13 | 4691, 5318, 5326, 5404, 5463, 5578, 5620, 5721, 5744, 5843, 5894, 6379, 6620 |
| Nepenthes_maxima_ERR3662875 | 0.00377 | 10 | 4527, 5138, 5348, 5721, 5816, 5822, 6320, 6379, 6860, 7363 |
| Nepenthes_merrilliana_ERR3662291 | 0.00766 | 4 | 5502, 6216, 6376, 6717 |
| Nepenthes_micramphora_ERR3657989 | 0.0055 | 4 | 5770, 6000, 6034, 6376 |
| Nepenthes_mikei_ERR3662089 | 0.01066 | 6 | 5477, 5578, 5642, 5821, 6003, 6962 |
| Nepenthes_mindanaoensis_ERR3661609 | 0.00827 | 5 | 5816, 6072, 6432, 6860, 7363 |
| Nepenthes_minima_ERR3662292 | 0.00578 | 8 | 5138, 5348, 5578, 5620, 6320, 6379, 6713, 7363 |
| Nepenthes_mirabilis_ERR3657198 | 0.01223 | 1 |  |
| Nepenthes_mirabilis_ERR3657199 | 0.01422 | 4 | 6303, 6532, 6782, 7067 |
| Nepenthes_mirabilis_ERR3661610 | 0.01348 | 13 | 4932, 4951, 5355, 5502, 5772, 6072, 6130, 6432, 6528, 6620, 6848, 7029, 7313 |
| Nepenthes_mirabilis_ERR3661611 | 0.01583 | 9 | 5206, 5816, 6000, 6532, 6848, 6860, 6946, 6962, 7028 |
| Nepenthes_mirabilis_ERR3661613 | 0.01658 | 5 | 5138, 5770, 6378, 7029, 7174 |
| Nepenthes_mirabilis_ERR3661615 | 0.01637 | 3 | 4848, 6909, 6913 |
| Nepenthes_mirabilis_ERR3661616 | 0.01133 | 9 | 4724, 4848, 5162, 5366, 5950, 6098, 6378, 6528, 6859 |
| Nepenthes_mirabilis_ERR3661617 | 0.01148 | 4 | 6378, 6913, 7241, 7324 |
| Nepenthes_mirabilis_ERR3661618 | 0.01731 | 1 |  |
| Nepenthes_mirabilis_ERR3661619 | 0.01509 | 4 | 4848, 6620, 6667, 6909 |
| Nepenthes_mirabilis_ERR3661620 | 0.01575 | 7 | 5950, 6383, 6450, 6527, 6532, 6667, 6909 |
| Nepenthes_mirabilis_ERR3661621 | 0.01562 | 4 | 4848, 6378, 6528, 6909 |
| Nepenthes_mirabilis_ERR3661622 | 0.01412 | 2 | 6528, 6909 |
| Nepenthes_mirabilis_ERR3662877 | 0.01777 | 6 | 5427, 5513, 5594, 6528, 6955, 7029 |
| Nepenthes_mirabilis_ERR3662928 | 0.01311 | 1 |  |
| Nepenthes_mirabilis_ERR3662929 | 0.01562 | 9 | 5449, 5744, 5770, 6072, 6303, 6620, 6848, 7028, 7067 |
| Nepenthes_mirabilis_ERR3672063 | 0.014 | 3 | 6016, 6420, 7029 |
| Nepenthes_mirabilis_var_echinostoma_ERR3657204 | 0.02241 | 5 | 5936, 6299, 6393, 6962, 7029 |
| Nepenthes_mira_ERR3662090 | 0.00883 | 8 | 4527, 5502, 5821, 6528, 6685, 6717, 7324, 7577 |
| Nepenthes_mollis_ERR3712179 | 0.01273 | 11 | 4848, 4989, 5299, 5355, 5699, 5926, 5960, 6004, 6492, 6507, 6601 |
| Nepenthes_monticola_ERR3712102 | 0.00318 | 5 | 5427, 5454, 6041, 6924, 7028 |
| Nepenthes_muluensis_ERR3657371 | 0.01333 | 4 | 5366, 5821, 6016, 6860 |
| Nepenthes_murudensis_ERR3662293 | 0.01252 | 5 | 5620, 5772, 6000, 6492, 6992 |
| Nepenthes_naga_ERR3662294 | 0.01124 | 7 | 5355, 5513, 5894, 5950, 6746, 6882, 7174 |
| Nepenthes_neoguineensis_ERR3657378 | 0.00528 | 4 | 5702, 5821, 6559, 7363 |
| Nepenthes_neoguineensis_ERR3662053 | 0.00695 | 5 | 5404, 6303, 6685, 7028, 7324 |
| Nepenthes_northiana_ERR3662295 | 0.0074 | 4 | 5894, 5977, 6216, 6303 |
| Nepenthes_oblanceolata_ERR3712103 | 0.00227 | 5 | 4691, 4954, 5464, 6717, 7028 |
| Nepenthes_ovata_ERR3662296 | 0.00757 | 3 | 5974, 6051, 6320 |
| Nepenthes_palawanensis_ERR3712010 | 0.00993 | 11 | 4951, 5177, 5404, 5469, 5551, 5554, 5772, 6003, 6320, 6383, 6528 |
| Nepenthes_papuana_ERR3657379 | 0.00615 | 11 | 4691, 5463, 5772, 5841, 5958, 6572, 6685, 6746, 6979, 7028, 7336 |
| Nepenthes_peltata_ERR3662297 | 0.00835 | 8 | 4691, 5463, 5977, 6299, 6649, 6746, 6914, 7028 |
| Nepenthes_pervillei_ERR3657380 | 0.0061 | 7 | 5138, 5299, 5477, 5821, 6068, 6216, 7313 |
| Nepenthes_petiolata_ERR3662091 | 0.01038 | 3 | 5859, 5977, 7028 |
| Nepenthes_philippinensis_ERR3662299 | 0.01313 | 8 | 5162, 5264, 5366, 5840, 6454, 6531, 6717, 6746 |
| Nepenthes_pitopangii_ERR3712011 | 0.00602 | 2 | 5428, 5840 |
| Nepenthes_platychila_ERR3662300 | 0.01738 | 8 | 4691, 4989, 5366, 5469, 5578, 5772, 6507, 6528 |
| Nepenthes_pulchra_ERR3657381 | 0.00743 | 7 | 4848, 5090, 5489, 5531, 5942, 6860, 7363 |
| Nepenthes_rafflesiana_ERR3657382 | 0.0097 | 7 | 4691, 5206, 5531, 6432, 6746, 7028, 7029 |
| Nepenthes_rafflesiana_ERR3657383 | 0.01264 | 7 | 4989, 5430, 5721, 6004, 6432, 6882, 7028 |
| Nepenthes_rafflesiana_ERR3662092 | 0.01638 | 9 | 5123, 5343, 5449, 5721, 6432, 6848, 7028, 7029, 7363 |
| Nepenthes_rajah_ERR3657984 | 0.00561 | 1 |  |
| Nepenthes_ramispina_ERR3662301 | 0.01179 | 16 | 4802, 5280, 5299, 5434, 5513, 6003, 6282, 6299, 6538, 6559, 6570, 6782, 6875, 6909, 6914, 7029 |
| Nepenthes_reinwardtiana_ERR3662093 | 0.01951 | 9 | 5032, 5463, 5644, 5772, 5926, 6439, 6825, 6909, 6914 |
| Nepenthes_rhombicaulis_ERR3662302 | 0.00884 | 10 | 4527, 4889, 5744, 5849, 5950, 6405, 6782, 6825, 7174, 7324 |
| Nepenthes_rigidifolia_ERR3712180 | 0.01331 | 15 | 4848, 4893, 4951, 5304, 5428, 5502, 5949, 5974, 6130, 6531, 6782, 6848, 6962, 7174, 7331 |
| Nepenthes_robcantleyi_ERR3712181 | 0.0044 | 9 | 4691, 5220, 5642, 5913, 5919, 5936, 6792, 7135, 7336 |
| Nepenthes_rowaniae_ERR3662094 | 0.01722 | 3 | 5138, 6072, 6528 |
| Nepenthes_sanguinea_ERR3662095 | 0.01255 | 12 | 5460, 5531, 5744, 6130, 6462, 6782, 6848, 6865, 6909, 6914, 6946, 6992 |
| Nepenthes_sibuyanensis_ERR3662096 | 0.00698 | 4 | 5123, 5138, 5940, 6366 |
| Nepenthes_singalana_ERR3662303 | 0.01019 | 13 | 4890, 5038, 5502, 5702, 5849, 5949, 5960, 5974, 6216, 6363, 6527, 6860, 7331 |
| Nepenthes_smilesii_ERR3662924 | 0.01362 | 8 | 5206, 5421, 5821, 6003, 6004, 6298, 6462, 7324 |
| Nepenthes_smilesii_ERR3662925 | 0.01186 | 9 | 5460, 5464, 6003, 6004, 6130, 6298, 6620, 6660, 6733 |
| Nepenthes_sp_adrianii_ERR3712004 | 0.01058 | 4 | 4890, 5434, 5599, 7331 |
| Nepenthes_sp_Anipahan_ERR3712104 | 0.00744 | 17 | 4527, 4951, 5188, 5264, 5551, 5894, 5913, 5940, 6051, 6198, 6689, 6860, 6913, 7174, 7333, 7371, 7577 |
| Nepenthes_spathulata_ERR3662304 | 0.00846 | 5 | 5206, 5463, 6068, 6110, 6216 |
| Nepenthes_sp_BM-2019_ERR3712119 | 0.01063 | 10 | 4691, 4802, 4951, 5206, 5562, 5599, 5974, 6003, 6649, 7174 |
| Nepenthes_spectabilis_ERR3657985 | 0.00806 | 3 | 5206, 5949, 6041 |
| Nepenthes_stenophylla_ERR3662305 | 0.01273 | 10 | 5463, 5469, 5477, 5960, 6393, 6420, 6792, 6848, 6913, 7324 |
| Nepenthes_sumagaya_ERR3712012 | 0.00964 | 3 | 5721, 6072, 6713 |
| Nepenthes_suratensis_ERR3712013 | 0.01583 | 5 | 4890, 5299, 5919, 6303, 7324 |
| Nepenthes_surigaoensis_ERR3662315 | 0.00816 | 10 | 5123, 5463, 5477, 5843, 6216, 6494, 6528, 6825, 6848, 7577 |
| Nepenthes_talangensis_ERR3662326 | 0.00885 | 12 | 5428, 5502, 5849, 5945, 5949, 5960, 5974, 6068, 6216, 6320, 6913, 6924 |
| Nepenthes_tenax_ERR3662327 | 0.01521 | 3 | 6034, 6527, 6913 |
| Nepenthes_tentaculata_ERR3662056 | 0.01088 | 3 | 5177, 6528, 6746 |
| Nepenthes_tentaculata_ERR3662328 | 0.00741 | 12 | 4989, 5264, 5578, 5620, 5772, 5974, 6198, 6601, 6860, 6924, 7313, 7331 |
| Nepenthes_tenuis_ERR3662329 | 0.00947 | 7 | 5162, 5894, 6216, 6284, 6859, 7174, 7628 |
| Nepenthes_thai_ERR3657986 | 0.01422 | 0 |  |
| Nepenthes_thorelii_ERR3662097 | 0.0074 | 6 | 4951, 5398, 5949, 5977, 6462, 7135 |
| Nepenthes_tobaica_ERR3662098 | 0.00946 | 10 | 5477, 5551, 5921, 5960, 6016, 6114, 6216, 6320, 6528, 6550 |
| Nepenthes_tobaica_ERR3662099 | 0.00981 | 5 | 5594, 5620, 6068, 6947, 6962 |
| Nepenthes_tomoriana_ERR3662330 | 0.01079 | 7 | 5264, 5428, 5721, 6528, 6860, 6979, 7324 |
| Nepenthes_treubiana_ERR3657197 | 0.01042 | 8 | 5578, 6003, 6130, 6303, 6636, 6685, 6979, 7324 |
| Nepenthes_treubiana_ERR3662100 | 0.011 | 10 | 4890, 5427, 6003, 6048, 6378, 6393, 6601, 6636, 6685, 7324 |
| Nepenthes_truncata_ERR3662057 | 0.00445 | 7 | 5335, 5536, 5721, 6299, 6865, 6914, 7363 |
| Nepenthes_veitchii_ERR3662101 | 0.01586 | 5 | 5463, 6298, 6450, 6848, 6909 |
| Nepenthes_ventricosa_ERR3672064 | 0.00684 | 5 | 4527, 5940, 6198, 6393, 6601 |
| Nepenthes_villosa_ERR3662058 | 0.00591 | 6 | 4992, 5463, 5772, 6068, 6825, 7363 |
| Nepenthes_vogelii_ERR3672065 | 0.01079 | 9 | 4691, 5744, 5770, 5821, 5918, 6393, 6462, 6507, 6979 |
| Nepenthes_rafflesiana_x_ampullaria_ERR3662102 | 0.02816 | 12 | 4989, 5477, 5853, 6004, 6048, 6130, 6299, 6420, 6528, 6733, 6782, 6848 |
| Nepenthes_xiphioides_ERR3662331 | 0.00565 | 6 | 5772, 5980, 6130, 6216, 6531, 6913 |
| Nepenthes_ephippiata_G07857 | 0.01309 | 5 | 5464, 5469, 6450, 6507, 6848 |
| Nepenthes_ephippiata_G07900 | 0.00884 | 2 | 5355, 6393 |
| Nepenthes_eymae_G07887 | 0.00649 | 16 | 4691, 5721, 5815, 5977, 5980, 6051, 6098, 6130, 6303, 6363, 6393, 6507, 6532, 6860, 6909, 7028 |
| Nepenthes_eymae_G08121 | 0.00611 | 10 | 4691, 5578, 5822, 6379, 6507, 6540, 6631, 6860, 6909, 6955 |
| Nepenthes_faizaliana_G08090 | 0.01103 | 6 | 5449, 5744, 5772, 5936, 6420, 6947 |
| Nepenthes_lowii_x_campanulata_G08109 | 0.02194 | 13 | 5463, 5469, 5772, 5894, 6000, 6378, 6450, 6528, 6649, 6733, 6779, 6792, 6865 |
| Nepenthes_flava_G08119 | 0.01165 | 9 | 4890, 5138, 5620, 5949, 5960, 5974, 6462, 6527, 6909 |
| Nepenthes_zakriana_G08120 | 0.01567 | 7 | 4848, 5821, 6000, 6507, 6792, 7363, 7572 |
| Nepenthes_glandulifera_G07830 | 0.01524 | 6 | 4691, 5123, 5772, 5821, 6130, 6528 |
| Nepenthes_gracilis_G08091 | 0.02278 | 2 | 5357, 6130 |
| Nepenthes_gymnamphora_G07893 | 0.01029 | 6 | 5032, 5822, 5949, 5960, 6029, 6527 |
| Nepenthes_gymnamphora_G07901 | 0.01123 | 0 |  |
| Nepenthes_izumiae_G07891 | 0.00941 | 9 | 4691, 5296, 5428, 5477, 5721, 5960, 6003, 6460, 6492 |
| Nepenthes_izumiae_x_ventricosa_G07866 | 0.02616 | 10 | 5123, 5304, 5463, 5469, 5822, 5894, 5940, 6003, 6299, 6848 |
| Nepenthes_jacquelineae_G07873 | 0.00859 | 8 | 4802, 5562, 5843, 6532, 6782, 6955, 7028, 7174 |
| Nepenthes_kampotiana_G07895 | 0.0277 | 11 | 5772, 5866, 5921, 5936, 6198, 6462, 6552, 6733, 6848, 6909, 7067 |
| Nepenthes_khasiana_G07874 | 0.00448 | 17 | 5162, 5299, 5430, 5463, 5469, 5477, 5922, 5936, 6068, 6130, 6384, 6420, 6454, 6507, 6533, 6685, 7324 |
| Nepenthes_kongkandana_G07869 | 0.00993 | 16 | 4848, 4890, 5220, 5449, 5821, 5849, 5913, 5919, 5921, 5949, 6238, 6432, 6924, 6962, 7174, 7583 |
| Nepenthes_maxima_G07859 | 0.00633 | 14 | 4691, 5578, 5772, 5942, 5977, 6303, 6507, 6526, 6848, 6860, 7028, 7135, 7336, 7363 |
| Nepenthes_maxima_G07861 | 0.00807 | 18 | 4691, 5318, 5335, 5348, 5721, 5772, 5816, 6004, 6130, 6379, 6384, 6507, 6540, 6713, 6860, 7135, 7136, 7583 |
| Nepenthes_maxima_G07864 | 0.00636 | 22 | 4691, 5200, 5335, 5348, 5578, 5620, 5702, 5721, 5770, 5772, 5822, 5942, 6004, 6320, 6379, 6447, 6649, 6667, 6860, 6882, 6995, 7135 |
| Nepenthes_maxima_G08103 | 0.0055 | 13 | 4691, 5168, 5578, 5772, 5942, 5977, 5980, 6303, 6507, 6848, 7028, 7336, 7363 |
| Nepenthes_maxima_G08107 | 0.00602 | 16 | 4691, 5168, 5348, 5578, 5772, 5913, 5942, 5977, 6303, 6507, 6738, 6848, 7028, 7135, 7336, 7363 |
| Nepenthes_merrilliana_G08089 | 0.00887 | 12 | 5123, 5304, 5463, 5721, 5958, 6226, 6393, 6636, 6641, 6848, 7136, 7363 |
| Nepenthes_minima_G07872 | 0.00767 | 13 | 4691, 5348, 5464, 5578, 5620, 5894, 6004, 6051, 6130, 6320, 6507, 6667, 6909 |
| Nepenthes_mirabilis_G08086 | 0.00839 | 14 | 4691, 4889, 4951, 5304, 6003, 6034, 6051, 6072, 6378, 6458, 6601, 6639, 7174, 7577 |
| Nepenthes_mirabilis_G08099 | 0.01748 | 17 | 5469, 5513, 5594, 5894, 5980, 6003, 6320, 6378, 6384, 6420, 6439, 6528, 6746, 6848, 6865, 6909, 6914 |
| Nepenthes_mirabilis_G08236 | 0.01076 | 8 | 4951, 5936, 6034, 6532, 6540, 6860, 6913, 7628 |
| Nepenthes_mirabilis_G08237 | 0.01295 | 3 | 5038, 6527, 6995 |
| Nepenthes_mirabilis_G08244 | 0.01117 | 5 | 4951, 5304, 5348, 6072, 6601 |
| Nepenthes_mirabilis_G08245 | 0.01126 | 4 | 4951, 5926, 6072, 7174 |
| Nepenthes_mirabilis_G08246 | 0.01121 | 3 | 4951, 5348, 5513 |
| Nepenthes_mirabilis_G08247 | 0.01079 | 3 | 4932, 6859, 6913 |
| Nepenthes_mirabilis_var_echinostoma_G08131 | 0.01783 | 11 | 5469, 5513, 5770, 6016, 6072, 6299, 6320, 6420, 6848, 6962, 7029 |
| Nepenthes_mirabilis_var_globosa_G08128 | 0.01664 | 10 | 4951, 5821, 5936, 5950, 6393, 6420, 6552, 6652, 6962, 7029 |
| Nepenthes_mirabilis_G08094 | 0.01722 | 15 | 4989, 5469, 5594, 5770, 5980, 6378, 6384, 6420, 6528, 6848, 6859, 6909, 6992, 7029, 7067 |
| Nepenthes_mirabilis_x_rowaniae_G08241 | 0.01364 | 0 |  |
| Nepenthes_mirabilis_x_tenax_G08242 | 0.01394 | 6 | 5348, 5936, 6527, 6601, 6865, 7174 |
| Nepenthes_mirabilis_x_tenax_G08243 | 0.01504 | 2 | 5980, 6527 |
| Nepenthes_mira_G07902 | 0.00811 | 5 | 4527, 5355, 5463, 5513, 5551 |
| Nepenthes_neoguineensis_G08125 | 0.00994 | 10 | 5318, 5744, 5822, 5894, 6130, 6303, 6320, 6685, 6860, 7028 |
| Nepenthes_northiana_G08135 | 0.0065 | 7 | 5463, 5857, 5977, 6003, 6130, 6216, 6303 |
| Nepenthes_ovata_G08118 | 0.00797 | 2 | 5032, 5355 |
| Nepenthes_parvula_G08240 | 0.01589 | 3 | 6034, 6527, 6532 |
| Nepenthes_parvula_G08248 | 0.00555 | 14 | 4691, 4951, 4989, 5038, 5280, 5578, 5772, 5981, 6034, 6393, 6860, 6913, 7363, 7583 |
| Nepenthes_pervillei_G08123 | 0.00767 | 18 | 4691, 5138, 5536, 5620, 5644, 5656, 5958, 5980, 6034, 6068, 6130, 6216, 6303, 6458, 6492, 6559, 6962, 7313 |
| Nepenthes_petiolata_G07881 | 0.00844 | 12 | 5138, 5428, 5464, 5477, 5821, 5960, 6528, 6746, 6914, 7028, 7331, 7336 |
| Nepenthes_philippinensis_G07878 | 0.01336 | 14 | 5123, 5304, 5449, 5551, 5918, 6383, 6526, 6532, 6552, 6559, 6649, 6865, 7174, 7331 |
| Nepenthes_rafflesiana_G08097 | 0.01649 | 8 | 4989, 5513, 5822, 6378, 6432, 6649, 6933, 7028 |
| Nepenthes_rafflesiana_G08127 | 0.01783 | 9 | 5477, 6299, 6303, 6407, 6432, 6532, 6848, 6924, 7028 |
| Nepenthes_rafflesiana_x_ampullaria_G08126 | 0.0247 | 7 | 5477, 5513, 6000, 6004, 6130, 6527, 7028 |
| Nepenthes_rajah_G08114 | 0.00545 | 6 | 4724, 5843, 6072, 6299, 6432, 6924 |
| Nepenthes_reinwardtiana_G08133 | 0.01946 | 8 | 4989, 5842, 5849, 6298, 6733, 6792, 6825, 7331 |
| Nepenthes_reinwardtiana_G08100 | 0.02258 | 7 | 5463, 6303, 6439, 6733, 6909, 6933, 7331 |
| Nepenthes_robcantleyi_G08111 | 0.00468 | 4 | 4691, 5335, 5428, 5642 |
| Nepenthes_rowaniae_G05999 | 0.00818 | 10 | 5304, 5348, 6363, 6376, 6378, 6450, 6527, 6601, 6909, 6913 |
| Nepenthes_rowaniae_G08233 | 0.01601 | 3 | 4527, 5280, 5958 |
| Nepenthes_rowaniae_G08234 | 0.01542 | 2 | 5858, 6034 |
| Nepenthes_rowaniae_G08239 | 0.01291 | 5 | 4951, 6034, 6527, 6860, 6913 |
| Nepenthes_rowaniae_G08249 | 0.01417 | 2 | 6034, 6527 |
| Nepenthes_sanguinea_G07858 | 0.01452 | 12 | 5460, 5477, 5770, 5949, 6462, 6531, 6570, 6782, 6792, 6848, 6865, 6962 |
| Nepenthes_sibuyanensis_G08113 | 0.00715 | 10 | 4989, 5123, 5138, 5355, 5936, 5940, 6540, 6570, 7336, 7628 |
| Nepenthes_singalana_G07880 | 0.01101 | 11 | 5434, 5502, 5620, 6407, 6462, 6527, 6667, 6848, 6859, 7028, 7331 |
| Nepenthes_spathulata_G07896 | 0.01025 | 14 | 4802, 5206, 5531, 5894, 6068, 6128, 6216, 6363, 6378, 6383, 6507, 6860, 6955, 7331 |
| Nepenthes_spathulata_G08116 | 0.01226 | 5 | 5206, 5343, 5463, 6068, 7331 |
| Nepenthes_spectabilis_G07890 | 0.0088 | 6 | 5177, 5599, 5639, 5936, 6004, 7324 |
| Nepenthes_stenophylla_G08115 | 0.01107 | 11 | 4691, 5463, 5744, 5772, 5821, 5822, 5842, 6000, 6034, 6299, 6962 |
| Nepenthes_talangensis_G07888 | 0.01312 | 6 | 5502, 5949, 6068, 6216, 6378, 7331 |
| Nepenthes_tenax_G06000 | 0.00668 | 15 | 4691, 5335, 5348, 5366, 5513, 5921, 5922, 6003, 6034, 6458, 6532, 6601, 6860, 6909, 6913 |
| Nepenthes_tenax_G08238 | 0.01259 | 1 |  |
| Nepenthes_tentaculata_G07885 | 0.0152 | 7 | 4989, 5168, 5531, 6003, 6649, 6797, 6924 |
| Nepenthes_thai_G07879 | 0.01213 | 11 | 5123, 5594, 5974, 6072, 6130, 6528, 6532, 6792, 6909, 6962, 7028 |
| Nepenthes_tobaica_G07871 | 0.00987 | 12 | 4802, 5460, 5721, 5822, 5894, 5960, 6000, 6462, 6507, 6914, 7174, 7331 |
| Nepenthes_tobaica_G08122 | 0.00724 | 8 | 4724, 4932, 5123, 5206, 5599, 5770, 5849, 5949 |
| Nepenthes_truncata_G08124 | 0.00588 | 9 | 5188, 5721, 5744, 5843, 6384, 6540, 6713, 6797, 7577 |
| Nepenthes_truncata_x_ventricosa_G08105 | 0.02602 | 4 | 5430, 5770, 6848, 6914 |
| Nepenthes_veitchii_G08088 | 0.0137 | 10 | 4691, 5355, 5849, 5894, 6363, 6462, 6507, 6713, 7141, 7363 |
| Nepenthes_veitchii_G08096 | 0.01618 | 3 | 5469, 5772, 6507 |
| Nepenthes_veitchii_x_eymae_G08108 | 0.01856 | 7 | 4691, 5469, 5620, 6507, 6528, 6532, 6848 |
| Nepenthes_ventricosa_G07860 | 0.00763 | 6 | 5703, 5841, 5940, 6064, 6098, 6198 |
| Nepenthes_ventricosa_G07865 | 0.00872 | 3 | 5940, 6717, 6792 |
| Nepenthes_vogelii_G08110 | 0.01553 | 12 | 4691, 5335, 5469, 5772, 5821, 6004, 6298, 6462, 6848, 6913, 6947, 7135 |
| Nepenthes_xiphioides_G07876 | 0.01376 | 11 | 4724, 5620, 5949, 5960, 6528, 6649, 6738, 6782, 6860, 7577, 7628 |
