## Appendix 9 - Phasing for "HybPhaser: a workflow for the detection and phasing of hybrids in target capture datasets"

Table 1. Accessions to be phased with selected clade references and their abbreviations.

| **Sample** | **Clade reference 1** | **Abbreviation 1** | **Clade reference 2** | **Abbreviation 2** |
| --- | --- | --- | --- | --- |
| bicalcarata x ampullaria | bicalcarata 3 | bica | ampullaria 6 | ampu |
| ampullaria x gracilis | gracilis 1 | graci | ampullaria 6 | ampu |
| boschiana x glandulifera | glandulifera 1 | glan | ventricosa 2 | vent |
| truncata x ventricosa | peltata | pelt | ventricosa 2 | vent |
| burkei x ventricosa 2 | sibuyanensis 2 | sibu | ventricosa 2 | vent |
| burkei x ventricosa 1 | sibuyanensis 2 | sibu | ventricosa 2 | vent |
| burkei x ventricosa 3 | sibuyanensis 2 | sibu | ventricosa 2 | vent |
| kampotiana 1 | mirabilis 25 | mira | suratensis | sura |
| sanguinea 1 | gracillima | grll | suratensis | sura |
| izumiae x ventricosa | ventricosa 2 | vent | jamban | jamb |
| gymnamphora 2 | gymnamphora 3 | gymn | jamban | jamb |
| xiphioides 1 | gymnamphora 3 | gymn | jamban | jamb |
| veitchii x eymae | veitchii 2 | veit | maxima 6 | maxi |
| bellii 1 | mirabilis 25 | mira | merrilliana 2 | merr |
| rafflesiana x ampullaria 1 | ampullaria 6 | ampu | rafflesiana 5 | rafl |
| ampullaria x rafflesiana | ampullaria 6 | ampu | rafflesiana 5 | rafl |
| bokorensis x ventricosa | ventricosa 2 | vent | suratensis | sura |
| sanguinea 2 | gracillima | grll | suratensis | sura |
| angustifolia | gracilis 1 | graci | hispida | hisp |
| rafflesiana x ampullaria 2 | ampullaria 6 | ampu | rafflesiana 5 | rafl |
| rigidifolia | suratensis | sura | spectabilis 1 | spec |
| ampullaria x tobaica | ampullaria 6 | ampu | tobaica 1 | toba |
| clipeata x ventricosa | vogelii 1 | voge | ventricosa 2 | vent |
| veitchii 1 | vogelii 1 | voge | veitchii 2 | veit |
| deaniana | graciliflora 2 | grafl | sp Anipaha | anip |
| lowii x campanulata | faizaliana 1 | faiz | campanulata 2 | camp |

Table 2. Percentages of reads matching unambiguously to each reference (from original read files consisting of on-target and off-target reads), with the exact proportions of the highest/lower value, approximate proportions using whole numbers.

| **Sample name** | **bica** | **ampu** | **graci** | **glan** | **vent** | **pelt** | **sibu** | **mira** | **sura** | **grll** | **jamb** | **gymn** | **veit** | **maxi** | **merr** | **rafl** | **hisp** | **spec** | **toba** | **voge** | **grafl** | **anip** | **faiz** | **camp** | **exact prop.** | **approx. prop.** |
| --- | --- | --- | --- | --- | --- | --- | --- | --- | --- | --- | --- | --- | --- | --- | --- | --- | --- | --- | --- | --- | --- | --- | --- | --- | --- | --- |
| ampullaria x gracilis |  | 3.4 | 4.2 |  |  |  |  |  |  |  |  |  |  |  |  |  |  |  |  |  |  |  |  |  | 1.24 | 1/1 |
| ampullaria x rafflesiana |  | 3.4 |  |  |  |  |  |  |  |  |  |  |  |  |  | 3.1 |  |  |  |  |  |  |  |  | 1.08 | 1/1 |
| ampullaria x tobaica |  | 3.0 |  |  |  |  |  |  |  |  |  |  |  |  |  |  |  |  | 3.2 |  |  |  |  |  | 1.07 | 1/1 |
| angustifolia |  |  | 4.0 |  |  |  |  |  |  |  |  |  |  |  |  |  | 1.3 |  |  |  |  |  |  |  | 3.03 | 3/1 |
| bellii 1 |  |  |  |  |  |  |  | 3.2 |  |  |  |  |  |  | 3.4 |  |  |  |  |  |  |  |  |  | 1.08 | 1/1 |
| bicalcarata x ampullaria | 7.4 | 2.5 |  |  |  |  |  |  |  |  |  |  |  |  |  |  |  |  |  |  |  |  |  |  | 3.01 | 3/1 |
| bokorensis x ventricosa |  |  |  |  | 5.2 |  |  |  | 5.1 |  |  |  |  |  |  |  |  |  |  |  |  |  |  |  | 1.00 | 1/1 |
| boschiana x glandulifera |  |  |  | 3.2 | 1.1 |  |  |  |  |  |  |  |  |  |  |  |  |  |  |  |  |  |  |  | 2.79 | 3/1 |
| burkei x ventricosa 1 |  |  |  |  | 3.7 |  | 5.8 |  |  |  |  |  |  |  |  |  |  |  |  |  |  |  |  |  | 1.55 | 3/2 |
| burkei x ventricosa 2 |  |  |  |  | 3.6 |  | 5.8 |  |  |  |  |  |  |  |  |  |  |  |  |  |  |  |  |  | 1.60 | 3/2 |
| burkei x ventricosa 3 |  |  |  |  | 2.1 |  | 3.2 |  |  |  |  |  |  |  |  |  |  |  |  |  |  |  |  |  | 1.47 | 3/2 |
| clipeata x ventricosa |  |  |  |  | 4.5 |  |  |  |  |  |  |  |  |  |  |  |  |  |  | 4.9 |  |  |  |  | 1.09 | 1/1 |
| deaniana |  |  |  |  |  |  |  |  |  |  |  |  |  |  |  |  |  |  |  |  | 5.0 | 12.3 |  |  | 2.47 | 2/1 |
| gymnamphora 2 |  |  |  |  |  |  |  |  |  |  | 3.8 | 8.8 |  |  |  |  |  |  |  |  |  |  |  |  | 2.34 | 2/1 |
| izumiae x ventricosa |  |  |  |  | 3.8 |  |  |  |  |  | 3.3 |  |  |  |  |  |  |  |  |  |  |  |  |  | 1.15 | 1/1 |
| kampotiana 1 |  |  |  |  |  |  |  | 3.6 | 4.0 |  |  |  |  |  |  |  |  |  |  |  |  |  |  |  | 1.12 | 1/1 |
| lowii x campanulata |  |  |  |  |  |  |  |  |  |  |  |  |  |  |  |  |  |  |  |  |  |  | 2.7 | 6.9 | 2.54 | 3/1 |
| rafflesiana x ampullaria 1 |  | 2.7 |  |  |  |  |  |  |  |  |  |  |  |  |  | 2.4 |  |  |  |  |  |  |  |  | 1.12 | 1/1 |
| rafflesiana x ampullaria 2 |  | 9.0 |  |  |  |  |  |  |  |  |  |  |  |  |  | 9.8 |  |  |  |  |  |  |  |  | 1.10 | 1/1 |
| rigidifolia |  |  |  |  |  |  |  |  | 6.8 |  |  |  |  |  |  |  |  | 8.9 |  |  |  |  |  |  | 1.31 | 1/1 |
| sanguinea 1 |  |  |  |  |  |  |  |  | 4.1 | 6.8 |  |  |  |  |  |  |  |  |  |  |  |  |  |  | 1.63 | 3/2 |
| sanguinea 2 |  |  |  |  |  |  |  |  | 8.6 | 13.9 |  |  |  |  |  |  |  |  |  |  |  |  |  |  | 1.62 | 3/2 |
| truncata x ventricosa |  |  |  |  | 2.7 | 3.2 |  |  |  |  |  |  |  |  |  |  |  |  |  |  |  |  |  |  | 1.17 | 1/1 |
| veitchii 1 |  |  |  |  |  |  |  |  |  |  |  |  | 4.7 |  |  |  |  |  |  | 4.9 |  |  |  |  | 1.06 | 1/1 |
| veitchii x eymae |  |  |  |  |  |  |  |  |  |  |  |  | 5.7 | 4.2 |  |  |  |  |  |  |  |  |  |  | 1.33 | 3/2 |
| xiphioides 1 |  |  |  |  |  |  |  |  |  |  | 4.3 | 8.6 |  |  |  |  |  |  |  |  |  |  |  |  | 1.99 | 2/1 |
