## Appendix 10 - Processing of phased accessions for "HybPhaser: a workflow for the detection and phasing of hybrids in target capture datasets"

Appendix 10 – Assembly of phased accessions – HybPhaser stats

Phased accessions have been

Results from the script ‘Rscript2__dataset_optimization.R’.

**1) Dataset optimisation: Samples and loci removed to reduce missing data**

Graphical overview:

66 loci are below the threshold (0.2) for proportion of recovered samples:

4744, 5034, 5038, 5064, 5177, 5260, 5271, 5328, 5339, 5348, 5354, 5357, 5422, 5513, 5660, 5733, 5802, 5815, 5822, 5843, 5893, 5936, 5941, 5944, 5990, 6026, 6050, 6056, 6114, 6148, 6150, 6164, 6270, 6398, 6401, 6406, 6407, 6430, 6447, 6448, 6457, 6506, 6514, 6526, 6540, 6557, 6565, 6705, 6732, 6780, 6785, 6791, 6864, 6886, 6893, 6954, 6969, 6977, 6995, 7013, 7024, 7135, 7194, 7296, 7325, 7361

11 loci are below the threshold (0.45) for proportion of recovered target sequence length:

| Locus | proportion of recovered target sequence length |
| --- | --- |
| 4793 | 0.384 |
| 5264 | 0.394 |
| 5273 | 0.25 |
| 5333 | 0.338 |
| 5339 | 0.288 |
| 5670 | 0.326 |
| 5816 | 0.386 |
| 5921 | 0.347 |
| 6363 | 0.406 |
| 6544 | 0.341 |
| 7128 | 0.434 |

In total 76 loci were removed: 4744, 5034, 5038, 5064, 5177, 5260, 5271, 5328, 5339, 5348, 5354, 5357, 5422, 5513, 5660, 5733, 5802, 5815, 5822, 5843, 5893, 5936, 5941, 5944, 5990, 6026, 6050, 6056, 6114, 6148, 6150, 6164, 6270, 6398, 6401, 6406, 6407, 6430, 6447, 6448, 6457, 6506, 6514, 6526, 6540, 6557, 6565, 6705, 6732, 6780, 6785, 6791, 6864, 6886, 6893, 6954, 6969, 6977, 6995, 7013, 7024, 7135, 7194, 7296, 7325, 7361, 4793, 5264, 5273, 5333, 5670, 5816, 5921, 6363, 6544, 7128

The average number of missing samples for all loci is: 4.67 (of 25 samples)

The median number of missing samples for all loci is: 2 (of 25 samples)

The chosen threshold for removing loci with missing data is: none

0 loci will be removed.

Cleaning step 1c: missing data (loci) of samples

The average number of missing loci for all samples is: 56.08 (of 300 loci)

The median number of missing loci for all samples is: 51 (of 300 loci)

The chosen threshold for removing samples with missing data is: none.

Cleaning step 1d: Combined sequence length recovered for each sample

The total length of combined target sequences is: 288822 bp. The average proportion of targets length recovered is: 0.94 .The median number of missing loci for all samples is: 0.94 .The chosen threshold for proportion of combined target length to exclude low recovery samples: none .

**Paralogs for all accession**

The 24 putative paralogous accessions from the HybPhaser dataset optimization step for the original dataset were removed:

6412, 6227, 5968, 4471, 6679, 6954, 4757, 5899, 5865, 4806, 6487, 6404, 6563, 6038, 7367, 6274, 5870, 7111, 6883, 5347, 5943, 6373, 4942, 6488.

**Paralogs for each accession**

Table 1. Putative paralogous genes removed for each accession.

| Accession | # of removed loci | name of removed loci |
| --- | --- | --- |
| N_ampullariaXgracilis_G08101_to_ampu | 1 | 5460 |
| N_ampullariaXgracilis_G08101_to_grac | 5 | 6048, 6528, 7067, 7174, 7331 |
| N_ampullariaXtobaica_G08098_to_ampu | 3 | 5090, 5894, 6432 |
| N_ampullariaXtobaica_G08098_to_jamb | 3 | 5469, 6068, 6216 |
| N_ampullariaXtobaica_G08098_to_raja | 5 | 6303, 6420, 6531, 6909, 7331 |
| N_ampullariaXrafflesiana_G08104_to_ampu | 10 | 4992, 5090, 5460, 5894, 6004, 6130, 6265, 6299, 6384, 7273 |
| N_ampullariaXrafflesiana_G08104_to_rafl | 6 | 5772, 6004, 6909, 6924, 7028, 7331 |
| N_bellii_G08102_to_merr | 3 | 4724, 6000, 7029 |
| N_bellii_G08102_to_mira | 4 | 4890, 5531, 6405, 7602 |
| N_bicalcarataXampullaria_G08132_to_ampu | 11 | 4893, 5090, 5296, 5434, 5562, 5594, 6072, 6779, 6859, 7331, 7602 |
| N_bicalcarataXampullaria_G08132_to_bica | 11 | 4890, 4989, 5464, 6072, 6198, 6572, 6962, 6992, 7028, 7174, 7331 |
| N_burkeiXventricosa_G07783_to_sibu | 10 | 5032, 5188, 5469, 5842, 5940, 6068, 6299, 6384, 6660, 7136 |
| N_burkeiXventricosa_G07783_to_vent | 5 | 5032, 5842, 5940, 6068, 6779 |
| N_burkeiXventricosa_G07854_to_sibu | 12 | 5032, 5138, 5551, 5842, 5894, 5940, 5950, 5980, 6068, 6299, 6384, 7333 |
| N_burkeiXventricosa_G07854_to_vent | 7 | 5032, 5449, 5477, 5945, 6649, 6779, 7371 |
| Nepenthes_x_hookeriana_ERR3662102_to_ampu | 6 | 5090, 5477, 5702, 6130, 6299, 7174 |
| Nepenthes_x_hookeriana_ERR3662102_to_rafl | 10 | 5477, 5642, 6048, 6528, 6733, 6782, 6909, 7028, 7174, 7331 |
| N_izumiaeXventricosa_G07866_to_jamb | 5 | 5131, 5304, 5940, 6601, 7279 |
| N_izumiaeXventricosa_G07866_to_merr | 6 | 5032, 5940, 5974, 6299, 6848, 7577 |
| N_rafflesianaXampullaria_G08126_to_ampu | 9 | 5090, 5343, 5620, 6130, 6265, 6299, 6432, 6570, 7028 |
| N_rafflesianaXampullaria_G08126_to_rafl | 11 | 5469, 5477, 5639, 5699, 6110, 6909, 6924, 6992, 7021, 7028, 7331 |
| N_truncataXventricosa_G08105_to_merr | 4 | 5032, 5940, 6383, 6978 |
| N_truncataXventricosa_G08105_to_ceci | 6 | 5940, 6048, 6238, 6383, 6527, 7602 |
| N_veitchiiXeymae_G08108_to_maxi | 8 | 4691, 5339, 5551, 5656, 5770, 6393, 6507, 6528 |
| N_veitchiiXeymae_G08108_to_veit | 14 | 4691, 5335, 5460, 5464, 5620, 5921, 6439, 6458, 6713, 6779, 6792, 6914, 7028, 7141 |

**HybPhaser Step: Heterozygosity assessment**

Table 2. Summary statistics of phased accessions before and after phasing.

| Short name | phased | bp | bp of target | bp clean | bp of target clean | paralogs all | paralogs each | nloci | Locus hetero- zygosity | Allele divergence |
| --- | --- | --- | --- | --- | --- | --- | --- | --- | --- | --- |
| ampullaria x gracilis | no | 441894 | 153 | 400056 | 162.7 | 24 | 8 | 275 | 97.1% | 1.22% |
| ampullaria x rafflesiana | no | 454423 | 157.3 | 417129 | 169.6 | 24 | 12 | 277 | 95.3% | 1.05% |
| ampullaria x tobaica | no | 314306 | 108.8 | 289633 | 117.8 | 24 | 7 | 259 | 93.8% | 1.13% |
| angustifolia | no | 291298 | 100.9 | 270242 | 109.9 | 24 | 3 | 151 | 97.4% | 1.51% |
| bellii 1 | no | 317247 | 109.8 | 289177 | 117.6 | 24 | 7 | 257 | 91.4% | 1.21% |
| bicalcarata x ampullaria | no | 259379 | 89.8 | 236810 | 96.3 | 24 | 8 | 260 | 84.2% | 0.79% |
| bokorensis x ventricosa | no | 139944 | 48.5 | 127329 | 51.8 | 24 | 2 | 98 | 93.9% | 1.12% |
| boschiana x glandulifera | no | 145871 | 50.5 | 130469 | 53 | 24 | 8 | 207 | 41.6% | 0.28% |
| burkei x ventricosa 1 | no | 416431 | 144.2 | 378174 | 153.8 | 24 | 13 | 278 | 83.5% | 0.36% |
| burkei x ventricosa 2 | no | 399549 | 138.3 | 363682 | 147.9 | 24 | 13 | 273 | 85.7% | 0.36% |
| burkei x ventricosa 3 | no | 361999 | 125.3 | 328941 | 133.7 | 24 | 20 | 265 | 81.5% | 0.32% |
| clipeata x ventricosa | no | 468685 | 162.3 | 424278 | 172.5 | 24 | 6 | 274 | 95.6% | 1.16% |
| deaniana | no | 324868 | 112.5 | 301591 | 122.6 | 24 | 11 | 153 | 81.1% | 0.39% |
| gymnamphora 2 | no | 144730 | 50.1 | 136463 | 55.5 | 24 | 0 | 78 | 94.9% | 0.42% |
| izumiae x ventricosa | no | 394144 | 136.5 | 359713 | 146.2 | 24 | 10 | 267 | 95.1% | 1.08% |
| kampotiana 1 | no | 440586 | 152.5 | 399435 | 162.4 | 24 | 11 | 284 | 95.8% | 1.27% |
| lowii x campanulata | no | 451830 | 156.4 | 410562 | 166.9 | 24 | 13 | 282 | 94.3% | 0.87% |
| rafflesiana x ampullaria 2 | no | 584706 | 202.4 | 535631 | 217.8 | 24 | 12 | 277 | 94.6% | 1.24% |
| rafflesiana x ampullaria 1 | no | 424810 | 147.1 | 386986 | 157.3 | 24 | 7 | 280 | 95.7% | 0.98% |
| rigidifolia | no | 501658 | 173.7 | 456628 | 185.7 | 24 | 15 | 268 | 89.2% | 0.47% |
| sanguinea 2 | no | 582558 | 201.7 | 527250 | 214.4 | 24 | 12 | 268 | 82.5% | 0.41% |
| sanguinea 1 | no | 408434 | 141.4 | 370059 | 150.5 | 24 | 12 | 274 | 83.9% | 0.44% |
| truncata x ventricosa | no | 310700 | 107.6 | 286291 | 116.4 | 24 | 4 | 258 | 92.6% | 1.07% |
| veitchii 1 | no | 419359 | 145.2 | 382266 | 155.4 | 24 | 10 | 277 | 87.0% | 0.39% |
| veitchii x eymae | no | 534053 | 184.9 | 487548 | 198.2 | 24 | 7 | 280 | 92.1% | 0.56% |
| xiphioides 1 | no | 483298 | 167.3 | 436980 | 177.7 | 24 | 11 | 275 | 89.8% | 0.50% |
| ampullaria x gracilis to ampu | yes | 317219 | 109.8 | 286055 | 137.3 | 24 | 2 | 227 | 37.4% | 0.22% |
| ampullaria x gracilis to graci | yes | 341075 | 118.1 | 302242 | 145 | 24 | 7 | 236 | 55.5% | 0.39% |
| ampullaria x rafflesiana to ampu | yes | 332348 | 115.1 | 300698 | 144.3 | 24 | 10 | 233 | 76.4% | 0.49% |
| ampullaria x rafflesiana to rafl | yes | 318397 | 110.2 | 286496 | 137.5 | 24 | 7 | 223 | 58.3% | 0.42% |
| ampullaria x tobaica to ampu | yes | 234212 | 81.1 | 212290 | 101.9 | 24 | 4 | 216 | 40.7% | 0.19% |
| ampullaria x tobaica to toba | yes | 236692 | 82 | 210983 | 101.2 | 24 | 6 | 209 | 43.1% | 0.31% |
| angustifolia to graci | yes | 456125 | 157.9 | 400357 | 192.1 | 24 | 7 | 242 | 89.7% | 1.00% |
| angustifolia to hisp | yes | 143847 | 49.8 | 131617 | 63.2 | 24 | 1 | 78 | 55.1% | 0.32% |
| bellii 1 to merr | yes | 232358 | 80.5 | 206768 | 99.2 | 24 | 2 | 208 | 44.2% | 0.31% |
| bellii 1 to mira | yes | 233855 | 81 | 212580 | 102 | 24 | 2 | 214 | 38.8% | 0.16% |
| bicalcarata x ampullaria to ampu | yes | 155681 | 53.9 | 139554 | 67 | 24 | 11 | 200 | 52.5% | 0.23% |
| bicalcarata x ampullaria to bica | yes | 166449 | 57.6 | 144803 | 69.5 | 24 | 11 | 191 | 61.8% | 0.34% |
| bokorensis x ventricosa to sura | yes | 267788 | 92.7 | 239237 | 114.8 | 24 | 7 | 221 | 52.9% | 0.17% |
| bokorensis x ventricosa to vent | yes | 283542 | 98.2 | 251476 | 120.7 | 24 | 3 | 233 | 65.7% | 0.32% |
| boschiana x glandulifera to glan | yes | 102601 | 35.5 | 93771 | 34.9 | 24 | 4 | 173 | 34.1% | 0.17% |
| boschiana x glandulifera to vent | yes | 30734 | 10.6 | 24389 | 9.1 | 24 | 1 | 66 | 15.2% | 0.14% |
| burkei x ventricosa 1 to sibu | yes | 285489 | 98.8 | 259083 | 124.3 | 24 | 10 | 224 | 63.8% | 0.17% |
| burkei x ventricosa 1 to vent | yes | 234530 | 81.2 | 203665 | 97.7 | 24 | 6 | 214 | 57.5% | 0.21% |
| burkei x ventricosa 2 to sibu | yes | 271156 | 93.9 | 244826 | 117.5 | 24 | 12 | 223 | 65.5% | 0.17% |
| burkei x ventricosa 2 to vent | yes | 216889 | 75.1 | 186209 | 89.4 | 24 | 7 | 208 | 59.1% | 0.21% |
| burkei x ventricosa 3 to sibu | yes | 223393 | 77.3 | 202710 | 97.3 | 24 | 10 | 213 | 66.2% | 0.19% |
| burkei x ventricosa 3 to vent | yes | 179313 | 62.1 | 157045 | 75.4 | 24 | 8 | 193 | 59.1% | 0.20% |
| clipeata x ventricosa to vent | yes | 334173 | 115.7 | 296801 | 142.4 | 24 | 3 | 233 | 57.5% | 0.32% |
| clipeata x ventricosa to voge | yes | 378277 | 131 | 334223 | 160.4 | 24 | 8 | 239 | 54.0% | 0.21% |
| deaniana to anip | yes | 227131 | 78.6 | 196016 | 94.1 | 24 | 7 | 112 | 79.5% | 0.36% |
| deaniana to grafl | yes | 277445 | 96.1 | 246220 | 118.1 | 24 | 5 | 193 | 54.9% | 0.21% |
| gymnamphora 2 to gymn | yes | 384390 | 133.1 | 345136 | 165.6 | 24 | 7 | 234 | 77.8% | 0.27% |
| gymnamphora 2 to jamb | yes | 253561 | 87.8 | 218770 | 105 | 24 | 11 | 201 | 34.8% | 0.10% |
| izumiae x ventricosa to jamb | yes | 295873 | 102.4 | 265710 | 127.5 | 24 | 5 | 225 | 25.3% | 0.10% |
| izumiae x ventricosa to vent | yes | 296583 | 102.7 | 264430 | 126.9 | 24 | 6 | 231 | 42.9% | 0.17% |
| kampotiana 1 to mira | yes | 322598 | 111.7 | 292553 | 140.4 | 24 | 4 | 233 | 51.5% | 0.29% |
| kampotiana 1 to sura | yes | 323938 | 112.2 | 286684 | 137.6 | 24 | 5 | 229 | 38.0% | 0.21% |
| lowii x campanulata to camp | yes | 368558 | 127.6 | 327968 | 157.4 | 24 | 9 | 241 | 88.0% | 0.48% |
| lowii x campanulata to faiz | yes | 259916 | 90 | 226813 | 108.8 | 24 | 5 | 223 | 31.4% | 0.13% |
| rafflesiana x ampullaria 2 to ampu | yes | 436932 | 151.3 | 391650 | 187.9 | 24 | 6 | 233 | 57.1% | 0.40% |
| rafflesiana x ampullaria 2 to rafl | yes | 450911 | 156.1 | 401449 | 192.6 | 24 | 10 | 223 | 65.5% | 0.52% |
| rafflesiana x ampullaria 1 to ampu | yes | 279573 | 96.8 | 252545 | 121.2 | 24 | 9 | 231 | 78.4% | 0.45% |
| rafflesiana x ampullaria 1 to rafl | yes | 257648 | 89.2 | 228062 | 109.4 | 24 | 12 | 216 | 69.0% | 0.40% |
| rigidifolia to spec | yes | 365741 | 126.6 | 323963 | 155.5 | 24 | 13 | 233 | 79.8% | 0.38% |
| rigidifolia to sura | yes | 230881 | 79.9 | 205167 | 98.4 | 24 | 6 | 169 | 56.8% | 0.19% |
| sanguinea 2 to grll | yes | 435859 | 150.9 | 390890 | 187.6 | 24 | 9 | 229 | 76.4% | 0.35% |
| sanguinea 2 to sura | yes | 281347 | 97.4 | 245482 | 117.8 | 24 | 6 | 177 | 65.5% | 0.27% |
| sanguinea 1 to grll | yes | 302323 | 104.7 | 272679 | 130.8 | 24 | 9 | 232 | 78.5% | 0.36% |
| sanguinea 1 to sura | yes | 173626 | 60.1 | 147197 | 70.6 | 24 | 6 | 177 | 61.6% | 0.33% |
| truncata x ventricosa to pelt | yes | 224000 | 77.6 | 199367 | 95.7 | 24 | 1 | 210 | 27.1% | 0.10% |
| truncata x ventricosa to vent | yes | 223210 | 77.3 | 197809 | 94.9 | 24 | 4 | 212 | 31.6% | 0.12% |
| veitchii 1 to veit | yes | 254492 | 88.1 | 224300 | 107.6 | 24 | 6 | 212 | 70.3% | 0.29% |
| veitchii 1 to voge | yes | 269087 | 93.2 | 238511 | 114.4 | 24 | 5 | 223 | 68.2% | 0.25% |
| veitchii x eymae to maxi | yes | 321083 | 111.2 | 291582 | 139.9 | 24 | 7 | 224 | 49.6% | 0.19% |
| veitchii x eymae to veit | yes | 376664 | 130.4 | 334723 | 160.6 | 24 | 12 | 239 | 69.5% | 0.24% |
| xiphioides 1 to gymn | yes | 377272 | 130.6 | 339162 | 162.7 | 24 | 11 | 235 | 76.2% | 0.27% |
| xiphioides 1 to jamb | yes | 243540 | 84.3 | 208636 | 100.1 | 24 | 4 | 200 | 30.0% | 0.11% |
